# Ancient whole genomes reveal regional selection during adaptation to Neolithic lifestyle in Western Eurasia

**DOI:** 10.64898/2026.07.15.738373

**Authors:** Miquel Àngel Schikora-Tamarit, Pablo Carrión, Simon Orozco-Arias, Aida Andrades Valtueña, Ana Ivanović, Christiana Lyn Scheib, Helja Kabral, Ivo Glynne Gut, Kristiina Tambets, Marta Gut, Marko Porčić, Sofija Stefanović, Vladimir Zhurov, Lepenski Vir consortium, Carles Lalueza-Fox, Miodrag Grbic, Toni Gabaldón

**Affiliations:** Barcelona Supercomputing Centre (BSC-CNS). Plaça Eusebi Güell, 1-3, 08034 Barcelona, Spain; Institute for Research in Biomedicine (IRB Barcelona), The Barcelona Institute of Science and Technology, Baldiri Reixac, 10, 08028 Barcelona, Spain; CIBER de Enfermedades Infecciosas, Instituto de Salud Carlos III, Madrid, Spain; Department of Archaeogenetics, Max Planck Institute for Evolutionary Anthropology, Leipzig, Germany; Institut de Biologia Evolutiva (CSIC - UPF), Barcelona, Spain; Max Planck-Harvard Research Center for the Archaeoscience of the Ancient Mediterranean, Leipzig, Germany; University of Belgrade, Faculty of Biology, Belgrade, Serbia; Estonian Biocentre, Institute of Genomics, University of Tartu, Tartu, Estonia; Centro Nacional de Análisis Genómico, Barcelona, Spain; Universitat de Barcelona, Barcelona, Spain; Laboratory for Bioarchaeology, Department of Archaeology, Faculty of Philosophy, University of Belgrade.; University of Western Ontario, Department of Biology, London, Canada; Natural Sciences Museum of Barcelona (MCNB), Barcelona, Spain; Department of Agriculture and Food Science, University of La Rioja, Logroño, Spain; University of Montenegro, Center for Interdisciplinary and Multidisciplinary Studies, Podgorica, Montenegro; Catalan Institution for Research and Advanced Studies (ICREA), Barcelona, Spain

**Author notes:** These authors share senior authorship and correspondence. Equal contribution (Miquel Àngel Schikora-Tamarit, Pablo Carrión). see authors and affiliations in **Consortia Information**.

## Abstract

The introduction of agriculture in Western Eurasia during the Mesolithic-Neolithic transition reshaped human lifestyle, demography and pathogen exposure. Yet, the associated selective pressures remain incompletely resolved because previous ancient DNA studies relied on targeted SNP panels and yielded non-overlapping results. Here, using high-coverage whole-genome data from 152 ancient individuals, including 12 newly sequenced genomes from the Iron Gates, we show that adaptation during this transition was geographically structured and involved selection of both Early Farmer and Hunter-Gatherer alleles. By combining scans of population differentiation, extended haplotype homozygosity, and adaptive admixture, we identify candidate loci under selection in ancestral and admixed populations. Extending the analysis beyond SNPs, we also detect differential copy-number variants, ancestry-associated shifts in transposable-element abundance, and an early Neolithic hepatitis B infection. Functional analyses including effects on gene expression suggested selection on immune, metabolic, reproductive, sensory, and behavioural processes. Taken together, our results reveal a broader and more regionally variable adaptive landscape than previously appreciated.

## INTRODUCTION

The adoption of agriculture was a major technological and cultural revolution that transformed human societies during the Mesolithic-Neolithic transition. In Western Eurasia, farming emerged in the Fertile Crescent ∼10,000 years ago and spread into Europe over the subsequent millennia^1^. The study of ancient DNA (aDNA) from archaeological sites has shown that this cultural spread was accompanied by large-scale migration of Early Farmers (EF) and their admixture with local European Mesolithic Hunter-Gatherers (HG), resulting in late Neolithic populations with varying degrees of each of these divergent genetic ancestries^2–4^. The Mesolithic-Neolithic transition involved profound changes in lifestyle and demography, likely altering selective pressures on human populations. While Mesolithic period populations were nomadic or semi-sedentary and had relatively sparse settlements, Neolithic period populations had increased densities^5^ and closer contact with domesticated plants and animals, which likely exposed them to new pathogens and exerted selection on immune system genes^6–10^. Additionally, a shift towards less diverse, starch-rich diets may have impacted metabolism-related genes^6,8,10,11^. Finally, it has been proposed that the higher complexity of Neolithic societies may have shaped sociability traits^6,8,10^. Consistent with these expectations, recent aDNA studies have reported signals of selection at loci related to pigmentation, lactose intolerance, lipid metabolism, and immune response genes^6,12–14^. Some of these studies identified differential selective pressures suffered by ancestral EF/HG populations^12,14^, while others analyzed the signs of adaptive admixture in admixed late Neolithic populations ^6,12,13^.

Despite this progress, several important gaps remain. First, most studies relied on panels of single-nucleotide polymorphisms (SNPs) in coding regions, which cannot fully resolve selective pressures on non-targeted regions, and generally preclude the analysis of copy-number variants (CNVs) and transposable elements (TEs), which may have contributed substantially to adaptation^15–18^. Similarly, panel-based data do not allow the study of non-human DNA present in ancient samples, including pathogens that may have shaped selective processes^6,7,9,19^. Second, inference of biological processes under selection has mostly relied on the analysis of (generally a few) genes near SNPs of interest, rather than on systematic examination of functional enrichments and associations with gene regulation or phenotypic changes. Third, different studies yielded largely non-overlapping results, requiring further efforts to clarify the relevant selective pressures. This may be due to the use of different analytical frameworks, such as the study of differential selective pressures between ancestral populations^12,14^ vs adaptive admixture^6,12,13^, or the analysis of SNPs^6,12–14^ vs haplotypes^12,13^, which interrogate selection at different timescales and therefore provide different results that are difficult to reconcile. Fourth, most adaptive-admixture studies treated late Neolithic admixed individuals as part of a single, homogeneous population with fixed ancestry proportions, defined on chronological rather than genetic grounds^6,13^. This may bias the results due to i) the inclusion of individuals with more complex demographies, and ii) the limited resolution to detect local adaptation in different geographies, which may be relevant ^20,21^.

Here, we addressed these gaps using high-coverage aDNA whole-genome sequencing (WGS) data, integrating distinct selection-detection methods, and providing a comprehensive functional analysis of variant effects. We assembled a dataset of 152 individuals spanning EF, HG, and late Neolithic admixed groups, including 12 newly generated genomes that increase the representation of HG individuals. We combined complementary selection scans capturing differential selective pressures between ancestral populations and adaptive admixture, at both SNP and haplotype levels. In addition to SNP-based signals, we identify differentiated CNVs and ancestry-associated shifts in TE family abundance, uncovering a previously-neglected layer of genomic change across the Mesolithic–Neolithic transition. Additionally, we inferred biological processes under selection using predicted regulatory effects and variant-phenotype associations, suggesting strong immune-related signals alongside impacts on metabolic and reproductive processes. Importantly, our results reveal a previously underappreciated contribution of HG-derived alleles to adaptive processes in late Neolithic populations.

## RESULTS AND DISCUSSION

### A comprehensive dataset for inferring selection during the Mesolithic-Neolithic transition

To investigate selective pressures during the Mesolithic-Neolithic transition, we first compiled publicly available WGS individuals from the Allen Ancient DNA Resource (AADR)^22^ that fulfilled a set of inclusion criteria. In brief, we selected data from Western Eurasian individuals with low contamination rates, no close kinship relationship, coverage > 0.5x, and assignment to the EF, HG, or EF+HG admixed populations according to qpAdm ancestry modeling (see **Methods**, **Fig. 1a, Extended Data Fig. 1,2**). This resulted in 48 EF, 11 HG and 82 admixed individuals, providing a relevant dataset for inferring Mesolithic-Neolithic selection. However, the small number of HG individuals (11) limited our ability to accurately model the contribution of HG alleles, prompting us to generate additional high-coverage WGS data.

**Fig. 1.**
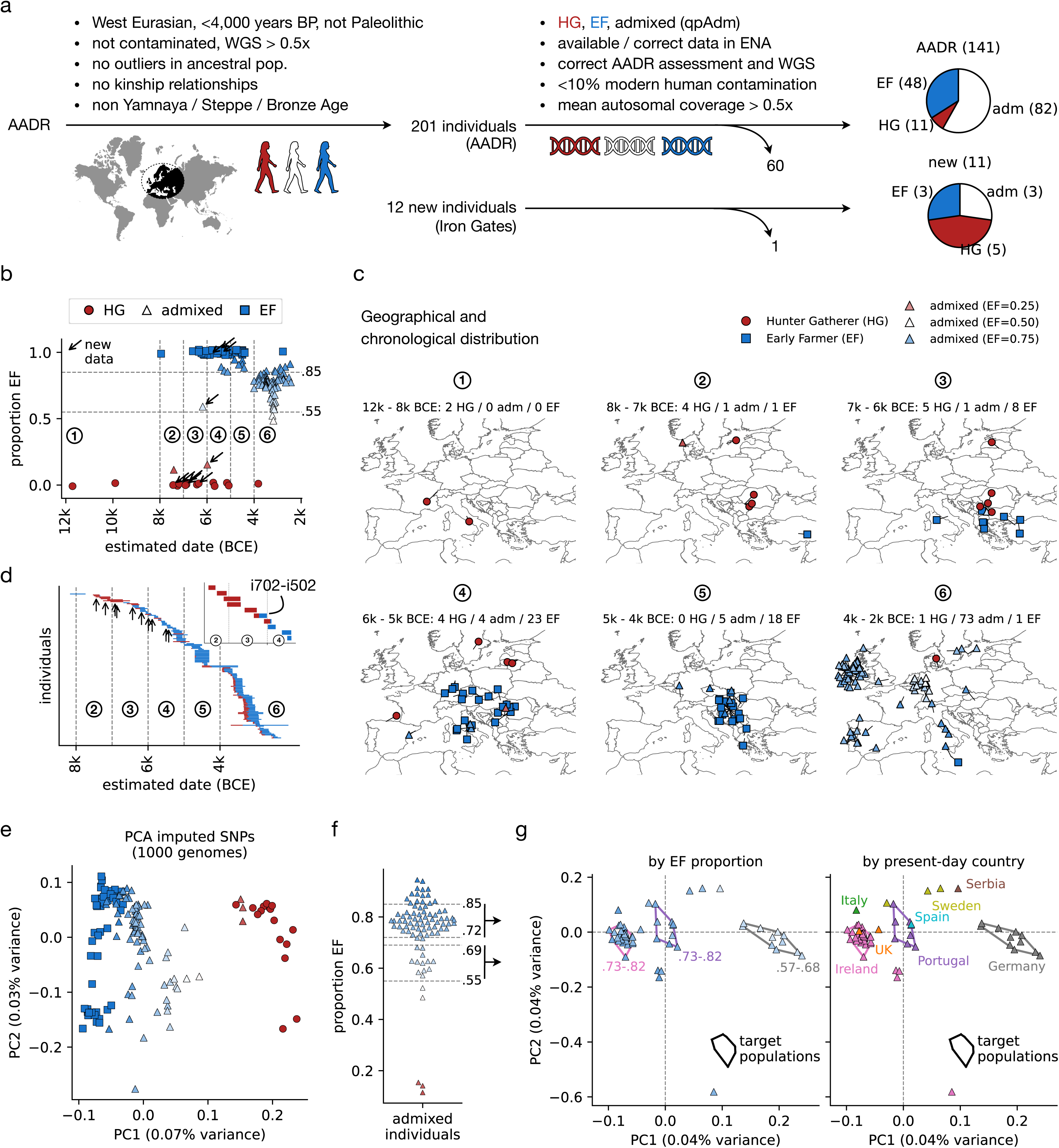
Overview of the samples analyzed here. **a.** To study selection during the Mesolithic-Neolithic transition we assembled high-coverage WGS, carefully curated datasets for Western-Eurasian individuals. We include all publicly available datasets from the Allen Ancient DNA Resource (AADR, top)^22^, as well as 11 newly-sequenced ones (bottom). The pie charts represent the numbers of individuals with an Early Farmer (EF), Hunter Gatherer (HG) or admixed (adm) ancestry in each of the collections. See **Methods** for details regarding the filtering steps (bullet points). **b.** Chronological distribution of the analyzed individuals, where the y axis and symbols and colors represent the ancestry (EF, HG, admixed). Note that the pure HG / EF samples have a little jittering along the y axis to enable better visualization. Also, the arrows indicate the newly-sequenced individuals, while the small numbers point to temporal bins used for subsequent geographical visualization. Individual i704−i504 does not appear in the plot as it lacked RC dating. **c.** Geographical origin of the studied individuals along the six temporal bins from **b**. **d.** Timeplot and dating (Radio Carbon (RC) calibrated) of the studied individuals. Each bar represents the ancestral proportions of their modeled ancestry (see **Methods** and **Supplementary Results**), and the small inset represents the newly sequenced individuals with the early admixed i702-i502 highlighted.. Also, note that the two HG individuals from <8,000 BCE (see **b**) are not included to enhance resolution. **e.** PCA based on imputed SNPs for all autosomes for our whole dataset, with the colors and symbols equivalent to **b**. **f.** Distribution of inferred EF ancestry proportion for the admixed individuals. To identify target admixed sub-populations for adaptive admixture scans we focused on those within the major distribution peaks (0.55–0.69 and 0.72–0.85), outlined with the arrows. **g.** PCA based on imputed SNPs for individuals with 0.55–0.69 or 0.72–0.85 EF proportions, colored by EF proportion (left) or geographical origin as indicated by the present-day countries of isolation (right). We define as target sub-populations for adaptive admixture scans those groups of individuals with a similar EF proportion, geographical origin, and PCA coordinates, as outlined with the polygon lines. Throughout the text, we call these sub-populations nGer, nIre and nPor, matching the present-day countries of isolation (Germany, Ireland, Portugal).

We therefore sequenced the genomes of 12 new individuals from the Lepenski Vir archaeological site (present-day Serbia) within the Iron Gates region. Situated along the Danube River, this area offers a unique perspective on the Mesolithic-Neolithic interface, as its Mesolithic culture was characterized by complex HG societies, dense settlements, rich material culture, and the first appearance of monumental sculptures in Europe^23^. We assessed post-mortem damage patterns and screened for modern contamination using mtDNA haplogroup information in the obtained sequences. All samples were compatible with ancient authenticity (see **Methods**, **Supplementary Table 1**) and low contamination (<3.3% for all samples). Applying the same inclusion criteria as for the AADR individuals, we selected 3 EF, 5 HG and 3 admixed individuals (**Fig. 1a**). While the number of newly sequenced individuals is modest, we significantly increase the number of HG (11 to 16), substantially improving our ability to analyse the contribution of HG alleles to selective processes. In total, our final dataset comprises 51 EF, 16 HG and 85 admixed individuals (**Supplementary Table 1**), providing a valuable resource to study selection during the Mesolithic-Neolithic transition.

### Analysis of individual ancestries reveals incipient population admixture in the Iron Gates

To obtain a more detailed view on the demographic history of the analyzed individuals, we examined qpAdm-inferred ancestries (see **Methods**) in the context of each individual’s chronology and geographic origin. Individuals with mixed HG and EF ancestry begin to appear in our dataset during the early–mid 5th millennium BCE, and largely replace genetically unadmixed HG and EF individuals by the late 5th to early 4th millennium BCE across the studied regions (**Fig. 1 b**). These admixed individuals are distributed across much of Western Eurasia (**Fig. 1 c**), consistent with previous aDNA studies showing that the Mesolithic-Neolithic transition was not a single, uniform event, but a prolonged process marked by regionally variable admixture between incoming EF and local HG^24,25^. Overall, these results support a demographic transition that unfolded progressively over the 5th millennium BCE.

One notable exception to this trend is a newly sequenced admixed individual from Lepenski Vir, i702–i502, who carries 59% EF ancestry yet dates before ∼6000 BCE. This finding suggests that the Iron Gates region was an unusually early zone of interaction and admixture between HG and EF populations. To validate this and further characterize the newly sequenced individuals we projected their genetic variability into present-day Western Eurasian diversity using Principal Component Analysis (PCA, see **Methods**). This revealed three clearly separated clusters along PC1 (**Extended Data Fig. 3a**). Eight individuals had lower PC1 values, overlapping with previously reported Mesolithic individuals from the Iron Gates^26,27^. By contrast, three individuals yielded positive PC1 values (**Extended Data Fig. 3a**), similar to previously reported Iron Gates Neolithic individuals^26,28,29^. We therefore classified these two clusters as Western Hunter-Gatherers (WHG, a subgroup of HG) and Early European Farmers (EEF, a subgroup of EF), respectively. The remaining individual, i702−i502, occupied an intermediate position between the WHG and EEF clusters, further supporting its admixed ancestry.

We next examined qpAdm ancestry models for the newly sequenced individuals in more detail (see **Methods, Supplementary Table 1**). The three-individual cluster with positive PC1 values was best modelled with Balkan EEF ancestry (**Extended Data Fig. 3a**). The intermediate i702−i502 individual could be modeled as a near 50-50 mix of EEF and WHG ancestries, confirming its recent admixed origin (**Fig. 1d**). For the larger cluster, six individuals formed a genetically homogeneous cluster with 100% ancestry from Iron Gates WHG populations, while the remaining two were best modelled combining WHG and EEF ancestries, with the latter accounting for 15.6 +/- 2.5% (i703-i503) to 9.3 +/- 2.8% (i704-i504). Together, these findings indicate close interactions between the local Balkan WHG and incoming EEF populations during the Neolithic transition, and confirm the Iron Gates region as an unexpectedly early contact zone between these ancestral populations.

### Signatures of selection during the Mesolithic-Neolithic transition

To identify loci affected by selection during the Mesolithic-Neolithic transition from aDNA shotgun data, we first benchmarked the performance of various variant-calling software and filtering parameters when applied to simulated aDNA sequencing reads matching the properties of each of our individuals (see **Methods**, **Extended Data Fig. 4,5,6**). Although this benchmarking provides a useful framework for future variant-calling efforts in ancient genomes, we could not find a combination of settings achieving acceptable performance for accurate downstream analysis (**Supplementary Information, Extended Data Fig. 7,8,9**). We therefore adopted a SNP imputation strategy based on mapped reads and the 1000 Genomes reference panel^30^, which has been validated for similar aDNA datasets^31^. This yielded 6,008,635 SNPs with MAF>=0.05 across all individuals of HG/EF groups, and was sufficient information to clearly differentiate the HG, EF and admixed populations in a PCA (**Fig. 1 e**).

To enable accurate adaptive admixture scans, we tested whether admixed individuals could reasonably be treated as a single late Neolithic population with fixed average ancestry proportions, as assumed in previous studies^6,13^. In contrast to this assumption, EF ancestry proportions showed a broad distribution, with most individuals falling into two peaks centered at 0.55–0.69 and 0.72–0.85 (**Fig. 1f**), suggesting that previous assumptions could yield misleading results. To define more homogeneous subpopulations for adaptive admixture scans, we jointly considered EF ancestry proportion, geographical origin, and coordinates in a SNP-based PCA. This identified three subgroups, including certain Neolithic period individuals from present-day Germany (nGer, n=11), Ireland (nIre, n=32) and Portugal (nPor, n=8), each characterized by homogeneous EF proportions (range < 12%), the same present-day country of origin, and specific PCA clustering (**Fig. 1g**). We therefore used these three subgroups as target populations for adaptive admixture scans, enabling the analysis of geographically constrained adaptation during the Mesolithic-Neolithic transition.

To infer differential selective pressures between ancestral HG/EF populations, we used two complementary approaches. First, we used PCAdapt^14,32^ to detect imputed SNPs driving the HG/EF separation along PC1, resulting in 8 non-redundant significant SNP hits (p < 4.42·10^−77^) (**Fig. 2a-d, 3, Supplementary Table 2**). Second, we performed a haplotype-based cross-population Extended Haplotype Homozygosity (XP-EHH) scan^12,33^ to identify recent selective sweeps, yielding 12 and 8 hits non-redundant hits (Bonferroni-corrected p<0.05) in EF and HG populations, respectively, indicating selection on both ancestral backgrounds (**Fig. 2e-g, 3**).

**Fig. 2.**
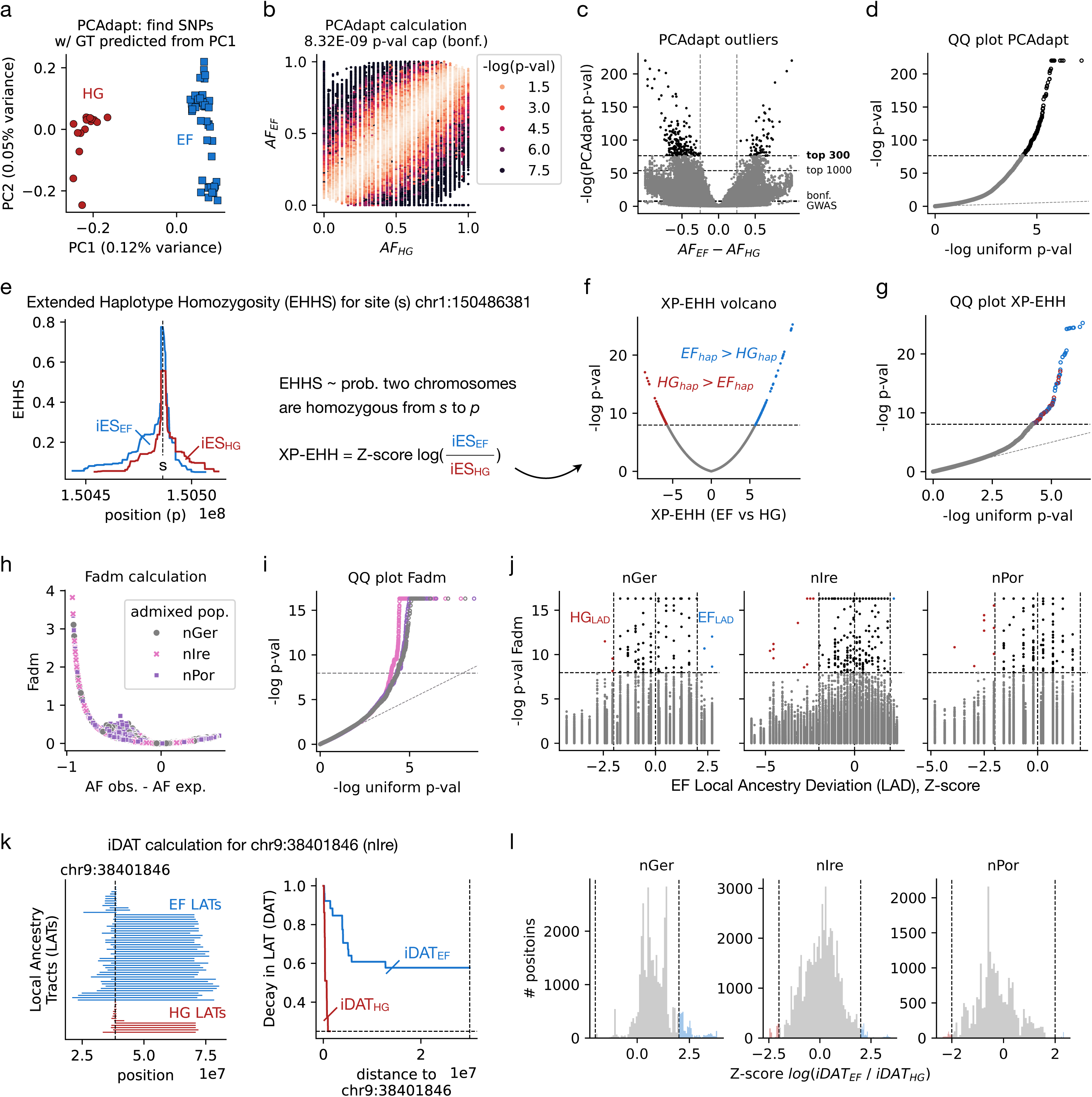
Integrating various methods to detect SNPs under selection. **a.** Principal Component Analysis (PCA) based on all imputed SNPs for HG and EF individuals. Given the separation of the two ancestral populations by PC1, we performed a component-wise PCAdapt scan to find SNPs whose genotype in a given individual (GT) is strongly predictable from PC1, indicating selection. **b.** For each SNP, allele frequency (AF) in each of the HG / EF populations (x and y-axis, respectively) vs the -log p-value as calculated by the PCAdapt PC1 scan. **c.** Difference in AF between the ancestral populations (x-axis) vs the -log PCAdapt p-value, with different horizontal lines reflecting various p-value thresholds defining ‘outlier’ significant SNPs (GWAS standard 5·10^−8^, Bonferroni, top 1000 and top 300 lowest). The black points reflect SNPs passing the used threshold (top 300 lowest p-value). **d.** Quantile-Quantile (QQ) plot for uniformly-distributed vs observed -log PCAdapt p-values, where black points indicate SNPs passing the significance threshold. **e.** Calculation of cross-population Extended Haplotype Homozygosity (XP-EHH) scores for an example site *s* (chr1:150486381). For each position *p* surrounding *s*, we calculate the site-specific Extended Haplotype Homozygosity (EHHS) as the probability that two chromosomes in a given population (EF or HG) are homozygous from *s* to *p*, and then we define iES as the area under the *p* vs EHHS curve. Finally, XP-EHH is calculated as the Z-score (across all sites) of the natural logarithm of the ratio between the iES of each population. **f.** Volcano plot of the XP-EHH metrics across the genome and their corresponding -log p-values. In red and blue we show SNPs passing the Bonferroni correction, reflecting sites under selection in HG or EF, respectively. **g.** QQ-plot of the XP-EHH p-values, as in **d**. **h.** Fadm statistics calculation for all SNPs in each of the admixed populations (nGer, nIre and nPor). For each SNP, we calculated the AF observed or expected from a mixture model based on the genome-wide EF and HG ancestry proportions (AF difference in the x-axis), which are used to infer Fadm values (y-axis). **i.** QQ-plot of the p-values associated to the Fadm metrics, as in **d**, where the horizontal line reflects the Bonferroni-correction. **j.** EF Local Ancestry Deviation, calculated as the Z-score of the fraction of EF Local Ancestry Tracts (LATs) for a given site, vs the -log Fadm p-value for each SNP in each admixed population; used to provide directionality to the Fadm scan. The horizontal lines reflect the Bonferroni threshold for Fadm p-values, and the vertical ones indicate the +-2 EF LAD threshold used to define sites with a significant enrichment of the EF or HG ancestry, respectively. **k.** Calculation of iDAT scores for an example site of chromosome 9 in the nIre admixed population. The left panel shows the span of each of the EF and HG LATs, while the right panel shows the Decay in LAT fraction (DAT) for positions increasingly distant to the site. We calculated iDAT_HG_ and iDAT_EF_ as the area under the distance vs DAT curve for each of the HG and EF LATs, respectively. **l.** Distribution of the iDAT ratio for each of the admixed populations, calculated as the Z-score (across all sites) of the natural logarithm of iDAT_EF_ / iDAT_HG_. The vertical lines indicate the +-2 thresholds used to identify sites with particularly high or low iDAT ratio scores, indicating selection of EF or HG alleles, respectively.

To infer signs of adaptive admixture in each of the three regional admixed populations, we again used two approaches. First, we calculated Fadm statistics reflecting SNPs with unexpected AF given the HG+EF mixture model^13,24^, and measured EF Local Ancestry Deviation (EF LAD) to infer whether AF changes reflect selection of EF or HG alleles^13,34^ (**Fig. 2h-j, 3, Extended Data Fig. 10**). We kept 27 non-redundant hits (Bonferroni-corrected p<0.05, 12 from nPor, 9 from nIre, 6 from nGer), which mostly (26/27) lack clear directionality (EF LAD between -2 and 2), suggesting that most AF deviations cannot be attributed to a specific ancestry. Second, to find adaptive admixture specifically favouring EF or HG alleles, we calculated haplotype-based iDAT (integrated Decay in Ancestry Tracts) scores^35^, resulting in 30 non-redundant hits (iDAT ≥ 2 or iDAT ≤ -2), 10 with highest absolute iDAT value each population (**Fig. 2k,l, 3, Extended Data Fig. 11**). Of these, 19 favour EF ancestry and 11 favour HG ancestry, again indicating contributions from both backgrounds to post-admixture adaptation. See **Supplementary Information** for further details.

These scans identified 85 hits, which we tried to prioritize using two strategies. First, since different methods may capture equivalent selective processes (e.g. a given selective sweep in EF captured by PCAdapt and XP-EHH), we tried to prioritize SNPs supported by different scans. However, only 9/85 hits are shared across methods and reflect equivalent sweeps (**Extended Data Fig. 12,13, Supplementary Information**), suggesting that different approaches capture complementary selection acting at different evolutionary timescales and geographical regions. Second, we compared our hits to six loci highlighted in previous studies^6,12–14^, but we could only find overlaps for *FADS1* and *HLA* (**Extended Data Fig. 14,15, Supplementary Table 3, Supplementary Information**). This may result from differences in methodologies and studied populations, consistent with the similarly low overlap across previous studies (**Extended Data Fig. 14,15**). Hence, we considered that overlap across scans and/or studies provides a weak basis for prioritization, and thus retained all 85 top-hits for downstream analyses.

### Analysis of Copy Number Variants and Transposable Elements reveal new candidate selection signals during the Mesolithic-Neolithic transition

Next, we investigated the contribution of CNVs, which are often overlooked in aDNA studies^36^, by developing a depth-of-coverage framework tailored to such challenging data (see **Methods, Fig. 4a, Extended Data Fig. 16**). This analysis identified eight deletions and two duplications with significantly divergent allele frequencies (AFs) between HG and EF individuals **(Fig. 4b-d, Extended Data Fig. 17**). Additionally, we identified a duplication with a much lower AF than expected under the demographic mixture model in admixed individuals from nGer, suggesting directional selection (**Fig. 4e-h**). To assess whether these CNVs capture the same selective processes as the 85 top SNP hits (**Fig. 3**), we examined their genomic proximity (**Fig. 4i**). Most CNVs were not closely related to such SNPs, indicating that they mostly reflect independent, rather than redundant, events. These results suggest that CNVs add another layer of adaptation to Neolithic lifestyles.

**Fig. 3.**
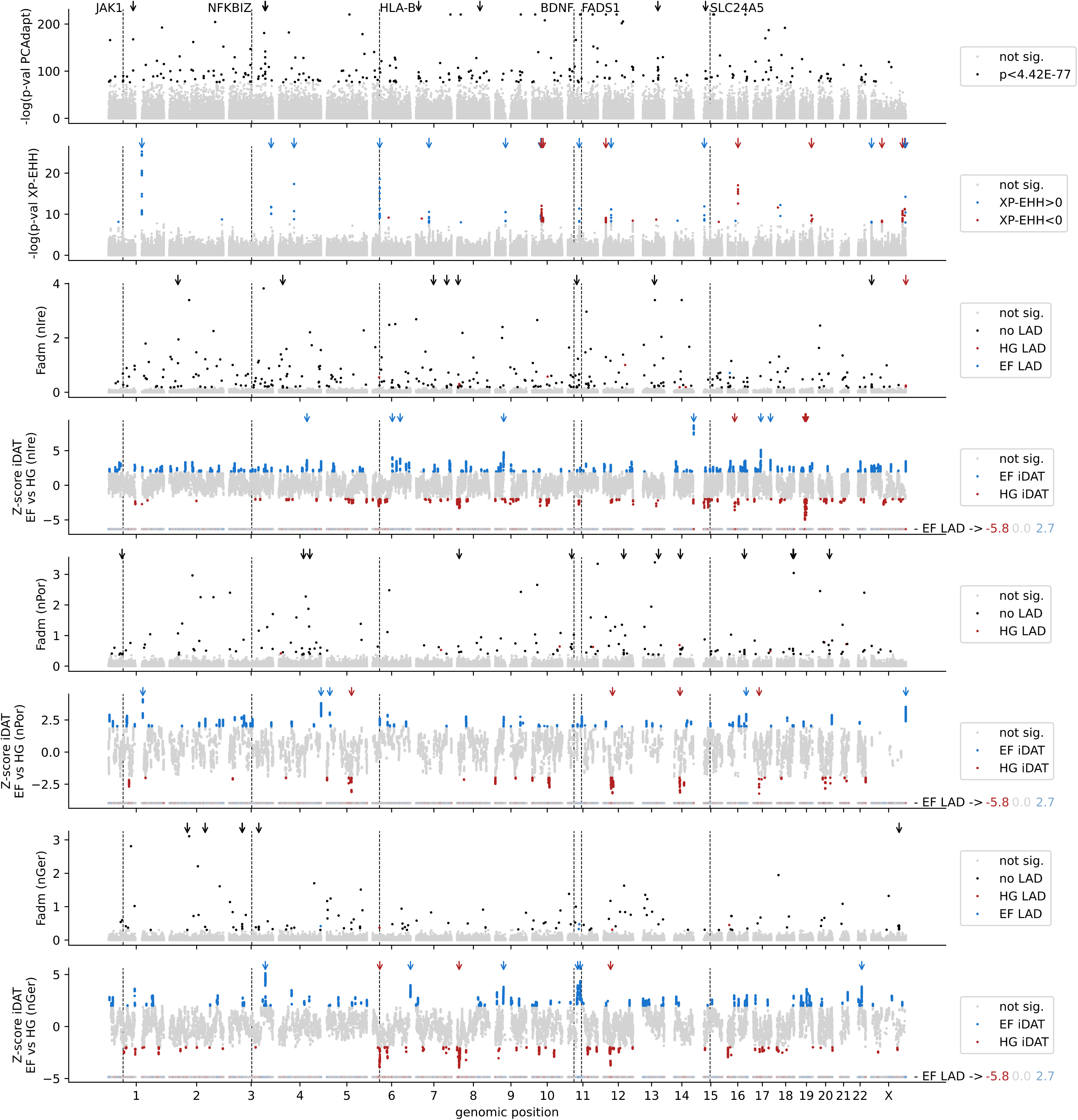
Overview of the SNPs under selection. Distribution of the significance metrics of each of the SNP selection scans across the genome (-log PCAdapt PC1 p-value, -log XP-EHH p-value, Fadm and iDAT scores). The gene names and vertical lines reflect the position of genes previously related to selection (see **Extended Data** Fig. 14**,15** for more details). Arrows indicate the top hits produced by each method, with color representing the directionality of selection (red for HG, blue for EF and black for unknown). See **Methods** and Fig. 2 for additional details.

**Fig. 4.**
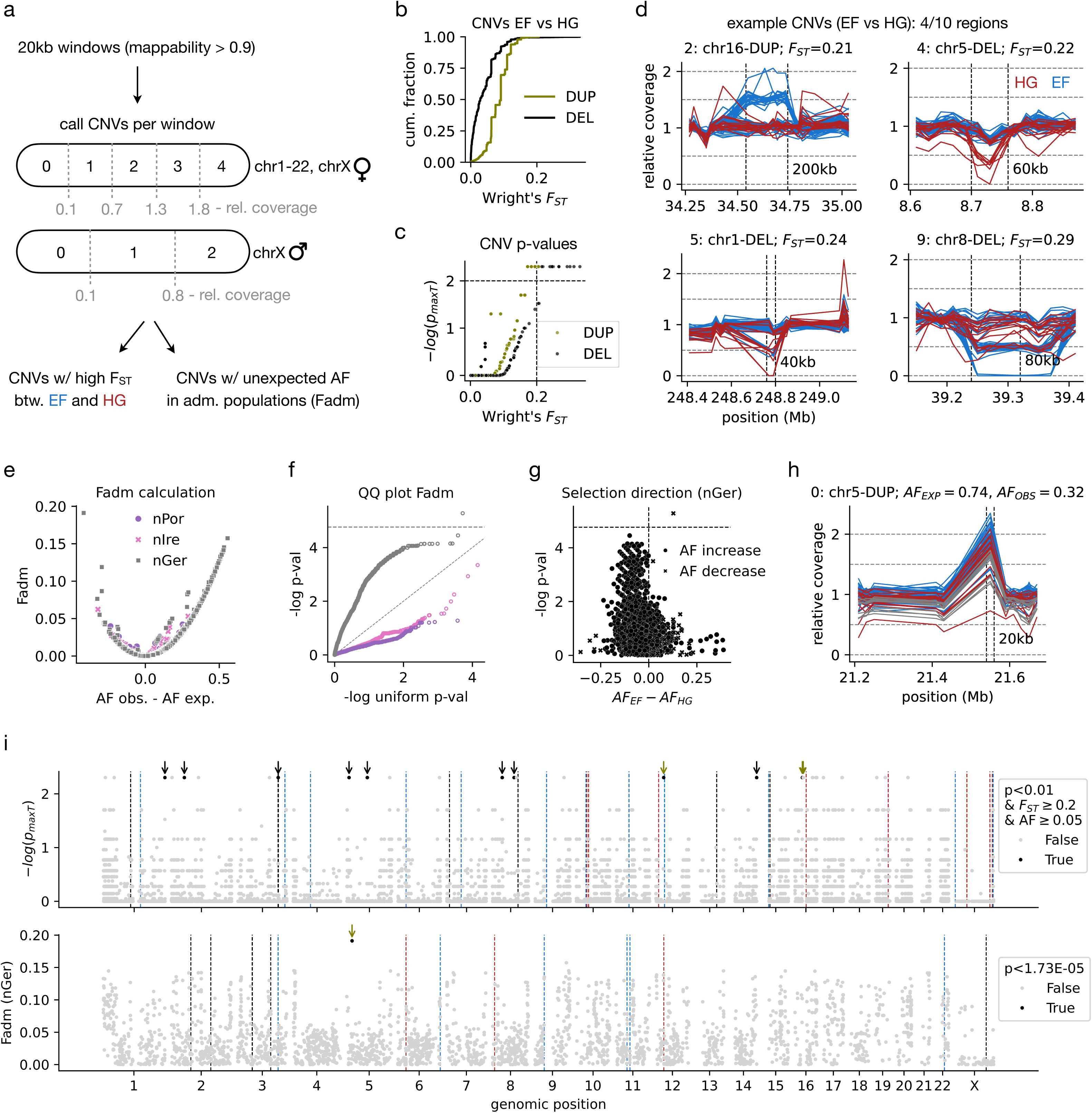
Copy Number Variation (CNV) analyses. **a.** Schematic representation of the CNV analysis pipeline. For 20kb windows with mappability > 0.9, we identified windows with CNVs (deletions or duplications) as those with a copy number (CN) different from 2 (for chromosomes 1-22 and females’ chromosome X) or 1 (for males’ chromosome X). The CN was inferred based on relative coverage following the thresholds in gray. Then, we identified CNVs with high Fst between EF and HG, or unexpected AF in each of the admixed populations as indicated by Fadm statistics. **b.** Distribution of the Fst for duplications (DEL) and deletions (DEL). **c.** For each CNV, Fst vs -log maxT p-values, indicating a probability, already corrected for multiple-testing, to observe such a high Fst by chance. The vertical and horizontal lines indicate the thresholds applied on these metrics to define significantly differentiated CNVs between the HG and EF populations. **d.** Examples of four significantly differentiated CNVs. Each line indicates the relative coverage for a given EF (blue) or HG (red) individual around the duplicated or deleted region. The vertical lines indicate the CNV region. To visualize the remaining CNVs check **Extended Data** Fig. 17. **e.** Fadm statistics calculation for all CNVs in each of the admixed populations (nGer, nIre and nPor). For each CNV, we calculated the AF observed or expected from a mixture model based on the genome-wide EF and HG ancestry proportions (AF difference in the x-axis), which are used to infer Fadm values (y-axis). **f.** QQ-plot of the p-values associated with the Fadm metrics, where the horizontal line reflects the Bonferroni-correction. **g.** Direction of selection for the CNVs in the nGer population. The x-axis reflects the difference in AF between ancestral EF and HG populations, while the y-axis indicates the -log Fadm p-value from **f**. Also, the symbols indicate whether there was an AF increase (circles) or decrease (crosses) vs the expectation from the mixture model. **h.** Relative coverage visualization for the one significant duplication in nGer, equivalent to **d**. **i.** Distribution of the significance metrics of each of the CNV selection scans yielding significant hits across the genome (-log maxT p-value, and Fadm in nGer). The vertical lines reflect top hit SNPs from Fig. 3, with the directionality of selection indicated by the color as in Fig. 2. Arrows indicate the CNV regions, where the color represents DEL / DUP as in **b**.

Similarly, we developed an aDNA-specific analytical framework, AncienTE, to quantify TE family abundances in ancient genomes (**see Methods, Fig. 5a, Extended Data Fig. 18,19,20, Supplementary Table 4**), which yielded 47 TE families displaying significant differences between HG and EF after Bonferroni correction (22 enriched in HG and 25 in EF; **Fig. 5 b,c**, **Extended Data Fig. 21**). These differences remained significantly associated with ancestry after controlling for sequencing coverage and sample age using linear models, indicating that these patterns are not explained by technical or temporal confounders (**Extended Data Fig. 22**). The differentially abundant families were dominated by evolutionarily young retrotransposons, including LTR/ERV1 (18 families), LINE/L1 (12), LTR/ERVK (9), SINE/Alu (6), one SVA and one DNA/TcMar-Tigger family. Notably, the composition of enriched superfamilies differed markedly between populations: HG-enriched families were primarily represented by LTR/ERVK and SINE/Alu elements, whereas EF-enriched families were predominantly LTR/ERV1 and LINE/L1. These findings reveal a marked shift in the superfamily composition of differentially abundant elements between populations.

**Fig. 5.**
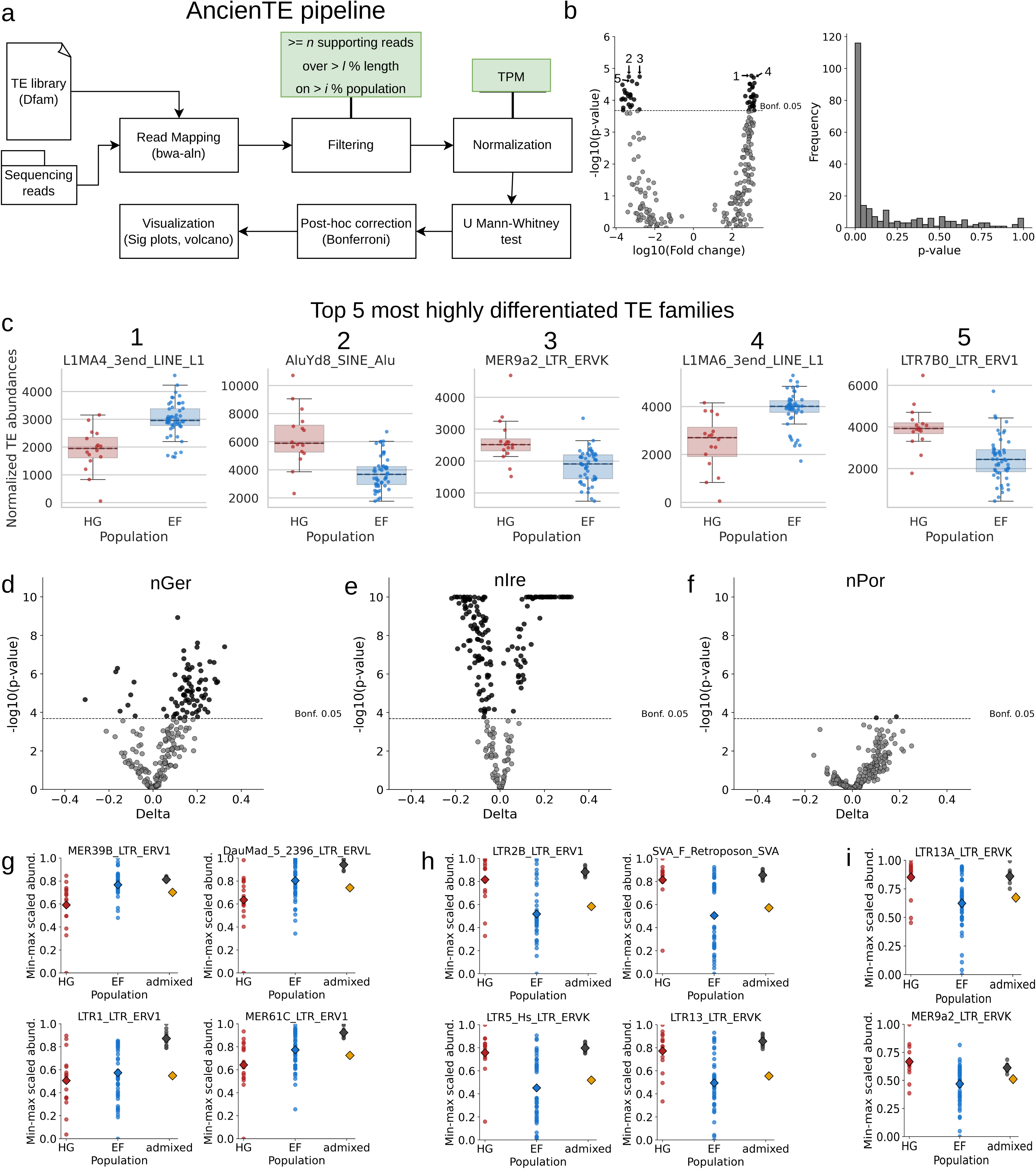
TE families showing significant abundance shifts in ancient populations. **a**. Schematic overview of the AncienTE pipeline. **b**. Volcano plot showing differential abundance of TE families between HG and EF individuals; the bar plot shows the distribution of p-values. The five TE families with the lowest Bonferroni-adjusted p-values are numbered. **c**. Boxplots illustrating abundance distributions of the five most strongly differentiated TE families in HG and EF individuals. Barplots for the 47 significant TE families can be found in **Extended Data** Fig. 21. Numbers correspond to the TE families indicated in **b**. **d-f**, Volcano plots showing TE families with significant deviations between observed and admixture-expected abundances in admixed individuals from **d**. nGer; **e**. nIre; and **f**. nPor. **g-i**. Observed versus expected abundances for the four most differentiated TE families in admixed individuals from g, nGer; h, nIre; and i, nPor (only two families reached statistical significance). Plots for all the significant TE families in the nGer population can be found at **Extended Data** Fig. 23**,24**.

To investigate TE dynamics following population admixture, we developed an ancestry-based expectation model that predicts TE family abundance from individual ancestry proportions (see **Methods**) and compared these expectations with observed values in admixed individuals. This analysis identified 78, 171 and 2 TE families with significant deviations from neutral ancestry-driven expectations in nGer, nIre and nPor, respectively **(Fig. 5d-i, Extended Data Fig. 23,24, Supplementary Table 5**). Strikingly, the vast majority of significant families in both nGer (87%) and nIre (73%) belonged to endogenous retroviral lineages, particularly LTR/ERV1, LTR/ERVK and LTR/ERVL (see **Supplementary Information**). These results show that TE abundance profiles not only capture broad ancestry-associated variation but also reveal regional departures from neutral admixture expectations. Despite the limitations imposed by highly fragmented ancient DNA, the consistent enrichment of specific retrotransposon families across admixed populations suggests that evolutionarily young TE lineages contributed disproportionately to genomic variation after admixture, providing a complementary perspective to SNP- and CNV-based analyses of ancient population history.

### Variant-phenotype associations provide functional insights into the selection process

To identify which of the 85 top SNP hits may reflect the most biologically relevant selective pressures, we assessed phenotype associations for these variants in COSMIC^37^, dbGAP^38^, ClinVar^39^, GEFOS^40^, GIANT^41^, MAGIC^42^, and the GWAS catalogue^43^ databases (see **Methods**, **Supplementary Table 2**). Three of the 85 SNPs were identified in COSMIC, and were associated with sites recurrently affected by somatic mutations in human liver and thyroid tumors. However, because COSMIC does not provide allele-specific information, their relevance to the selection process studied here remains unknown. By contrast, we found allelic-specific phenotype associations for three SNPs in the GWAS catalogue, enabling us to infer how selection may have shaped trait-associated variation during the Mesolithic-Neolithic transition. Although these associations do not establish causality, they provide valuable functional context for interpreting the selective signals. First, we identified rs2054349-C, an allele associated with decreased risk of neuroticism, which was more frequent in HG (AF_HG_=0.88, AF_EF_=0.42), and falls within HG haplotypes favored in the admixed populations from nGer (iDAT=-3.95) and nIre (iDAT=-2.58). Although highly speculative, this observation raises the possibility that some of the selective pressures acting in Neolithic societies involved behavioural or psychological traits related to sociability, as previously proposed^6,8,10^.

Second, we found rs6020624-G, an allele associated with lower haemoglobin levels, which was more frequent in EF (AF_HG_=0.63, AF_EF_=0.91), but was selected against in the admixed population of nPor (AF_EXPECTED_ =0.85, AF_OBSERVED_=0.31, Fadm=0.52). This pattern suggests that selection favored the alternative allele, generating a genetic predisposition toward higher haemoglobin levels. This signal may reflect selection on metabolic traits associated with shifts towards starch-rich, less diverse diets in Neolithic societies, potentially requiring improved handling of nutrient deficiencies^6,8,10,11^. In particular, lower intake of red meat may have increased the risk of iron deficiency, potentially favoring alleles associated with improved iron handling or higher haemoglobin levels^44–46^. Our findings may therefore represent an additional example of selection on iron-related physiology during the Neolithic transition.

Third, we identified rs6476690-A, an allele negatively correlated with bacterial abundance in the gut microbiome, which was more frequent in EF (AF_HG_=0.25, AF_EF_=0.50), and lies within EF haplotypes favoured in the admixed population of nIre (iDAT=4.77). This finding may reflect selective pressures on the immune system, and is consistent with the hypothesis that more frequent contact with domesticated animals during the Neolithic would have increased pathogen exposure, leading to selection for more reactive immune responses despite the tradeoff of increased risk towards inflammatory, autoimmune and cardiovascular diseases^6–10,47^. In this context, the preferential selection of EF haplotypes carrying alleles associated with lower bacterial abundance may reflect adaptation to a more pathogen-rich environment.

All in all, the analysis of variant-phenotype association suggests that selection during the Mesolithic-Neolithic transition shaped traits related to sociability, haemoglobin metabolism, and pathogen exposure. Most notably, we predict that these phenotypes were shaped by both HG and EF alleles under selection, implying a relevant contribution of both ancestral genetic pools to the adaptive process.

### Ancient pathogen screening reveals an early Neolithic Hepatitis B viral infection

To explore the potential differential presence of pathogens in our samples, we analyzed sequencing reads that did not map to the human genome (see **Methods**). A recent study empirically showed that pathogen load increased across the Mesolithic-Neolithic chronology^7^, but it remains unclear whether this increase reflected differences between ancestral EF / HG populations. To clarify this while minimizing pathogen survivor biases, we filtered out samples with DNA extracted from petrous bones, as these are less likely to have been colonized by ancient pathogens^7^. This resulted in 5 HG and 1 EF, including i) some of the newly sequenced Iron Gates individuals, and ii) three AADR HG individuals from Northern European countries (see **Methods**). Because geographical distance may influence pathogen dispersal, we only analyzed the newly reported Iron Gates individuals (2 HG, 1 EF) to ensure a geographically balanced comparison. While this is a small dataset not allowing for a rigorous statistical comparison, it provides an opportunity to infer whether pathogen exposure may already have differed between co-localized ancestral groups.

None of the samples provided clear evidence for bacterial infections. However, the one EF individual (i701-i501) presented 2061 reads assigned to Hepatitis B virus (HBV). Subsequent read recruitment against the HBV genome confirmed this signal, yielding 731 reads with high quality mappings, resulting in 9.8x coverage. Consistently, skeletal remains of this individual show signs of femoral bone deterioration (**Supplementary Information**) as observed in modern chronic hepatitis patients^48^. Variant analysis classified this HBV strain as belonging to the WENBA lineage (**Fig. 6**), which spread during the Neolithic transition^19^, supporting the idea that i701-i501 was infected by HBV.

**Fig. 6.**
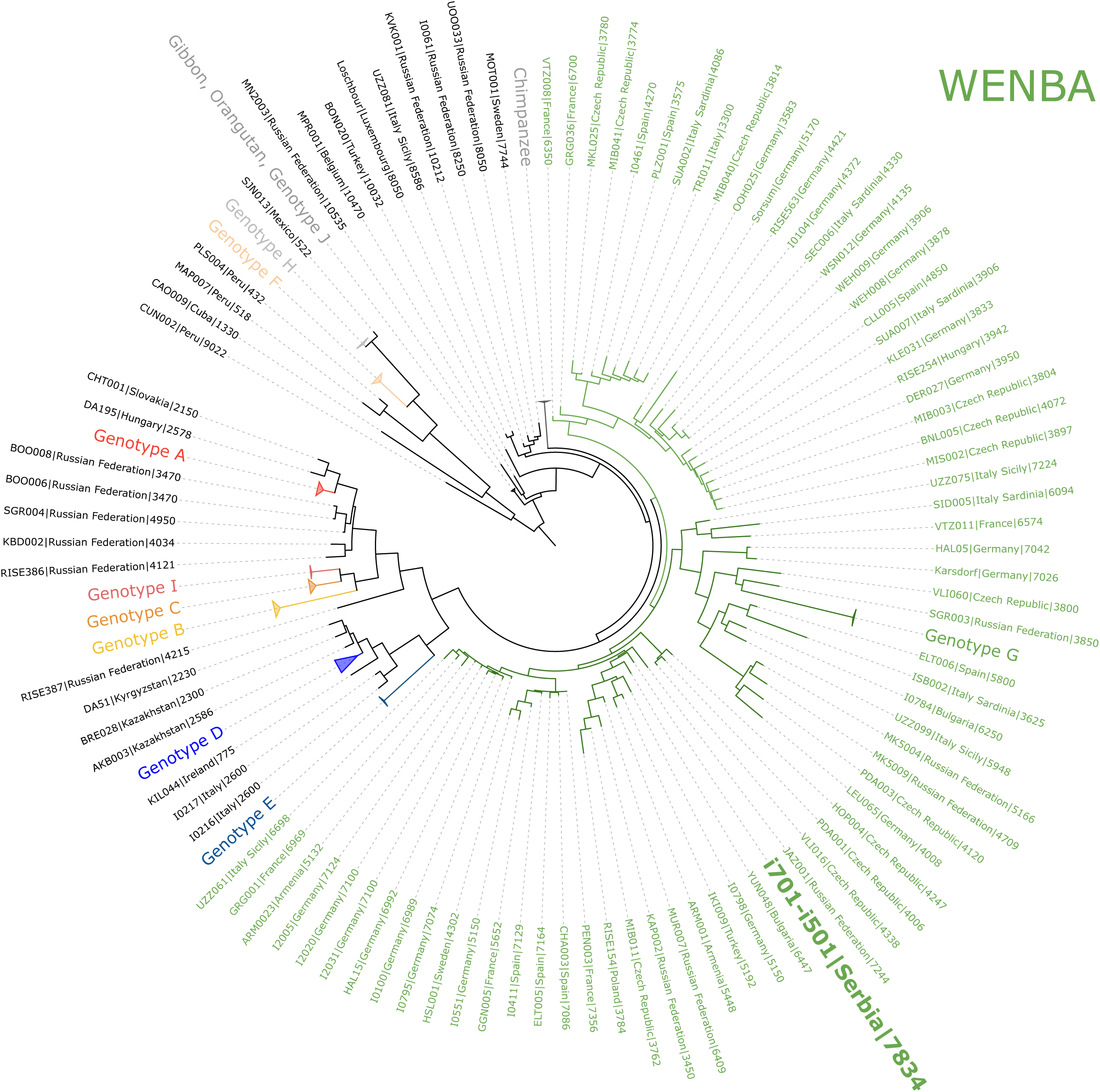
Phylogenetic analysis of Hepatitis B Virus (HBV). Maximum Likelihood tree was constructed with the HBV dataset presented in ^19^, including the newly reconstructed genome i701-i501, indicated in bold and bigger font size. Colours correspond to different HBV genotypes: genotype A (red), genotype B (yellow), genotype C (orange), genotype D (blue), genotype F (soft orange), genotype H (grey), genotype I (soft red), genotype J (dark grey, including HBV isolated from Gibbons and orangutans), Chimpanzee isolates (dark grey), WENBA lineage (green). The tree was visualised with iTOL^178^ (v.7).

Notably, radiocarbon dating of this individual places the HBV strain very early in the WENBA lineage (at 5986-5783 BCE, see **Supplementary Table 1**), making this HBV strain one of the earliest representatives of the WENBA lineage and roughly contemporaneous with the previously described Mesolithic 2 haplotype^19^. These findings indicate that the WENBA haplotype was circulating in the Balkans earlier than previously reported^19^, likely introduced by EF individuals migrating from the Fertile Crescent. This further supports the role of the Balkans as an early contact zone during the Neolithic expansion and refines the phylogeographic history of HBV in western Eurasia. Although the limited sample size prevents strong conclusions about differences in pathogen burden between HG and EF populations, the presence of chronic HBV infection in an EF individual is consistent with the hypothesis that EF populations experienced increased pathogen exposure during the Mesolithic-Neolithic transition. More broadly, our analyses showcase the potential of WGS approaches to study host-pathogen coevolution from aDNA data.

### Functional enrichments link selected variants to adaptive biological processes

Variant-phenotype correlations provided functional context for only three of the 85 top hit SNPs under selection. To broaden this analysis, we predicted the impact of top hit SNPs and differential CNVs on nearby genes and other genomic elements using the Ensembl Variant Effect Predictor (VEP)^49^. Additionally, we identified SNPs annotated as expression quantitative trait loci (eQTLs) in the GTEx database^50^, to infer potential transcriptional effects (see **Methods, Fig. 7, Supplementary Table 2**). With the exception of a missense mutation in *RIPOR3*, the top SNPs do not alter protein sequences and were mapped to 243 nearby genes (**Fig. 7a**), suggesting that much of the adaptive signal may have operated through regulatory rather than protein-altering changes. We therefore focused on 174 genes predicted to be differentially regulated by 44 of the top SNPs.

**Fig. 7.**
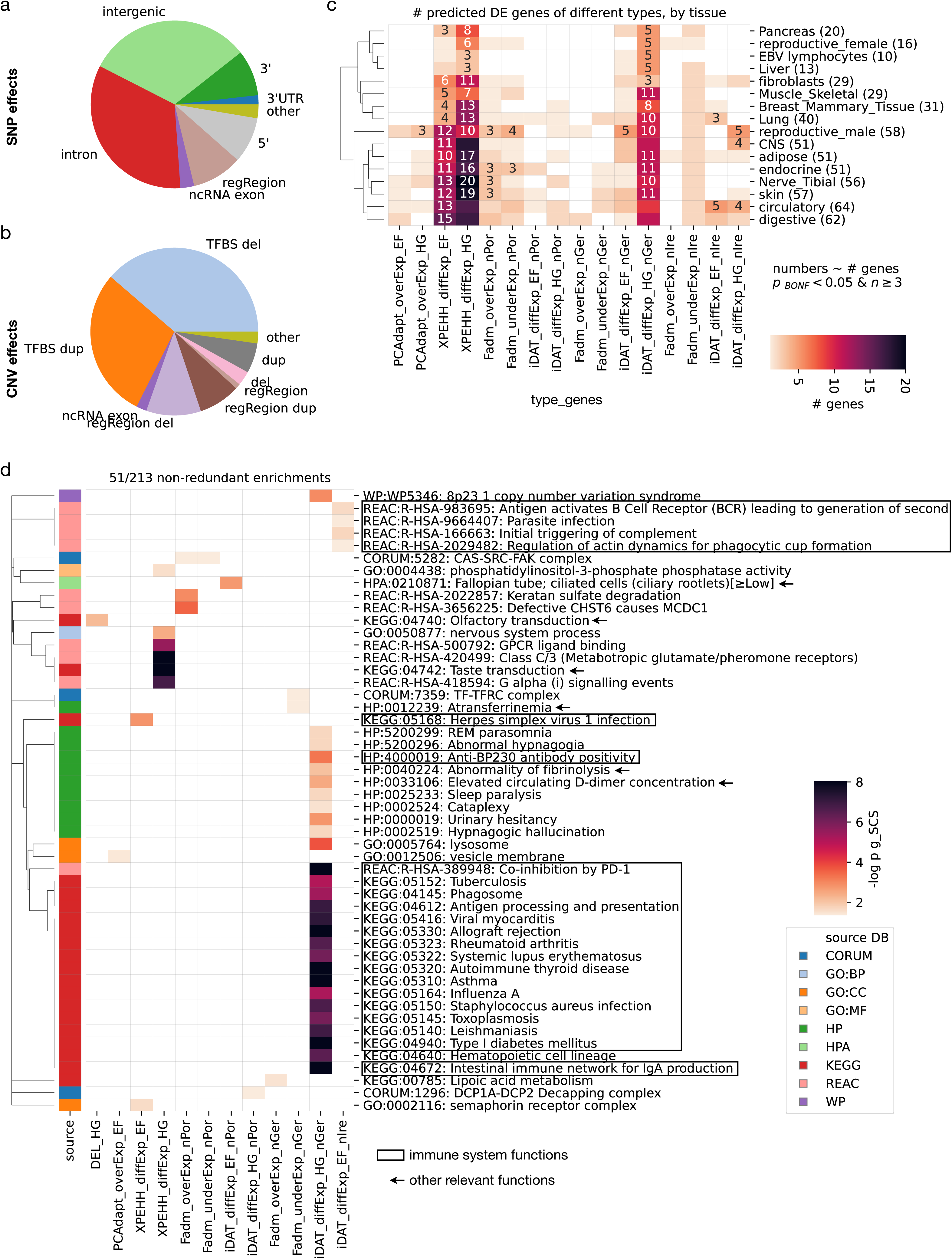
Functional variant effects. **a,b.** SNP and CNV functional effects on nearby genes, as predicted by the Variant Effect Predictor. **c.** Heatmap showing the number of predicted differentially expressed genes of each type (columns) in each tissue (rows), according to eQTL annotations. Each gene type corresponds to a set of genes predicted to be differentially expressed by certain SNPs under selection. For instance, we identified genes where the allele that leads to their predicted upregulation is a PCAapt hit, and has higher AF in either EF (gene set=PCAdapt_overExp_EF), or HG (gene set=PCAdapt_overExp_HG). In addition, we focused on genes with predicted changed expression by alleles under selection by XP-EHH in EF (gene set=XPEHH_diffExp_EF), or HG (XPEHH_diffExp_HG). Furthermore, we identified genes where the allele that leads to their predicted upregulation (gene set=Fadm_overExp_<POPULATION>) or downregulation (gene set=Fadm_underExp_<POPULATION>) increased AF in admixed individuals from a given analyzed admixed populations (nIre, nGer or nPor). Finally, we defined genes with predicted changed expression by EF alleles (gene set=iDAT_diffExp_EF_<POPULATION>) or HG ones (gene set=iDAT_diffExp_HG_<POPULATION>) favoured in admixed individuals from each population, according to iDAT. Cells where the number is shown reflect tissues where there are >=3 genes, and it is highly unlikely (empirical Bonferroni-corrected p<0.05) to find the observed number of genes when randomly drawing n eQTLs from the whole GTEx database, where n is the number of eQTLs related to a given type of genes. The number in parenthesis reflects the sum of the number of DE genes across all gene sets. **d.** Non-redundant functions enriched in each of the types of genes, using the standard g_SCS method from G:Profiler. In **Supplementary Table 6** you can find all of them. The boxes indicate immune functions, while the arrows point to other relevant processes discussed in the text. The row colors indicate the source database of the terms.

To integrate the different selection signatures and eQTL annotations, we grouped genes predicted to be differentially expressed (in any tissue) into 16 gene sets depending on the identification method (PCAdapt, XP-EHH, Fadm, and iDAT) and enriched population (HG, EF, or a particular admixed neolithic population) (see **Methods, Fig. 7**). Each of these gene sets may capture biological processes under selection in different evolutionary periods and populations. We next asked whether these gene sets particularly involved certain tissue groups (i.e. sets of tissues of the same system, e.g. grouping all CNS or circulatory system ones, see **Methods**). For each gene set we calculated the number of predicted differentially expressed (DE) genes in each tissue group, and we focused on gene sets and tissue groups with i) at least three DE genes, and ii) a low empirical probability (Bonferroni-corrected p<0.05) of observing such a number of DE genes or higher by chance (see **Methods**) (**Fig. 7c**). This yielded 16 tissue groups, 11 of which were related to three or more gene sets, including pancreas, fibroblasts, skeletal muscle, mammary tissue, lung, male reproductive system, adipose tissue, endocrine system, tibial nerve, skin, and circulatory system.

Several of the affected tissues are consistent with expected selective pressures during the Mesolithic-Neolithic transition. For instance, alterations in the pancreas (three gene sets) may relate to insulin metabolism, expected to be affected by selection during adaptation to starch-rich diets in EFs^6,14,51^. Changes affecting the male reproductive system (eight gene sets) may reflect selection on fertility during this transition, consistent with a proposed increase in fertility rates during the Neolithic^52,53^. Finally, changes in the skin (four gene sets) are expected by the hypothesis that a reduction in skin pigmentation was adaptive in Neolithic societies due to reduced dietary vitamin D intake^12,13^.

We next examined the functions enriched in each gene set using g:Profiler^54^ across GO, KEGG, Reactome, WikiPathways, Human Protein Atlas, CORUM and HP annotations (**Fig. 7d**, **Supplementary Table 6**). We identified 209 enrichments (203 unique terms) across 11 of the 16 gene sets, which we grouped into 50 non-redundant enrichments with 49 unique terms using a REVIGO-like^55^ approach (see **Methods**, **Fig. 7d**). Most enrichments (203/209) were specific to a gene set, indicating that distinct selective episodes affected different biological processes across evolutionary timepoints and locations (**Fig. 3**). The differences observed across geographies underscore the importance of studying adaptive events at a local scale to uncover the fine-scale complexities of human adaptation, as previously proposed^56^.

Despite this heterogeneity, some general trends emerged. Most prominently, immune-related functions were recurrently enriched, including antigen recognition and processing, and pathways related to infectious and autoimmune diseases (**Fig. 7d**). These findings provide further support for the hypothesis that increased pathogen exposure favored selection on immune responsiveness (discussed above), consistent with earlier reports of immune gene selection during the Neolithic^6,12–14^. Notably, immune enrichments were detected in gene sets linked to selection in EF, but also in the admixed populations of nIre and nGer favouring EF and HG alleles, respectively. This indicates that pathogen adaptation did not simply involve a single selective sweep favouring “pathogen-adaptive” EF variants, but rather was a more complex evolutionary process involving both ancestries. For instance, the selected EF allele rs11849201-G in the nIre admixed population (iDAT=8.65), with a predicted impact on immunoglobulin genes *IGHV1-69* and *IGHV2-70*, may have been already mildly selected in the EF population (AF_HG_=0.09, AF_EF_=0.30, pcadapt p<0.03), perhaps reflecting an equivalent adaptive process. Conversely, the HG allele rs9274229-A (AF_HG_=0.91, AF_EF_=0.64), which affects various subunits of the Major Histocompatibility Complex class II encoded by *HLA* genes, is specifically selected in nGer (iDAT_GER_=-3.89, iDAT_IRE_=-0.56, iDAT_POR_=0.21).

Such population-specific patterns may explain previous inconsistencies regarding the directionality of selection on *HLA* genes, considered a major responsible for pathogen adaptation in this period^12,13^. Comparison with previous studies^12,13^ confirmed the selective sweeps involving *HLA-B, HLA-DQB1* and *HLA-DBQB2* but not those related to *HLA-A* or *HLA-E*, and our study uncovered a novel one involving *HLA-DRB5* (**Extended Data Fig. 14**). With respect to the directionality of selection, we find that *HLA-B* HG alleles are selected in admixed populations from nIre and nGer, consistent with previous findings of selection on this gene in HG populations^12^. Similarly, around *HLA-DQB1* we detect selection of HG alleles in the nGer population (rs9274229-A described above), which matches previous analysis of adaptive admixture^13^. In contrast, we infer selection of HG alleles around *HLA-DQB2* in the nGer population, whereas previous studies reported selection of EF alleles^12^. Finally, we detect XP-EHH-related selection in EFs overlapping an enhancer linked to *HLA-DRB5* (on rs112687068-C), and this region also has signs of i) selection of EF alleles in the nPor population, and ii) selection of HG alleles in the nGer population. Together, these results suggest that the previously reported inconsistencies in the direction of *HLA*-related selection reflect a genuinely complex evolutionary landscape, with multiple selective sweeps acting at different times and in different populations. The finding that HG *HLA* alleles can be under selection, even against EF alleles selected in the ancestral EF population (as for rs112687068-C), shows the importance of HG ancestry in the adaptive process. We propose that EF alleles may have helped incoming EF groups cope with pathogens associated with neolithic lifestyles, whereas HG alleles may have contributed adaptation to local pathogen environments in specific regions, as suggested in ^13^.

Beyond immunity, our enrichment analysis revealed trends related to fertility, diet, and hematological adaptation. Genes with predicted DE by EF alleles selected in the nPor population have enriched expression in ciliated cells of the fallopian tube, which are important for oocyte collection and fertility^57^. This finding might reflect selection for higher intrinsic fertility rate, a proposed driver of the population increase during the Neolithic^52,53^. Additionally, genes with predicted DE by alleles under selection in HG are enriched in taste transduction functions, perhaps related to their highly distinct diet as compared to EF. Finally, we observe various hematological functions enriched in genes under selection in the nGer population, including ‘atransferrinemia’, ‘abnormality of fibrinolysis’, and ‘elevated circulating D-dimer concentration’. These may reflect changes in hematological parameters, similar to the selective process on haemoglobin levels in the nPor population (rs6020624-G) discussed above.

We also explored the functional relevance of differential CNVs, which are near 60 genes (**Fig. 4,7b Supplementary Table 2**). First, we assessed the variant effects of the three detected duplications. Two of them affected only pseudogenes and predicted regulatory regions (**Fig. 4 h, Extended Data Fig. 17**), suggesting that they may have resulted from genetic drift and a lack of purifying selection against them. We thus focused on the duplication 16:34540000-34740000_DUP, more frequent in EF (**Fig. 4d**), which results in the amplification of various pseudogenes and of the intergenic lncRNA (*LINC01566*). This lncRNA has a proposed role as a protective factor towards the autoimmune disease myasthenia gravis^58^. Its higher frequency in EF may result from selection favouring protection against autoimmune diseases, which could have been key to counteracting the tradeoffs associated with having more reactive immune systems to better handle pathogens (discussed above).

Second, we assessed variant effects of deletions using functional enrichment analysis for genes close to deletions with higher AF in either EF or HG. We could not find significant enrichments for the two deletions with higher AF in EF, and they both overlapped regulatory regions or pseudogenes, suggesting uncertain functional impacts. Conversely, for genes affected by the six deletions with higher AF in HG we found functional enrichments related to “olfactory transduction” (**Fig. 7d, Supplementary Table 6**). Notably, the only transcript-ablation effect associated with this enrichment involves a single deletion (1:248760000-248800000_DEL, see **Extended Data Fig. 17**) overlapping the olfactory receptor gene *OR2T11*. This may reflect differences in olfactory perception which, similar to the ones in taste transduction described above, may have been under selection in HG due to their highly distinct diets or lifestyles as compared to EF. Overall, mapping selected variants to nearby genes and predicted regulatory effects reveals recurrent signatures in immunity, fertility, and metabolism. These results expand the interpretation of the selection signals beyond individual loci and suggest that adaptation during the Mesolithic-Neolithic transition involved multiple biological systems, often in region-specific ways.

## CONCLUSIONS

By combining available ancient human genomes with newly generated data from the Iron Gates, we assembled a large high-coverage WGS dataset to study selection during the Mesolithic-Neolithic transition. Compared with previous SNP-panel studies, our dataset enabled a genome-wide view of adaptation and substantially increased the representation of HG genomes. Our analyses identify the Iron Gates as an important early zone of interaction and admixture between incoming EF and local HG groups, and show that later admixed neolithic populations cannot be treated as homogeneous. Instead, our data support geographically structured adaptive admixture, with both HG and EF alleles contributing to adaptation in late Neolithic groups. This finding challenges simplified views in which the adaptive potential required for the transition was carried primarily by EF ancestry.

Using four complementary SNP selection scans, we uncover signals that both confirm previously proposed adaptive trends and point to novel candidates. Notably, the limited overlap among scans suggests that selection during this transition was complex, acting differently across populations and evolutionary timepoints. This spatiotemporal complexity may also explain the limited overlap among previous studies. To leverage the full potential of WGS data beyond SNPs, we established novel approaches to study CNVs and TEs in ancient genomes. This uncovered additional signals of selection, which underscores the feasibility and importance of studying generally-overlooked structural variation. WGS also enabled analysis of pathogen DNA, refining the early divergence of HBV.

Using a novel approach that predicts phenotypic and transcriptomic effects of selected variants beyond nearby genes, we infer selection on both expected and novel biological processes. Our findings on immune-related signals align with previous evidence for selection toward a more reactive immune system during the Neolithic, likely in response to higher pathogen exposure, despite increased risk for autoimmune and cardiovascular diseases. We also identify novel signals involving reproduction, haemoglobin metabolism, sensory perception, and behaviour. These may reflect selection on fertility, dietary change, and the social demands of increasingly structured Neolithic societies. Notably, both HG and EF alleles contributed to adaptation in late Neolithic populations across these processes, highlighting the importance of studying HG ancestry and the value of our dataset.

More broadly, our findings support the importance of regional analyses for understanding the Mesolithic-Neolithic transition. Selective processes were likely heterogeneous across space and time, and may be obscured by commonly used large-scale continental models. Although the functional inferences presented here remain to be experimentally validated, and larger sample sizes will be needed to refine these patterns, our study provides both a new empirical resource and a broader framework for investigating how admixture, structural variation, and local ecological pressures shaped human adaptation during this major prehistoric transition.

## MATERIALS AND METHODS

### Code and software environments

All code required to run the analysis described below is available on indicated sources or (mainly) in the GitHub repository https://github.com/Gabaldonlab/MesoNeoSelection. Regarding the code in the latter repository, we used the script pipelines_IronGates.py to generate processed datasets and paper_IronGates.py to generate plots and tables. Also, we ran various scripts from the folder ‘scripts’ to perform various specific analysis steps, as described below (e.g. map_reads.py, get_adapters.py, run_pcadapt.R or AncienTE.py). These scripts mostly rely on IronGates_functions.py, a module comprising all used python functions. In the sections below we refer to specific functions in this module as IGfun.<function name>. We ran most of this code on the conda environment ‘IronGates_env’, defined in the folder ‘envs’ of the repository, which includes matplotlib^59^ v3.8.0, numpy^60^ v1.26.0, python^61^ v3.9.7, scipy^62^ v1.11.3, seaborn^63^ v0.13.2, biopython^64^ v1.78, pandas^65^ v2.1.1, openpyxl^66^ v3.0.10, statsmodels^67^ v0.14.0 scikit-learn^68^ v1.3.0, sra-tools^69^ v3.0.5, plink^70^ v1.90b6.18, vcftools^71^ v0.1.16, goatools^72^ v1.1.12, gprofiler-official^73^ v1.0.0, matplotlib-venn^74^ v0.11.10, gsl^75^ v2.5, openblas^76^ v0.2.20 and fftw^77^ v3.3.8.

Beyond this main environment, various parts of the pipeline were run on various other conda environments (contammix_env, IronGates_R_env, ariadna_env, gargammel_env, perSVade_env, perSVade_env_picard_env, perSVade_env_RepeatMasker_env, AncienTE_env, eager_env), defined next. The ‘contammix_env’ includes ContamMix^78^ v1.0.11, samtools^79^ v1.12, iVar^80^ v1.4 and mafft^81^ v7.525. The ‘IronGates_R_env’ includes r-base^82^ v4.3, r-pcadapt^32^ v4.3.5, r-argparser^83^ v0.7.1, R’s package rehh^33^ v3.2.2, r-fitdistrplus^84^ v1.1_11. The ‘ariadna_env’ includes python v2.7.15 and scikit-learn v0.18.2. The ‘gargammel_env’ includes python v2.7.15, ms^85^ v2014_03_04, seq-gen^86^ v1.3.4, numpy v1.16.6, scipy v1.2.1, biopython v1.74, msprime^87^ v0.3.1 and gargammel^88^ v1.1.2. The environments starting with ‘perSVade_’ are those related to the perSVade pipeline^89^ (commit ‘e3bf913’ from 09/09/2025 at https://github.com/Gabaldonlab/perSVade). The AncienTE_env, includes the following dependencies: python v3.10.16, pysam^90^ v0.23.3, tqdm^91^ v4.67.1, Biopython v1.86, SciPy v1.15.2, scikit-learn v1.7.2, Matplotlib v3.10.8, and Seaborn v0.13.2.

The ‘eager_env’ includes nextflow^92^ v21.10.6, python v3.9.4, markdown^93^ v3.3.4, pymdown-extensions^94^ v8.2, pygments^95^ v2.14.0, bioconda rename^96^ v1.601, openjdk^97^ v8.0.144, fastqc^98^ v0.11.9, adapterremoval^99^ v2.3.2, adapterremovalfixprefix^100^ v0.0.5, bwa^101^ 0.7.17, picard^102^ v2.26.0, samtools v1.12, dedup^103^ v0.12.8, angsd^104^ v0.935, circularmapper^105^ v1.93.5, gatk4^106^ v4.2.0.0, gatk^106^ v3.5, qualimap^107^ v2.2.2d, vcf2genome^108^ v0.91, damageprofiler^109^ v0.4.9, multiqc^110^ v1.16, pmdtools^111^ v0.60, bedtools^112^ v2.30.0, libiconv^113^ v1.16, pigz^114^ v2.6, sequencetools^115^ v1.5.2, preseq^116^ v3.1.2, fastp^117^ v0.20.1, bamutil^118^ v1.0.15, mtnucratio^119^ v0.7, pysam v0.16.0, kraken2^120^ v2.1.2, pandas v1.2.4, freebayes^121^ v1.3.5, sexdeterrmine^122^ v1.1.2, multivcfanalyzer^123^ v0.85.2, hops^124^ v0.35, malt^125^ v0.61, biopython v1.79, xopen^126^ v1.1.0, bowtie2^127^ v2.4.4, eigenstratdatabasetools^128^ v1.0.2, mapdamage2^129^ v2.2.1, bbmap^130^ v38.92, bcftools v1.12^131^, and rfmix^132^ v2.03.

We provide the .yml files to reproduce all the environments generated in this work in the folder ‘envs’ of the GitHub repository. In the sections below we refer to the specific environments used for each step, using the text “env:name” in parenthesis. Note that for sections in which the environment is not mentioned were run in the main ‘IronGates_env’. Also, we used various extra software tools, described next. First, for variant calling-related operations we used the nf-core/eager nextflow pipeline^133^ (v2.5.0), downloaded at https://github.com/nf-core/eager/archive/refs/tags/2.5.0.tar.gz. Second, we called variants with AriaDNA^134^ (v5), downloaded from https://osf.io/5bph4/. Third, for variant functional annotations we used the Ensembl Variant Effect Predictor (VEP)^49^ singularity image (v111.0), installed with ‘singularity build --docker-login ensembl-vep_111.0.sif docker://ensemblorg/ensembl-vep:release_111.0’. Fourth, for integrated Decay in Ancestry Tract (iDAT) scans we based our code on the one from https://github.com/agoldberglab/CV_DuffySelection (commit ‘21a3a6b’ from 01/10/2025)^135^. Fifth, to estimate admixture timings we used DATES v753 from https://github.com/priyamoorjani/DATES, customizing the Makefile before compilation to match our IronGates_env environment, as explained in the IronGates_env.yml provided in our GitHub repository. Sixth, for SNP imputation we used the GLIMPSE2^136^ singularity image (v2.0.0), installed with ‘singularity build --docker-login glimpse_2.0.0.sif docker://simrub/glimpse:v2.0.0-27-g0919952_20221207’.

### Selection of publicly available individuals for analyses

From the publicly available and curated ancient genomes compiled in the Allen Ancient DNA Resource (AADR)^22^ (version 62; accessed Dec. 2024, metadata available in AADR_v62.0_1240k_public.anno), we selected individuals with Whole Genome Shotgun sequence data, and dated up to 14,000 years before present (BP), which include Mesolithic and Neolithic samples. We then applied a minimum genome-wide coverage threshold of 0.5x to ensure sufficient data for imputation^31^. To focus our analysis on Western Eurasian (WE) populations and exclude very early or genetically distinct hunter-gatherer groups, we removed individuals designated as Paleolithic by archaeological context and chronology (as annotated in the AADR metadata). We further excluded individuals flagged in the AADR for low genome-wide coverage or evidence of contamination (as annotated in the AADR metadata; under fields: ‘SNPs hit on autosomal targets’, ‘Damage rate in first nucleotide on sequences overlapping 1240k targets (merged data)’, ‘hapConX 95% CI truncated at 0 (only if male and >=2000 SNPs covered on X chromosome)’, ‘SNPs hit on autosomal targets’, ‘ASSESSMENT’ and ‘ASSESSMENT WARNING’).

Subsequent filters targeted population structure and genetic ancestry to minimize confounding effects. We removed individuals identified as genetic outliers within their assigned populations (individuals considered are flagged as genetic outliers as annotated in the AADR metadata; under field: ‘Group ID’ ). To ensure a dataset of unrelated individuals, we pruned one individual from any pair with kinship relationships exceeding a first-degree relative threshold (as annotated in the AADR metadata, under field: ‘Group ID’). To specifically isolate patterns not driven by steppe-related ancestry, we systematically removed all individuals associated with the Yamnaya culture (as annotated in the AADR metadata, under field: ‘Group ID’). As this ancestry was introduced into the continent after the Neolithic spread^137^, with this filter, we aimed to isolate the selective signal of the Anatolian Neolithic-like ancestry spread. After these filters, we projected the remaining individuals to a West Eurasian Principal Component Analysis (PCA) space on the remaining dataset computed using a set of 1081 present-day West-Eurasian individuals genotyped using the Human Origins array (using smartpca tool in EIGENSOFT software version 8.0.0 with lsqproject=YES and autoshrink=YES^138,139^, to identify and remove any additional outliers not captured by prior filters.

To refine the geographic and temporal scope of our study, we excluded all individuals from regions representing major steppe and immediately surrounding regions (tagged as present-day Russia, Ukraine, Moldova, Armenia, Georgia). We also removed individuals from the Early Bronze Age to focus on earlier Neolithic periods. Finally, we excluded specific individuals and groups (France_HautsDeFrance_LN.SG, Croatia_Popova_CA.SG, Germany_Niedertiefenbach_Wartberg_LN.SG, Germany_LN_Alberstedt.SG) that either showed significant steppe admixture or were genetically heterogeneous (as shown by their PC Values in the Wes-Eurasian PCA).

This curation process resulted in a final dataset of 201 publicly available whole shotgun-sequenced genomes.

### DNA extraction and sequencing of newly sequenced individuals

A total of twelve individuals were analysed in this study (**Supplementary Table 1**). Ancient DNA was sampled, extracted and converted into sequencing libraries in the Ancient DNA Laboratory at the Core Facilities of the Institute of Genomics, University of Tartu.

#### DNA sampling and extraction

Skeletal elements included multiple tissue types: a) the petrous portion of the temporal bone, b) dentine, and 3) auditory ossicles (incus and malleus). For each sample, one or more subsamples were taken and processed independently.

All sampling was performed in a dedicated clean hood under strict contamination control. Work surfaces and equipment were routinely decontaminated using DNA Exitus, and metal tools were treated by immersion in 6% sodium hypochlorite, followed by rinsing in MilliQ water and 70% ethanol and air-drying prior to use. Sampling areas were covered with disposable aluminium foil, replaced between specimens, and consumables were single-use whenever possible. Petrous bones and teeth were mechanically cleaned prior to sampling. When present, auditory ossicles were removed from the petrous bones and retained separately.

Petrous sampling targeted the dense inner ear region and was adapted to preservation: in well-preserved specimens, a rotary drill fitted with a narrow drill bit was used to access the cochlear region at an oblique angle (∼25°), yielding a compact core and associated bone powder; in smaller or fragile specimens, a cutting disc was used to isolate a 3–4 mm bone section; and in poorly preserved material, powder was obtained by controlled abrasion of the internal surface.

Tooth sampling targeted the pulp chamber of permanent teeth. Teeth roots were sectioned at the enamel–cementum junction. A sterile drill with a ball-end bit was used to access the pulp chamber, and dentine powder was collected.

Petrous cores and ossicles were decontaminated^140^, and DNA was extracted as described in ^141^. DNA from bone and dentine powder was extracted in a dedicated clean hood as follows. Samples (typically ≤250–300 mg) were incubated in 0.5 M EDTA (pH 8.0) and Proteinase K (approximately 2 ml and 50 µl per 100 mg of powder, respectively) to facilitate decalcification and protein digestion.

Extraction mixtures were incubated on a rocker at room temperature for 24 h. All reagents and consumables were handled under strict contamination control, including prior surface decontamination and UV-exposure where appropriate. Extraction blanks were included to monitor potential contamination.

#### DNA library preparation

All extracts were purified^142^ and built into genomic libraries using the established double-stranded library preparation method of Meyer & Kircher^143^. Library preparation input volumes were typically 20–30 µl per extract, and no UDG treatment was applied. In all cases, extracts underwent inhibitor removal prior to library construction using the OneStepTM PCR Inhibitor Removal Kit (catalog number D6031, Zymo Research, Irvine).

Across the dataset, between one and three libraries were generated per individual (e.g., SG1–SG3) and included both single and double-indexed libraries. Indexing PCR was performed using 15-18 cycles. Library yields varied substantially across tissue types and individuals, with fluorometric concentrations (Qubit) ranging from ∼0.3 ng/µl to >50 ng/µl, with the highest yields generally observed in petrous-derived samples and ossicles. Fragment length distributions were measured by capillary electrophoresis (Fragment Analyzer or TapeStation). Libraries were purified using silica-column methods (MinElute, Qiagen), yielding final volumes of approximately 35 µl, and subsequently quantified and quality-controlled prior to downstream analyses.

All laboratory work was conducted in dedicated ancient DNA facilities following strict contamination control procedures, and libraries were stored under long-term aDNA conditions at -20 °C prior to sequencing.

#### DNA sequencing

DNA was sequenced using the Illumina NextSeq 500 platform with the 75-bp single-end or 150-bp paired-end method.

### Validation of newly sequenced individuals

For the twelve newly reported individuals, we performed various analyses related to ensuring their quality and their belonging to the expected ancestral populations.

#### Read mapping to the human reference genome

Raw reads were processed to remove library adapters using AdapterRemoval (v. 2.2.2) software, and default sequencing adapters: forward adaptor: AGATCGGAAGAGCACACGTCTGAACTCCAGTCAC, reverse adaptor: AGATCGGAAGAGCGTCGTGTAGGGAAAGAGTGTA. Subsequently, reads were aligned to the Human reference genome (human_g1k_v37.fasta, hg37 with chrM) using BWA (Burrows-wheeler aligner) (v.0.1.17) *aln - samse*, with parameters: -n 0.01 -l 16500, and a mapping quality threshold of 30. Then, aligned files were merged per individual using SAMTools (v. 1.12) merge. PCR duplicates were removed using SAMTools (v1.12) *rmdup*. We kept the unmapped reads for the pathogen analysis (see section “Pathogen detection” below).

#### Genotype calling

As the samples were not UdG treated, aligned reads were trimmed off the 10 base-pairs from their ends with BamUtil (v. 1.0.13) *trimBam*, to avoid the inclusion of post-mortem damage in downstream analyses. We performed pseudohaploid genotype calling on the samples with sequenceTools (v. 1.2.1) pileupCaller (-m RandomCall option), from a pileup file created using SAMtools (v. 1.12) mplieup (-R -B -q30 -Q30). Two datasets were created by genotyping the newly reported individuals to the Genome-Wide Human Origins (HO) SNP array and the 1240k SNP. We merged our data with publicly available genomic data of ancient and present-day individuals^22^ by using EIGENSOFT package (v. 6.0.1) *mergeit* tool^138,139^. The resulting HO and 1240k datasets were filtered by removing phenotypic SNPs that are under selective pressure using the EIGENSOFT package *convertf*^138,139^. We used all these genotype calls for various sub-sections below “Haplogroup determination” (Y chromosome), “aDNA authenticity procedures of newly sequenced individuals”, “Principal Component Analysis based on SNP panel genotypes”, “F4 statistics proximal ancestral profile modeling of newly sequenced individuals”, “F4 statistics broad ancestral profile modeling of publicly available and newly reported datasets”, and “Genetic Kinship identification”. The number of SNPs genotyped in each individual can be found in **Supplementary Table 1**.

#### Haplogroup determination

Y-chromosome haplogroups were assigned by manually annotating the identified genotyped variants using the International Society of Genetic Genealogy (http://www.isogg.org, accessed on 23 October 2023) version 15.73. mtDNA haplogroups were assigned by calling a consensus sequence from the ancient mtDNA trimmed reads using iVar consensus (v. 1.9.5), and calling the haplogroups by Haplogrep 2^144^. Each haplogroup call was then manually revised by exploring the alternative variants in using IGV^145^. These results can be found in **Supplementary Table 1**.

#### aDNA authenticity procedures of newly sequenced individuals

To ensure the quality of the sequenced datasets we inferred their Post-mortem damage (PMD) profiles. Final aligned reads (merged by individual and without PCR duplicates) were assessed for aDNA PMD. We analyzed the deamination ratio at the end of the aDNA fragments using mapDamage v2.0.8. Additionally, we assessed potential contamination from human sources using ContamMix as explained in the section “Assessment of modern human contamination”.

#### Principal Component Analysis based on SNP panel genotypes

To confirm that the newly sequenced individuals belong to the expected ancestral populations, we performed a Principal Component Analysis with smartpca from EIGENSOFT v8.0.0 with lsqproject=YES and autoshrink=YES^138,139^. We computed the Principal Components (PC) using a set of 1081 present-day West-Eurasian individuals genotyped using the Human Origins array, and projected the ancient individuals (**Extended Data Fig. 3a**).

#### F4 statistics proximal ancestral profile modeling of newly sequenced individuals

To further validate that the newly sequenced individuals belong to the expected ancestral populations we calculated their F4 statistics using the module qpAdm in ADMIXTOOLS^146^ v.6.0 with ‘allsnps = YES’ parameter, to infer possible ancestry mixtures. F4 tests were performed using the ‘1240k’ SNP array genotyped dataset. The target populations were each of the newly reported individuals. The proposed model used proximal sources of ancestry and was as follows; for the outgroups we used: Mbuti.DG^147^, Russia_Kostenki14_UP.SG^148^, Russia_MA1_UP.SG^149^, Ukraine_Mesolithic.SG^26,29,150^ and Turkey_Central_Musular_PPN.SG^151^. We used as possible source population proxies: Iron_Gates_Mesolithic.SG (WHG)^26–28^, Balkan_Early_Neolithic.SG (EF)^26,28,29^, Luxembourg_Mesolithic. DG (WHG)^152^ and Turkey_Marmara_Barcin_N.SG (EF)^153^ (see **Supplementary Results**). All data produced for these populations used for this analysis was generated using whole-genome shotgun technologies to avoid statistical distortion observed when calculating f4-statistics because of generation allelic bias^154^; and genotyped to the 1240k SNP panel array. These ancestry proportions can be found in **Supplementary Table 1**.

#### Genetic Kinship identification

We determined kinship relationships among the newly reported individuals using Relationship Estimation from Ancient DNA (READ) software^155^, using the proportion of non-matching alleles (P0) and normalizing these values by the P0 median value specifically by population (either Western Hunter Gatherer (WHG) or EF), using the default window-size value of 1 Mbps.

### Sequence data retrieval from public datasets

To obtain the raw reads for the initial 201 public samples (see above) we downloaded their corresponding runs from European Nucleotide Archive (ENA) at EMBL-EBI under accession number indicated in the ‘AADR_v62.0_1240k_public.anno’ file available at https://dataverse.harvard.edu/dataset.xhtml?persistentId=doi:10.7910/DVN/FFIDCW ^22^. Then, we obtained project metadata by running ‘curl https://www.ebi.ac.uk/ena/portal/api/filereport?accession=<ena_project_code>&result=read_run&fields=<fields>&format=tsv’. These metadata tables allowed us, with some manual supervision, to obtain the links to the fastq files for each run and download them, as implemented in IGfun.get_df_AADR_with_SRR_info and IGfun.download_one_srr_reads. We discarded a few runs due to unavailable or truncated data, as described in the section “Sample filtering”. Next, we used the script get_chunks_reads.py on all datasets to generate chunks of 500,000 read pairs (or single reads for the single-end runs), for more efficient parallelization in downstream analyses. This script uses the function IGfun.get_chunks_one_fastq_file to get the chunks of reads, followed by several checks to avoid data corruption.

### F4 statistics broad ancestral profile modeling of publicly available and newly reported datasets

To validate that the analyzed samples (201 publicly available and 12 newly sequenced) belonged to the expected ancestral populations, we calculated F4 statistics for them, inferring the proportions of Early Farmer (EF), Western Hunter Gatherer (WHG), and Eastern Hunter Gatherer (EHG) ancestries. In brief, we performed F4-statistics using the qpAdm module in ADMIXTOOLS v.6.0 with ‘allsnps: YES’. Analyses were restricted to the ‘1240k’ SNP panel. For each target individual, we tested a one-way, two-way, and three-way model comprising three source populations: Russia_YuzhniyOleniyOstrov_Mesolithic.AG (EHG)^24,156,157^, Turkey_Marmara_Barcin_N.AG (EF)^24,158^, and W_Mesolithic.AG (WHG)^4,159,160^. The following set of outgroups was used: Mbuti.DG, UstIshim.DG, Kostenki14.SG, MA1.SG, Italy_Epigravettian.AG.BY.AA, and Turkey_Central_Musular_PPN.SG. These ancestry proportions can be found in **Supplementary Table 1**.

### Sample filtering

To select a final set of high-confidence samples for subsequent analyses we applied various filtering criteria on the initial set of 201 public and 12 newly sequenced individuals. Initially, we used qpAdm results (described above) to only keep samples with a clear HG (WHG + EHG), EF or admixed HG/EF ancestry. To assign each sample to the “HG”, “EF” or “admixed” ancestry we used different criteria for either public or newly-sequenced datasets. For the public datasets we used the WHG, EHG and EF proportions inferred as described in “F4 statistics broad ancestral profile modeling of publicly available and newly reported datasets”, discarding 12 samples with unknown ancestry profiles (’Chan.SG’, ‘I4434.SG’, ‘I4553.SG’, ‘I4582.SG’, ‘I5407.SG’, ‘KGH6.SG’, ‘M95.SG’, ‘R15.SG’, ‘R7.SG’, ‘SRA62.SG’ and ‘VLASA32.SG’, ‘mus005.SG’). An unknown ancestry was assigned to an individual in two cases: i) if no fitting model was yielded via qpAdm, or ii) if an ambiguous ancestry was yielded with qpAdm modelling (i.e. both models with HG only and HG+EF ancestry gave a significant fit, so we could not determine if these individuals were admixed or unadmixed). Conversely, for the newly sequenced datasets we used these same WHG, EHG and EF proportions unless the ancestry was unknown, in which case we used the ancestry proportions based on the analysis described in “F4 statistics proximal ancestral profile modeling of newly sequenced individuals”. All the ancestral population proportions were parsed and formatted with IGfun.get_df_admixture_info, resulting in the metrics shown in **Supplementary Table 1**. At the end, the qpAdm-based filtering left 189 public individuals and 12 sequenced here with assigned ancestry profiles.

Furthermore, during the downloading of public datasets (see section “Sequence data retrieval from public datasets” above) we discarded 13 samples. We ignored sequencing runs for which the ENA reads were truncated (“ERR2225780”, “ERR2060277”, “ERR5295751”, “ERR8607964”, “ERR9124863”, “ERR8656795”, “ERR4057507”, “ERR4304564”, “ERR9236593” and “ERR2225787”),resulting in the filtering out of out one sample (’SF12.SG’) which only had the truncated run (“ERR2060277”). Furthermore, we filtered out 11 samples for which the reads were not available (’I4438.SG’, ‘I4550.SG’, ‘I4439.SG’, ‘I1763.SG’, ‘I4440.SG’, ‘I4432.SG’, ‘I4595.SG’, ‘I4552.SG’, ‘I4596.SG’, ‘I1734.SG’, ‘I4551.SG’), and one that came from methylome profiling according to the ENA annotations (’I1583.SG’). This resulted in a filtered dataset of 176 public datasets (52 EF, 13 HG, 111 admixed) and 12 sequenced here.

Finally, we discarded a few samples that may have faulty sequencing data. First, we kept only those public datasets that had, according to ‘AADR_v62.0_1240k_public.anno’ (described above), i) ‘Data type’ = ‘Shotgun’, and ii) ‘ASSESSMENT’ being ‘PASS’ or ‘MERGE_PASS’. Second, we kept only datasets with up to 10% of modern human contamination (see “Assessment of modern human contamination” below) (all except five admixed individuals), measured as described below. Note that the newly-sequenced datasets always had <3.3% of contamination, validating their quality. Also, although a few (15) AADR samples had small inferred contamination levels (3-10%), we consider them as valid due to them passing the rigorous quality assessment performed by the AADR curators (related to the ‘ASSESSMENT’ field, mentioned above). Third, we filtered out the run “ERR9118893” as it had an unexpectedly high FastQC max_quality (> 60, see “Read trimming and mapping” below), which may be due to invalid quality encodings. Fourth, we discarded samples that had a mean autosomal coverage < 0.5 (see “Coverage calculations” below), including seven admixed individuals from AADR (’PB754.SG’, ‘KH150626.SG’, ‘KH150195_KH150196_KH150616.SG’, ‘PN16.SG’, ‘PB1794.SG’, ‘KH150200.SG’ and ‘KH150618.SG’), one EF from AADR (’ANY-4027.SG’), one HG from AADR (’Hum1.SG’), and one of the HG sequenced here (’i722-i513’). Fifth, we validated that all datasets have genome-wide coverage by checking that there are no outliers in the general correlation between mean autosomal coverage and percent of the autosomal genome covered (**Extended Data Fig. 4a**). These filters resulted in a final high-confidence set of 141 public datasets (48 EF, 11 HG, 82 admixed) and 11 sequenced here (152 in total), found in **Supplementary Table 1**.

Note that we used such strict sequencing data-related filters because they mostly affected admixed individuals, for which we had sufficient datasets, enabling us to remove a few of them. We used these 152 samples in most analyses below, except when specifically indicated. For instance, for the analysis of pathogens we discarded various samples due to technical reasons of the DNA source (see “Pathogen detection” below). Also, for analyses that consider admixed populations specifically (e.g. adaptive introgression scans or population-specific parameter benchmarks, referred below), we only considered individuals belonging to the homogeneous admixed neolithic populations comprised of individuals samples in present-day Germany (n = 11 / 85), Ireland (n = 32 / 85) or Portugal (n = 8 / 85) (see **Main text, Fig. 1e-g**). Throughout the article, we refer to modern political entities only as geographical descriptors, without implying continuity or determinism. Thus, in the **Main text** we refer to the admixed populations to nGer, nIre and nPor, respectively.

### Read trimming and mapping

To enable read trimming we obtained quality control information for each sequencing run and read type (forward and reverse), using the script get_adapters.py (env:perSVade_env) on six representative chunks of 500,000 raw reads per run. This script i) concatenates the read chunks into one sequencing dataset, ii) runs FastQC on the concatenated reads with the function run_fastqc of scripts/sv_functions.py (see https://github.com/Gabaldonlab/perSVade) and iii) gets the FastQC-inferred overrepresented sequences with the function get_set_adapter_fastqc_report of scripts/sv_functions.py. We obtained various relevant data from these FastQC runs, including i) the overrepresented sequences corresponding to the used adapters in each run, ii) the quality encoding, which we verified to be always ‘Sanger / Illumina 1.9’, and iii) the maximum 90th percentile per-based read quality (max_quality). Finally, we processed all these measurements to get a per-run dataset that includes i) a table with all adapters (single-end runs) or all combinations of forward / reverse adapters (paired-end runs), and ii) the max_quality. Then, we discarded runs with max_quality > 60, which may reflect invalid quality encodings (mentioned above). Note that for runs in which we found no adapters we set the default Illumina ones: ‘AGATCGGAAGAGCACACGTCTGAACTCCAGTCAC’ (forward or single-end reads) and ‘AGATCGGAAGAGCGTCGTGTAGGGAAAGAGTGTA’ (reverse reads). All this adapter inference was performed as implemented in IGfun.get_df_adapters_per_run and IGfun.get_df_adapters_per_run_collapsed.

For mapping we used the hg37 (a version of GRCh37) reference genome with chrM, obtained from http://ftp.1000genomes.ebi.ac.uk/vol1/ftp/technical/reference/human_g1k_v37.fasta.gz. Also, to enable read alignment we generated / obtained various files. First, we retrieved the .fai index file from this same ftp site from which we got the reference sequence. Second, we used picard CreateSequenceDictionary (env:perSVade_env_picard_env) to get sequence dictionaries from the genome fasta file. Third, we ran bwa index (env:perSVade_env) to index the genome for bwa-based mapping.

Then, for each chunk of 500,000 raw reads for a given sample, we ran the script map_reads.py to trim and map the reads to the hg37 reference. This script runs the eager nextflow pipeline (env:eager_env) to i) remove adapters and collapse paired-end reads into one with AdapterRemoval, ii) read mapping with *bwa aln* (arguments -mapper bwaaln --bwaalnn 0.01 --bwaalnl 16500 --bwaalno 2) and iii) filtering out reads with mapping quality < 30 (arguments --run_bam_filtering --bam_mapping_quality_threshold 30). We used these mapping parameters as they have been widely used in aDNA studies^161,162^. Also, note that we tailored the AdapterRemoval arguments to each run, passing the adapters table (defined above) to --clip_adapters_list, and setting --qualitymax depending on max_quality (i.e. ‘--qualitymax 61’ if max_quality = 60; ‘--qualitymax 41’ if max_quality <= 41). This resulted in one coordinate-sorted filtered bam file for each chunk of reads, as well as the raw reads collapsed into single-end format (for paired-end runs) with adapters trimmed, as generated by AdapterRemoval.

Next, we integrated these bam files of different chunks using the script merge_chunks_bams.py, resulting in one bam for each chromosome in each sample. In brief, this script processes all bam files coming from different chunks in a given sample, using samtools commands (view, index and sort, on env:eager_env) to get the reads mapping to a given chromosome. This allowed us to process individual chromosomes in parallel in downstream analyses. The pipeline to obtain these mapped reads per chromosome is in IGfun.add_cmds_trim_and_map_reads.

### Assessment of modern human contamination

#### Calculation of the percent of modern human contamination

To validate the quality of the datasets we used ContamMix to assess the fraction of modern human contamination in all the samples analyzed here, processing the reads mapped to the mitochondrial (MT) chromosome with the script run_contammix.py on each sample. This script first runs the eager nextflow pipeline (env:eager_env) on the MT-mapped and filtered reads (argument --bam) to i) remove read duplicates with picard MarkDuplicates (argument --dedupper markduplicates), and ii) trim 10 bp from each read end with bamutils (arguments --run_trim_bam --bamutils_clip_double_stranded_none_udg_left 10 --bamutils_clip_double_stranded_none_udg_right 10 --bamutils_softclip). Also, it runs IGfun.generate_reads_fastqgz_from_bam_single_end on these ‘dedupped trimmed bam’ to obtain the corresponding ‘dedupped trimmed reads’.

Next, run_contammix.py generates a consensus mitochondrial sequence with ‘samtools mpileup’ (with -B -a) and ‘ivar consensus’ (with -t 0.6 -m 5 -q 20) (env:contammix_env) on the ‘dedupped trimmed bam’. Then, it uses mafft (env:contammix_env) to generate a Multiple Sequence Alignment (MSA) of this consensus and 313 potential mitochondrial contaminants. This list of potential contaminants include i) 311 sequences used in the ContamMix publication^78^, and ii) two sequences from the present-day DNA handlers (undisclosed for privacy). Further on, this script re-aligns the ‘dedupped trimmed reads’ to the mitochondrial consensus. Finally, it runs contammix with argument --tabOutput (env:contammix_env) using the re-aligned reads and the generated MSA, which results in the contamination estimates. The code to run this script on all samples is in IGfun.get_Contammix_results. Note that we took the ContamMix metric “2.5% authentic” as the authenticity rate, and thus we calculated the percentage of modern human contamination as (1 - ‘2.5% authentic’) * 100. Then, we kept only datasets with up to 10% of such modern human contamination (as described above).

#### Establishing a maximum read length threshold to avoid modern human reads

To further verify that the analyzed sequences belong to endogenous ancient DNA we evaluated the length distribution of reads mapped to chromosome 19, as bimodal length distributions may reflect a mix of ancient reads (shorter) and modern ones (longer)^133^. For this, we ran the eager nextflow pipeline (env:eager_env) on the chromosome 19 filtered bam (argument --bam) to remove read duplicates with picard MarkDuplicates (argument --dedupper markduplicates), and then used IGfun.generate_readlen_distribution_file to infer the read length distribution based on the de-dupped bam. The read length distribution of several AADR samples was bimodal (**Extended Data Fig. 2**), which could reflect some degree of contamination. We found particularly concerning the read length peaks at >100 bp observed in some samples as potentially representing modern DNA unfragmented molecules (**Extended Data Fig. 2**), so we discarded reads longer than 90 bp in most analyses (mentioned below) to ensure high-confidence results. Although such bimodal distributions could also be related to read-collapsing artifacts of paired-end aDNA sequencing, something that is plausible given the low inferred percentages of contamination (see above), we still decided to discard long reads (> 90 bp) to minimize the risk of including modern sequences in our analysis.

### Coverage calculations

To enable Copy Number Variant (CNV) analysis, sex inference, and assessment of average coverage statistics we performed various per-window coverage calculations. For this, we first processed the filtered and mapped reads for each chromosome in each sample (generated with IGfun.add_cmds_trim_and_map_reads as described above) with process_bams_for_varcall.py. This is a pipeline using IGfun.generate_bam_one_chrom_dedupped_and_long_reads_corrected to i) discard reads longer than 90 bp with samtools and awk (env:eager_env) (could be modern human contaminants) and ii) remove duplicates with eager (argument --dedupper markduplicates) (env:eager_env).

For various window sizes (1 kb for mtDNA, and 10 kb, 20 kb, 50 kb or 100 kb for gDNA chromosomes) we ran perSVade’s call_CNVs module (env:perSVade_env) on these processed bams to obtain per-window mean coverage estimates. Specifically, we used arguments ‘-p 2 --window_size_CNVcalling <window size, 1kb - 100kb> --skip_coverage_correction --mappability_file <mappability file> --average_cov_measure mean --max_fraction_N_bases 0.1 --max_fraction_repeats 0.1 --repeats_file <repeatmasker repeats> --min_median_mappability 0.9 --skip_CNV_calling’. We generated the per-position mappability file with get_mappability_file_whole_genome.py (env:perSVade_env), which runs perSVade’s function generate_genome_mappability_file (see scripts/sv_functions.py at https://github.com/Gabaldonlab/perSVade). Also, the repeats used were obtained running RepeatMasker with arguments ‘-poly -noint’ (env:perSVade_env_RepeatMasker_env), as implemented in IGfun.add_cmds_get_repeats_per_chrom. This generated mean coverage estimates for all windows with median mappability >= 0.9, fraction repeats <= 0.1 and fraction Ns <= 0.1, referred to as “mappable windows”, available in the file ‘final_df_coverage.tab’. For reference, the filtered 100 kb (gDNA) windows cover 86.87% of the total length of all autosomes.

On the one hand, we used the 100 kb gDNA measurements to infer various per-sample statistics, as implemented in IGfun.get_df_coverage_stats. For instance, we calculated the mean autosomal coverage for each sample, used for dataset filtering, following the expression *mean coverage = ∑ (mean coverage w_i_ * length w_i_ / autosomal length)*. Here, *w_i_* refers to each of the windows, and *autosomal length* is the sum of the length of all filtered windows for autosomes. Similarly, we calculated the percentage of the positions in these autosomal filtered windows that was covered by at least one read, useful for validating that all datasets have genome-wide coverage (**Extended Data Fig. 4a**). Also, we used these calculations to infer the sex of each individual based on the relative coverage of X / Y chromosomes. Specifically, we defined the relative coverage of X / Y chromosomes as the mean coverage of each sexual chromosome (across windows) normalized by the mean autosomal coverage. Then, we identified as females the samples that had a X relative coverage >= 0.8 and a Y relative coverage <= 0.1. Conversely, we identified as males the samples with relative coverage between 0.3-0.7 for both X and Y. Also, we found a sample with 0.79 relative coverage X, and 0.0 in Y, which we classified as female. On the other hand, we used the 10 kb-100 kb gDNA results for CNV analysis, as described below.

### Generation of raw SNP calls

To call SNPs in each dataset dataset we ran the script process_bams_and_call_variants_per_window.py in parallel across 1 Mbp windows of the genome, defined with ‘bedtools makewindows’ on the hg37 reference genome (env:perSVade_env). In brief, this is a pipeline that first extracts the target 1 Mbp region from the filtered bam, using samtools (env:eager_env). Then, it removes reads with >90 bp, which could be modern human contaminants, as defined above.

After this, our script runs eager (env:eager_env) on the read length-filtered bam to i) remove read duplicates with picard MarkDuplicates (argument --dedupper markduplicates), ii) infer PMD profiles with MapDamage (arguments ‘--damage_calculation_tool mapdamage --mapdamage_downsample 1000000’, only for chromosome 19), and iii) trimming 10 bp from each read end with bamutils, which are set to N (arguments --run_trim_bam --bamutils_clip_double_stranded_none_udg_left --bamutils_clip_double_stranded_none_udg_right 10). Furthermore, our script corrects the resulting bams to ensure properly set read groups, using picard AddOrReplaceReadGroups (env:eager_env). This generates a processed bam used for further SNP calling. Also, we used the calculated PMD profiles for the validation of simulations, as described below.

Regarding SNP calling, to enable benchmarking of different algorithms and parameters, the script process_bams_and_call_variants_per_window.py runs SNP calling on the processed bam using five algorithms. These include i) GATK HaplotypeCaller (HC) (env:perSVade_env), ii) bcftools call (bt) (env:perSVade_env), iii) freebayes (fb) (env:perSVade_env), iv) freebayes in a pooled ploidy-agnostic mode (pfb) (env:perSVade_env) and v) AriaDNA (AR) (env:ariadna_env), a machine learning-based pipeline thought for aDNA SNP calling^134^. The first three algorithms (HC, bt, fb) are run by executing perSVade’s call_small_variants module^89^ with arguments ‘-p <calling ploidy, 1 or 2> --callers HaplotypeCaller,bcftools,freebayes -c 0 --min_AF 0.0’. Similarly, the pfb algorithm is run by executing the same module, with arguments ‘-p <calling ploidy, 1 or 2> --callers freebayes -c 0 --min_AF 0.0 --pooled_sequencing’. Finally, the AR algorithm is run by executing the ARIADNA.py script from their repository in https://osf.io/5bph4/. Note that the calling ploidy for perSVade runs was set to 1 for chromosomes MT, Y and X (the latter only for males), and to 2 for all others. After running these algorithms, the script integrates all of the identified raw variants and their calling statistics (e.g. quality or allele coverage) into the file merged_variants.tab.

### Sample-tailored simulations of genomic datasets

#### Generation of simulated genomes

To enable parameter benchmarking for comparative genomic analyses, we generated simulated reads resembling each of the input individuals. For this, we first generated three types of genomes (in fasta format) to enable realistic simulations: ancient human genomes (resembling the ones we wanted to analyze), modern human genomes (contaminants) and microbial genomes (contaminants). For the ancient genomes, to get realistic results we simulated three diploid genomes based on chromosome 19, each with a set of SNPs corresponding to individuals “VLASA7” (western HG), “KK1” (caucasian HG) or “LEPE48” (EF), defined in ^28^. We chose these three samples because they represent individuals from the Mesolithic-Neolithic period, each including highly-distinct SNP patterns (see Figure 1A from ^28^).

Specifically, we first obtained the phased SNP calls from ftp://ftp.sra.ebi.ac.uk/vol1/ERZ128/ERZ12801402/Ancient10X_08-2021_mergeAll102Labelled2_phasedREF_Allchr.vcf.gz, based on the reference genome hs37d5, which we checked to be equivalent to the used hg37. Also, for fast access we obtained the corresponding index file from ftp://ftp.sra.ebi.ac.uk/vol1/ERZ128/ERZ12801402/Ancient10X_08-2021_mergeAll102Labelled2_phasedREF_Allchr.vcf.csi. Then, we used bcftools view (env:eager_env) to keep only the chromosome 19 SNPs. Finally, we ran the function IGfun.get_genome_dir_with_variants to simulate the genomes bearing the SNPs of each ancient individual (“VLASA7”, “KK1” or “LEPE48”). This function uses gatk FastaAlternateReferenceMaker (env:perSVade_env) to generate two haploid chromosome 19 sequences, each with the father/mother set of SNPs of the given simulated individual.

To generate a modern human genome contaminant we simulated the SNPs from “Mende-1”, also provided by ^28^, which was always the one with the largest genetic distance to the ancient samples (“VLASA7”, “KK1” and “LEPE48”, see Supplementary Table S2 from ^28^) We decided to use this farthest sample to simulate a human contamination that was as difficult to handle as possible. Specifically, we ran the function IGfun.get_genome_dir_with_variants, explained above, to simulate the genome bearing the “Mende-1” SNPs on chromosome 19.

Finally, to simulate a realistic microbial contaminant community we used the set of genomes found in bactDBexample/k14 community from the gargammel (aDNA simulator) repository https://github.com/grenaud/gargammel. This was also used in ^163^. To generate these files we downloaded gargammel’s repository release v1.1.4, and ran ‘make bacterialex’ (env:gargammel_env). The generation of these simulated genomes is implemented in IGfun.get_simulated_genomes_for_gargammel.

#### Generation of simulated reads

To generate realistic simulated reads from these simulated genomes we used the script simulate_reads_gargammel.py on all samples. It generates three sets of single-end reads, one for each ancient individual (“VLASA7”, “KK1” or “LEPE48”), with a certain fraction of ancient (endogenous), modern human (contaminant) and bacterial (contaminant) DNA. To ensure sample-tailored simulations we used a coverage, read length, PMD profile and fraction of contaminant reads matching those empirically calculated for each sample. Specifically, we defined the fraction of modern human contaminant reads (*f_cont*) as:

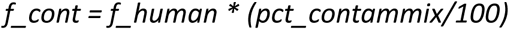

Where *f_human* is the fraction of reads mapped to the human reference genome, and *pct_contammix* is the percentage of these reads empirically predicted to be modern human contaminants (see section “Assessment of modern human contamination” above). Specifically, we defined *pct_contammix* as (1 - ‘2.5% authentic’) * 100. Also, we used IGfun.get_fraction_human_samples_chunks_bams to calculate *f_human* in these samples, which divides the number of reads in the filtered bam by the total number of raw reads. Based on these numbers, for each sample we also defined the fraction of bacterial reads (*f_bact = 1 - f_human*) and the fraction of endogenous reads (*f_endo = f_human - f_cont*). All of these were used by simulate_reads_gargammel.py to generate sample-tailored simulations.

This script first generates various calculations by running the eager nextflow pipeline (env:eager_env) on the chromosome 19 filtered bam (generated as explained in “Read trimming and mapping” above) (argument --bam) to i) remove read duplicates with picard MarkDuplicates (argument --dedupper markduplicates), ii) calculate coverage per 500 kbp windows on de-dupped bam with bedtools (arguments ‘--run_bedtools_coverage --anno_file <windows bed>’) and iii) infer PMD profiles with MapDamage (arguments ‘--damage_calculation_tool mapdamage --mapdamage_downsample 1000000’). Then, it uses the function IGfun.generate_readlen_distribution_file to infer the read length distribution based on the de-dupped bam. Next, it calculates the total number of human fragments to simulate so that the final coverage of the simulated reads after mapping, bam filtering and de-duplication is equivalent to the real one. Specifically, this number of human fragments is defined as the total number of reads in the de-dupped bam (calculated with IGfun.get_n_reads_bam) divided by the filtered/mapped ratio in the original bams (calculated with IGfun.get_fraction_reads_filtered_vs_mapped). Then, it sets the total number of fragments to simulate as the number of human fragments divided by *(f_endo + f_cont)*.

The final step in simulate_reads_gargammel.py is, for each simulated ancient individual (“VLASA7”, “KK1” or “LEPE48”), the generation of the simulated reads using IGfun.run_gargammel_inHouse based on the empirically calculated PMD profile, read length distribution and total number of fragments. In brief, this pipeline uses fragSim (simulator of fragments), deamSim (simulator of PMD), adptSim (simulator of adaptors) and art_illumina (read simulator), all ran on env:gargammel_env, to generate sample-tailored simulated reads, each with a particular mix of genomes according to *f_endo, f_cont* and *f_bact*. Given that we generated single-end reads, in adptSim we simulated the default Illumina ‘AGATCGGAAGAGCACACGTCTGAACTCCAGTCAC’ adapter. Also, note that we developed this custom pipeline due to unsolvable errors when running the gargammel.pl wrapper provided in https://github.com/grenaud/gargammel.

In summary, for each analyzed sample we generated three sets of single-end chromosome 19 simulated reads (one for each “VLASA7”, “KK1” or “LEPE48” genomes), each with sample-tailored technical properties and contamination profiles. These simulations were performed as implemented in IGfun.add_cmds_read_simulation_gargammel_one_chrom.

#### Generation of processed genomic datasets for simulated reads

To enable parameter benchmarking we processed these simulated reads in a way that was equivalent to the real ones. First, we mapped the reads as described in “Read trimming and mapping”, using the simulated default Illumina adapter for trimming. Second, we calculated coverage statistics as mentioned in “Coverage calculations”. Third, we generated raw SNP calls and other processed datasets as indicated in “Generation of raw SNP calls”. Fourth, we imputed variants for these datasets as explained in “SNP imputation”.

#### Validation of simulations

To validate that the simulated datasets resemble their corresponding real sequencing data we compared various statistics of the simulated and real datasets. For instance, we considered their mean autosomal coverage and percent of the autosomal genome covered (**Extended Data Fig. 4b**), calculated as described above. Also, we used IGfun.get_df_read_lens to calculate the read length distribution for all samples (real and simulated) for the de-dupped and trimmed bam of the first five “mappable” 1 Mb windows of chromosome 19, relying on function IGfun.generate_readlen_distribution_file (described above). These include windows with median mappability >= 0.9, fraction repeats <= 0.1 and fraction Ns <= 0.1, inferred with perSVade’s call_CNVs as described in “Coverage calculations” above. The read length distributions can be found in **Extended Data Fig. 5**. Finally, we ran IGfun.get_df_mapdamages to process the PMD profiles, calculated from the de-dupped bam as explained in “Generation of raw SNP calls” above, for the same five “mappable” 1 Mb windows of chromosome 19. Specifically, we recorded the content of the MapDamage output ‘5pCtoT_freq.txt’, containing the 5’ C>T mutation rate for the first and last 25 bps of reads (see **Extended Data Fig. 6**). Overall, coverage, read length and PMD metrics were highly similar between real and simulated samples (**Extended Data Fig. 4b,5,6**), validating our simulation pipeline.

### SNP calling parameter exploration

To evaluate whether SNP calling is feasible in our dataset, we analyzed the SNP calling performance of various algorithms and filtering parameters across the three simulated datasets (“VLASA7”, “KK1” and “LEPE48”) resembling each of the real samples. For these simulations we knew the set of true SNPs, so we could use the called SNPs to find optimal calling strategies. Also, this benchmarking allowed us to infer the calling performance in real datasets, which was essential to build trust in the obtained results. Specifically, for each simulation, we measured the calling performance of various callers / filters across samples when varying i) the minimum mean autosomal coverage (defined above) that a sample should have to be considered (‘cov_s’, set to 0, 1, 2, 3), ii) the set of samples considered (‘sample_set’, all datasets or various downsampled datasets, see **Supplementary Information**, **Extended Data Fig. 8a,9**), and iii) the minimum coverage that a position should have in all considered samples (depending on cov_s and sample_set) to be considered (‘cov_p’, set to 1, 2, 3, 4, 5). This benchmarking was performed as implemented in IGfun.add_cmds_varcall_parameter_optimization.

To be able to use cov_p we calculated the coverage per position in all simulated and real datasets with the script get_cov_per_pos.py, applied to each of the 1 Mbp-long dedupped and trimmed bam files. This pipeline i) clips out the 10 bp trimmed from each read end set to Ns, using samtools and awk (env:eager_env) and ii) calculates the coverage per position on this clipped bam with mosdepth (env:perSVade_env).

For each combination of ‘cov_s’, ‘cov_p’ and ‘sample_set’ we initially calculated the estimated percentage of the genome covered. First, we defined a collection of representative “mappable” 1 Mb autosomal windows, inferred as described in “Validation of simulations” above, taking 5 randomly chosen windows per chromosome. Second, we ran get_fraction_genome_cov.py to infer the percentage of covered windows, as a proxy the whole-genome coverage. This script relies on IGfun.get_file_positions_w_sufficient_coverage to keep positions with coverage >= cov_p in all considered samples in these representative windows, using the coverage results generated with get_cov_per_pos.py. The resulting coverage estimates are shown in **Extended Data Fig. 8a**.

Then, for combinations of ‘cov_s’, ‘cov_p’ and ‘sample_set’ with >5% of such genome coverage we calculated the precision, recall and F-value (harmonic mean between precision and recall) for finding simulated SNPs when using different combinations algorithms / parameters. For this we ran run_parameter_optimization.py on each ‘cov_s’, ‘cov_p’ and ‘sample_set’, to benchmark combinations of i) algorithms required (HC, bt, fb, pfb and/or AR) ii) minimum number of reads covering the variant (1 to 4), iii) whether to use or not some commonly used “best practices” for filtering HC and/or fb outputs^89^, and iv) different variant quality thresholds (QUAL) for each algorithm (from 0 to 50). For each parameter set (695,592 in total) this script calculates performance metrics (precision, recall, F-value), defining optimal parameters for a given simulation as those that i) have ‘F-value’ > t (t=0.1 if cov_s<=2; t=0.4 if cov_s=3) for all considered simulations, and ii) are most conservative but have an F-value >= (maximum F-value per simulation * 0.99). We designed this strategy to avoid overfitting that would be expected if simply choosing the parameter set with the highest F-value for a given simulation. Furthermore, the script calculates the performance metrics of each of these per-simulation optimal parameters on all other simulations.

Finally, to infer the SNP calling performance that can be achieved for a given combination of cov_p, cov_s and sample_set, we integrated the F-value results obtained for these per-simulation optimal parameters. First, for each such parameter set we calculated the minimum F-value obtained across considered simulations of the given cov_p, cov_s and sample_set. Then, we considered as the final SNP calling performance (for a given cov_p, cov_s and sample_set) the maximum of these minimum F-value measurements, which is shown in **Extended Data Fig. 8b,c**.

On another line, to investigate whether mean coverage per sample is the main determinant of SNP calling performance, we repeated all this benchmarking when only applying get_cov_per_pos.py to the simulations of each of the single samples, as presented in **Extended Data Fig. 7**.

### SNP imputation

Given that we could not reliably call SNPs directly, we imputed them based on a panel of 2504 unrelated individuals from the Phase 3 of the 1000 Genome project^30^, following a strategy similar to previous aDNA studies^164^. In the following sections we describe how we obtained this panel and used it to impute SNPs and run a PCA for all samples.

#### Generation of imputation panel

To generate the filtered imputation panel we first obtained the list of 2504 unrelated individuals from Phase 3 of the 1000 Genome project. For this, we obtained the full sample list from http://ftp.1000genomes.ebi.ac.uk/vol1/ftp/data_collections/1000G_2504_high_coverage/working/1kGP.3202_samples.pedigree_info.txt, and then removed the 698 individuals belonging to trios, obtained from the ENA run info for PRJEB36890^165^. Then, we obtained a filtered reference panel including all autosomes and chromosome X using the following steps. First, we obtained the raw phased SNPs from http://ftp.1000genomes.ebi.ac.uk/vol1/ftp/data_collections/1000G_2504_high_coverage/working/20201028_3202_phased/<varsfile>, where <vars file> was either ‘CCDG_14151_B01_GRM_WGS_2020-08-05_chr

<chrom>.filtered.shapeit2-duohmm-phased.vcf.gz’ (for autosomes) or ‘CCDG_14151_B01_GRM_WGS_2020-08-05_chrX.filtered.eagle2-phased.v2.vcf.gz’ (for chromosome X). Second, we used various bash commands to keep only SNPs for the 2504 unrelated individuals. Third, since these SNPs were in GRCh38 coordinates, we lifted them to GRCh37 ones matching our used reference genome with ‘gatk LiftoverVcf -C GRCh38_to_GRCh37.chain.gz --WARN_ON_MISSING_CONTIG true --WRITE_ORIGINAL_ALLELES true --WRITE_ORIGINAL_POSITION true’ (env:perSVade_env). Note that we obtained the chain file from https://ftp.ensembl.org/pub/release-111/assembly_chain/homo_sapiens/GRCh38_to_GRCh37.chain.gz.

Fourth, to keep high-confidence SNPs only we only kept those that had a ‘PASS’ filter in gnomAD^166^ v2.1.1. For the latter, we obtained the raw gnomAD GRCh37 SNPs from https://storage.googleapis.com/gcp-public-data--gnomad/release/2.1.1/vcf/genomes/gnomad.genomes.r2.1.1.sites.<chrom>.vcf.bgz, and then used various bcftools commands to keep the final list of filtered ‘PASS’ SNPs (‘bcftools annotate -c FILTER,INFO/gnomAD_FILTER:=FILTER’ and ‘bcftools filter -i ‘FILTER==\“PASS\“’’) (env:eager_env). All the code to reproduce this panel generation can be found in IGfun.generate_filtered_panel_1000G_2504_high_coverage_lifted_to_GRCh37. Furthermore, to run imputation separately for chromosome X pseudo-autosomal regions (PAR), we used bcftools (env:eager_env) to split the chromosome X filtered panel into the separate panels for regions ‘X_PAR1’ (X:60001-2699520) ‘X_PAR2’ (X:154931044-155260560) and ‘X_nonPAR (X:2699521-154931043)’; following the UCSC annotations (https://genome-euro.ucsc.edu/cgi-bin/hgGateway?hgsid=418968119_QAg0sDMDXISREg5iRafalttBi5eX). All the code to generate the imputation panel is in IGfun.add_cmds_variant_imputation.

#### SNP imputation pipeline

To run SNP imputation we ran run_imputation.py on each chromosome region (each autosome, X_PAR1, X_PAR2 and X_nonPAR), for each sample. In brief, this script takes i) the quality-filtered mapped reads with no de-duplication nor bam trimming (generated as described in ‘Read trimming and mapping’ above), ii) the filtered imputation panel for the chromosomal region, iii) the imputation ploidy (set to 1 for male chromosome X regions, 2 for all others), and iv) whether the imputation should be run on chromosomal chunks (set to True for all regions except X_PAR1, X_PAR2). Most importantly, this script performs the following steps. First, unless chunking is disabled (i.e. for X_PAR1, X_PAR2), it runs GLIMPSE2_chunk (on the glimpse_2.0.0.sif image, described above) to generate chunks of the chromosome that can be imputed separately. This required a genetic map file with recombination rate estimations, obtained from the GLIMPSE v2.0.0 release (https://github.com/odelaneau/GLIMPSE/archive/refs/tags/v2.0.0.tar.gz , file GLIMPSE-2.0.0/maps/genetic_maps.b37/chr

<chrom>.b37.gmap.gz). Note that the chromosome X map file did not include the X_PAR1 and X_PAR2 regions, which is why we did not run imputation in chunks on them.

Second, for each chromosomal chunk, run_imputation.py extracts the reads mapped to the chunk’s region using IGfun.get_bamfile_region (based on samtools (env:eager_env)), and generates a processed bam with removed read duplicates, no reads longer than 90 bp, and trimming of 10 bp from each read end. This processed bam is generated with IGfun.generate_bam_one_chrom_dedupped_and_trimmed_and_long_reads_corrected, which is equivalent to IGfun.generate_bam_one_chrom_dedupped_and_long_reads_corrected (defined above), but additionally includes bam trimming using eager with arguments ‘--run_trim_bam --bamutils_clip_double_stranded_none_udg_left 10 --bamutils_clip_double_stranded_none_udg_right 10’ (env:eager_env). Third, for each chunk, our script runs GLIMPSE2_split_reference (on glimpse_2.0.0.sif) with the argument ‘--keep-monomorphic-ref-sites’, resulting in the reference panel matching the chunk in binary format.

Fourth, taking the input chromosomal ploidy (i.e. 1 for males’ chromosome X, 2 for the others), the reference panel and processed reads for each chunk, run_imputation.py imputes the SNPs with GLIMPSE2_phase (on glimpse_2.0.0.sif) with the argument ‘--keep-monomorphic-ref-sites’. Fifth, if the ploidy was 1, for consistency with downstream analyses our script uses IGfun.generate_diploid_like_bcf_after_GLIMPSE2_phase, based on bcftools (env:eager_env), to re-format the GT values of imputed SNPs to be either 1|1 or 0|0, instead of 1 or 0, respectively, for compatibility with downstream PCAdapt analysis. Sixth, run_imputation.py integrates all the imputed SNPs into one with GLIMPSE2_ligate (on glimpse_2.0.0.sif), and converts them to a final SNP table called ‘imputed_variants_all.tab’. These imputed SNPs are the ones used for all downstream analyses referred below. All the code to run imputation is found in IGfun.add_cmds_variant_imputation.

#### PCA reconstruction based on imputed SNPs

To run a Principal Component Analysis (PCA) for all samples (**Fig. 1e**) or only admixed individuals (**Fig. 1g**) based on these imputed SNPs, we first generated an integrated dataset of biallelic imputed SNPs for various sets of samples (all or admixed). For each sample set we ran the python-based function IGfun.get_filtered_imputed_vars_one_chromosome for each chromosome region, which integrates the ‘imputed_variants_all.tab’ results and filters out non-biallelic positions within that sample set. Then, to integrate the SNPs for each sample set we ran IGfun.generate_df_vars_imputation_file_various_chromosomes_all_samples on the integrated variants of all autosomes, ignoring X chromosome regions for this analysis as we were only interested in the global population structure. Next, to run the PCA analysis on each of these integrated variants we ran IGfun.run_pcadapt with method_LD_thinning = “thinning_size_500_thr_0.1” on each sample set.

With the set parameters, this function first uses vcftools and plink to format the variants as .bed, .bim and .fam files, as implemented in IGfun.get_plink_files_from_biallelic_filtered_snps_df. Then, it runs PCAdapt, as implemented in run_pcadapt.R (env:IronGates_R_env), with K=<number of individuals> - 1, min.maf=0.05, ploidy=2, method=“componentwise”, LD.clumping=list(size=500, thr=0.1). This generates PCA results and component-wise p values for each SNP with MAF>=0.05. Finally, IGfun.run_pcadapt uses custom python code to i) calculate the fraction of variance explained by each PC as *v*^2^, where *v* is the PC singular value, ii) correct the p values with bonferroni (statsmodels.stats.multitest.multipletests), and iii) integrate and check the PCAdapt output files. We used these PCA results to infer the global population structure of our dataset ( **Fig. 1e,g**).

### SNP-based selection scans

In the sections below we explain how we performed the different selection scans (PCAdapt, XP-EHH, Fadm-LAD and iDAT) as well as how we compared them to previous similar studies of selection during the Mesolithic-Neolithic transition.

#### PCAdapt scan

To identify SNPs under differential selective pressures between the ancestral populations, we ran PCAdapt to find those driving the separation between EF and HG. For this, we integrated the imputed SNPs of all chromosomal regions of each of the EF / HG sample sets with the functions IGfun.get_filtered_imputed_vars_one_chromosome and IGfun.generate_df_vars_imputation_file_various_chromosomes_all_samples (defined above). Then, we ran PCAdapt on these filtered biallelic variants as explained in ‘PCA reconstruction based on imputed SNPs’ above, i.e. using IGfun.run_pcadapt with method_LD_thinning = “thinning_size_500_thr_0.1”. This generated files with the PCA information and PC1-related p values, where SNPs with low p values drive the separation of HG / EF along PC1 (**Fig. 2a**). We considered 6,008,635 SNPs that had MAF>=0.05 across the whole set of EF and HG individuals.

Out of these, we define as the significant ‘outlier’ SNPs those with the 300 lowest p values (p < 4.42·10^−77^), as done in a previous similar analysis^14^. We chose this threshold after trying various ones (passing bonferroni correction, lowest 1000 and lowest 300), and verifying that the ‘lowest 300’ was appropriate to single out true outliers (**Fig. 2c**). Also, to focus on independent positive selection events we identified clusters of at least 3 significant SNPs within 100kb of each other, and we kept one “leading SNP” per cluster, with lowest p-value. These “leading SNPs”, 8 in total, are the PCAdapt top hits mentioned through this work. The code to run the PCAdapt scan and define significant SNPs is in IGfun.run_SNP_selection_analyses, and the one for defining the top hits in IGfun.get_unique_SNPselection_hits.

#### Cross-population Extended Haplotype Homozygosity (XP-EHH) scan

To identify differential selective pressures between the ancestral HG / EF populations with a haplotype-based method we used the Cross-population Extended Haplotype Homozygosity (XP-EHH) method, as implemented in the rehh R package. For the autosomal variants, we used the same ones as for the PCAdapt scan (defined above). Conversely, to enable proper haplotype analyses for chromosome X variants we processed the imputed SNPs so that male (haploid) chromosome X variants have 1 or 0 GT values instead of 1|1 or 0|0 as yielded by run_imputation.py. For this, we ran the python-based function IGfun.generate_integrated_chrom_X_datasets, which integrates the variants of all chromosomal regions (X_PAR1, X_PAR2, X_nonPAR) for different sets of individuals (i.e. EF or HG) using IGfung.generate_df_vars_imputation_file_various_chromosomes_all_samples (defined above), and then re-defines the GT values for males to be 0 or 1.

To perform the XP-EHH scan we ran IGfun.run_XP_EHH_one_chromosome_set. This function first converts the processed SNPs mentioned above (for autosomes and chromosome X) to multi-sample .vcf files with IGfun.get_vcf_from_sns_df_for_plink for each EF / HG population, and then adds the population’s AF with ‘bcftools +fill-tags’ (env:eager_env) as implemented in IGfun.generate_vcf_with_AFinfo. Next, it runs run_XPEHH.R (env:IronGates_R_env) on these .vcf files to get the per-SNP XP-EHH calculation. In brief, run_XPEHH.R i) loads the variants of each EF / HG population with data2haplohh (rehh package), ii) filters them to keep those with intra-population MAF >= 0.05 with subset (rehh), iii) calculates the integrated site-specific Extended Haplotype Homozygosity (iES) with scan_hh (rehh), and iv) performs the XP-EHH scan with ies2xpehh (rehh) taking the EF / HG iES results as ‘target’ and ‘background’, respectively. This generated the XP-EHH ratio per position, where positive values reflect higher iES in EF (and *vice versa* for negative values), and its associated p-value. After the run_XPEHH.R run, IGfun.run_XP_EHH_one_chromosome_set also calculates the bonferroni-corrected p values with statsmodels.stats.multitest.multipletests. We could calculate the XP-EHH values for 4,491,626 sites, which are a subset of those where PCAdapt was run as we filtered out sites with MAF<0.05 in any of the populations, and we had to discard positions near the chromosome ends where iES could not be calculated due to missing haplotype information.

Out of these, we defined as significant sites those passing the bonferroni correction with a p<0.05 threshold. Also, to focus on independent positive selection events we identified clusters of at least 3 significant SNPs with the same selection direction (XP-EHH>0 or XP-EHH<0) within 100kb of each other, and we kept one “leading” site per cluster, with lowest p-value. These “leading sites”, 20 in total, are the XP-EHH top hits mentioned through this work. The code to run the XP-EHH scan and define significant positions is in IGfun.run_SNP_selection_analyses, and the one for defining the top hits in IGfun.get_unique_SNPselection_hits.

#### Fadm scans

To identify SNPs with an unexpectedly high or low AF in late Neolithic admixed populations (from present-day Ireland, Germany, Portugal, referred to as nIre, nGer, and nPor, respectively, in the **Main text**) as compared to the expectation from the genome-wide HG / EF mixture model, we calculated per-SNP Fadm statistics as performed in previous work on ancient selection^13,24^. For each admixed population, we first integrated the biallelic imputed SNPs of all chromosomal regions of each of the relevant sample sets (HG, EF or the admixed population), using IGfun.get_filtered_imputed_vars_one_chromosome and IGfun.generate_integrated_chrom_X_datasets (defined above). This resulted in a set of haplotype-corrected SNPs (i.e. with male chromosome X variants having 0 or 1 GT) for each sample and chromosome. Also, we defined the ancestral EF proportion of the population (pEF) as the average EF proportion across all individuals of that population, calculated with qpAdm as explained in ‘F4 statistics broad ancestral profile modeling of publicly available and newly reported datasets’ above.

Next, we ran IGfun.run_Fadm to get the per-SNP Fadm and associated p-value calculations. This function integrates all haplotype-corrected SNPs into a multi-sample .vcf file with AF per population measurements, generated as for the XP-EHH scan (described above). Then, it processes this vcf with python to generate the ‘vars_for_Fadm.tab’ table, with the AF and MAF of each biallelic variant site in each population (AF_HG_, AF_EF_, AF_ADMIXED_, MAF_HG_, MAF_EF_, MAF_ADMIXED_), including only variants with MAF MAF_HG_>0.05 and MAF_EF_>0.05. Also, for empirical p-value calculations it generates a ‘null’ subset of these variants called ‘vars_for_Fadm_null.tab’, including one SNP every 0.5 Mb. Further on, IGfun.run_Fadm calculates Fadm and related raw p-values by running calculate_Fadm_stats.R (env:IronGates_R_env) on these two .tab files. This script calculates, for each SNP, the genome-wide expected AF in the admixed population under neutrality (AF_EXP_ADMIXED_) as:

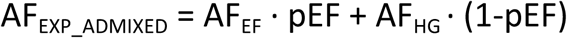

Then, based on AF_EXP_ADMIXED_ and AF_ADMIXED_ it calculates Fadm as:

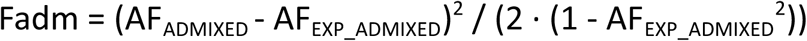

We defined the Fadm denominator to be the expected heterozygosity in the admixed population (2 · (1 - AF_EXP_ADMIXED_^2^)), as defined in a recent benchmark of adaptive admixture inference^34^, instead of the one applied to study selection during the Mesolithic-Neolithic transition^13^. We used the definition from the former study since the latter one defined the denominator as (2 · (1-AF_EXP_ADMIXED_)^2^ - (1-AF_EXP_ADMIXED_))^2^, which leads to misleading Fadm estimates tending to infinite when AF_EXP_ADMIXED_ tends to 0.5. Given that Davy et. al.^13^ references Cuadros-Espinoza et. al.^34^ as the source work for their Fadm definitions, we considered that the formulas they provide may be erroneous, perhaps due to misspelling, and thus we used the original ones from Cuadros-Espinoza et. al.

After the Fadm calculations, calculate_Fadm_stats.R runs fitdist from the fitdistrplus package with distr=“gamma” and method=“mle” on the Fadm calculations of the null SNP subset, resulting in a fit gamma distribution reflecting the null Fadm distribution. Then, it runs pgamma (native R package) on the Fadm calculated for all SNPs using the shape and rate parameters of the fit null gamma distribution, generating raw Fadm p-values. After the calculate_Fadm_stats.R run, IGfun.run_Fadm also calculates the bonferroni-corrected p values with statsmodels.stats.multitest.multipletests. We could calculate the Fadm and associated p-values for 4,495,609 (Germany), 4,495,621 (Ireland) and 4,495,585 (Portugal) SNPs, which are a subset of those where PCAdapt was run as we only kept sites i) with MAF>0.05 in both EF / HG populations, and ii) biallelic when considering variants imputed in the EF, HG and admixed individuals. Out of these, we defined as significant sites those passing the bonferroni correction with a p<0.05 threshold. The code to run the Fadm scan and define significant positions is in IGfun.run_SNP_selection_analyses. Note that we further coupled this analysis to a Local Ancestry Deviation (LAD) scan, explained below, to find the top Fadm hits.

#### Integrated Decay in Ancestry Tract (iDAT) scans

To specifically find signs of adaptive admixture favouring EF or HG alleles in admixed populations we used the haplotype-based iDAT (integrated Decay in Ancestry Tracts) method^135^. For each admixed population (present-day Ireland, Germany, Portugal, referred to as nIre, nGer, and nPor, respectively, in the **Main text**) we calculated unstandardized iDAT scores for each of the biallelic sites used for the Fadm scans (described above). This score reflects whether the EF Local Ancestry Tracts (EF LATs, i.e. chromosomal regions in admixed individuals with a predicted EF origin, as inferred by RFMix2) overlapping a given site are longer in average than the corresponding HG LATs. To enable LAT inference we first integrated the SNPs of EF / HG / admixed individuals into a single .vcf using bcftools merge (env:eager_env), and used them to infer the estimated number of generations between the initial admixture event and the (later) sampled individuals of each admixed population, which was required for RFMix2. For this we ran DATES as in ^12^, with default parameters (binsize = 0.001, jackknife = YES, lovalfit = 0.45, minparentcount = 10, maxdis = 1.0, qbin = 10, runfit = YES, afffit = YES), on the integrated SNPs and their genetic position mapped from the GLIMPSE ‘chr <chrom>.b37.gmap.gz’ files (defined above).

A potential issue with this approach may be that, given the different chronologies of the individuals within each admixed population, a single ‘average’ number of generations per population may not be appropriate. To address this we inferred the ‘chronological dispersion’ within each population by calculating the chronological difference between the oldest individual and the average population chronology, and converted this to a number of generations assuming 29 years / generation. In Portugal, we found that this chronological dispersion was of 27.45 generations, while the number of generations inferred by DATES was of 69.85. In Ireland, the chronological dispersion was 13.70 generations, while DATES inferred 54.14 generations. In Germany, the chronological dispersion was 3.99 generations, while DATES inferred 23.48 generations. Given that the dispersion was always significantly smaller than the inferred number of generations since initial admixture, we considered that our approach of using DATES’ ‘average’ number of generations was appropriate. Additionally, this analysis revealed a number of generations since initial admixture (23.48 - 69.85) comparable to the one for which iDAT was proposed (20 generations^135^), validating our usage of this method on our ancient populations.

Next, to obtain EF / HG LAT estimates we ran RFMix2 (command ‘rfmix (env:eager_env)) on the integrated SNPs of each chromosome using --em-iterations=2, the genetic map file ‘chr <chrom>.b37.gmap.gz’, and passing the estimated DATES generations through --generations. This worked well for autosomes, but not for chromosome X, as RFMix2 cannot interpret haploid GT calls for male chromosome X SNPs. To address this we ran this tool on a chromosome X multi-sample vcf, generated with various bcftools commands (env:eager_env) as implemented in IGfun.get_chromX_rfmix_inputs, where the samples’ columns include the GT of either i) females, or ii) pairs of males where the GT has the ‘<gt male 1>|<gt male 2>’ format. Most importantly, this generated, for each chromosome, i) the LAT assignment by the Viterbi algorithm (file ‘results.msp.tsv’), and ii) the RFMix2 log file with the estimated LAT classification balanced accuracy on simulated data generated by the program (file ‘log_file.txt’).

To validate the RFMix2 results we performed various analyses. First, to ensure correct global ancestry estimates per individual we compared the ones inferred by qpAdm (see ‘F4 statistics broad ancestral profile modeling of publicly available and newly reported datasets’) with those derived from the LAT inference as included in ‘results.msp.tsv’. Both estimates were highly correlated (**Extended Data Fig. 10a**), supporting our LAT inference. Second, to assess the LAT classification performance we checked the inferred balanced accuracy as reported in ‘log_file.txt’, which was >0.8 in all chromosomes and populations (**Extended Data Fig. 10b**), suggesting acceptable performance. Third, to validate the chromosome X male pairing strategy we checked if the RFMix2-inferred global EF ancestry for chromosome X was close to the average one across chromosomes. All individuals except three had a chromosome X ancestry within 2 standard deviations (SD) of the per-chromosome distribution, which was comparable to an equivalent analysis on autosomes 1 and 22 (two and three individuals outside the 2 SD range, respectively), supporting our male pairing strategy to enable the run of RFMix2 chromosome X.

After RFMix2 runs, we calculated the unstandardized iDAT scores using a custom python-based pipeline available at IGfun.calculate_iDAT_one_chrom, following the original definition^135^. For this, we first parsed the Viterbi results from ‘results.msp.tsv’ and used them to infer LATs. Then, for each SNP position (*p*) and ancestral EF / HG population (*pop*) we kept the corresponding LATs overlapping *p* (**Fig. 2k**) and we calculated, for all SNP positions *k* within 30Mb of *p*, the Decay in LAT (DAT) as:

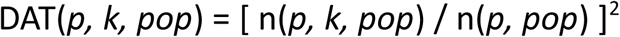

Where n(*p, k, pop*) reflects the number of chromosomes that have the same LAT of a given *pop* in both *k* and *p*, while n(*p, pop*) is the total number of LATs of a given *pop* at *p*. Note that we set 30Mb as the distance threshold for computational efficiency, and following the observation that 90% of all LATs in all populations spanned <15 Mb. Next, we calculated iDAT(*p, pop*) as the area under the curve where the x-axis is the distance between *k* and *p*, and the y-axis is DAT(*p, k, pop*), using from scipy.integrate.trapz (**Fig. 2k**). For performance reasons, we only calculated the AUC for distances where DAT(*p, k, pop*) was at least 0.25. Finally, we calculated the unstandardized iDAT for each position *p* as:

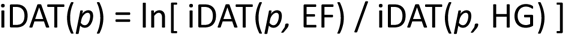

All the calculations referred to in the paragraphs above, regarding LAT inference and unstandardized iDAT calculations, were performed as implemented in IGfun.run_iDAT. To validate them, we next assessed whether we should filter out positions where either of the ancestral populations have an insufficient number of LATs (n(*p, pop*)). For this, for each admixed population (present-day Ireland, Germany, Portugal) and ancestral population *pop* (HG, EF) we randomly subsampled 100,000 *p* sites with at least 10 LATs (n(*p, pop*) == 10), which we considered to have high-confidence iDAT(*p, pop*) measurements. Then, to assess the accuracy of iDAT(*p, pop*) inference for positions with lower n(*p, pop*), we calculated iDAT(*p, pop*)_n_ values for the same *p* sites, which are equivalent to iDAT(*p, pop*) but when downsampling the number of individuals per admixed population so that we have n(*p, pop*) = 1 (iDAT(*p, pop*)_1_), 3 (iDAT(*p, pop*)_3_), 5 (iDAT(*p, pop*)_5_) or 10 (iDAT(*p, pop*)_5_). To choose an adequate threshold for the minimum n(*p, pop*) for reliable calculations, we visualized the correlation between the real, high-confidence iDAT(*p, pop*) and the downsampled iDAT(*p, pop*)_n_ values. Also, given that too strict thresholds may significantly reduce percent of genome where iDAT(*p, pop*) can be calculated for both HG and EF, we also considered the percentage of sites covered when using different thresholds (1, 3, 5, 10). Our analysis of these two variables (**Extended Data Fig. 11**) suggests that 5 is an adequate threshold to have reliable iDAT(*p, pop*) estimates while considering a significant part of the genome (30% in Portugal, 85% in Germany, 98% in Ireland), so we used it. All these calculations were implemented through IGfun.get_iDAT_tract_downsampling_simulations.

We highlight two limitations of the n(*p, pop*)>=5 threshold. First, due to limited sample size for the neolithic Portuguese population we had to filter out a large part of the genome, leading to only 30% of positions having n(*p, pop*)>=5 for both HG and EF. This means that the iDAT-based selection inference on this population may miss some true signals, and it may partially explain the low overlap between iDAT scans in different populations (see **Main text**, **Fig. 3**). Second, given the higher proportion of EF ancestry (**Fig. 1g**), we expect to miss positions with an extremely high EF LAD (i.e. where most LATs are EF as explained in the next section) when n(*p,* HG) < 5, which may unfortunately include sites where selection has favoured EF alleles. This is why we visualized the iDAT scores together with the EF LAD values (**Fig. 3**), and it may explain the low overlap between iDAT and Fadm scans. Also, this means that we may have higher power to detect HG-favouring selection with the iDAT scan. Despite these challenges, we still used the n(*p, pop*)>=5 threshold because it was essential to get robust estimates.

After these validations, we calculated the final standardized iDAT(*p*) as the z-score of all unstandardized iDAT(*p*) for all sites with n(*p, pop*) >= 5 for both HG and EF, and with MAF_EF_ > 0.05 and MAF_HG_ > 0.05. Specifically, we calculated this z-score with the numpy.mean and numpy.std methods as:

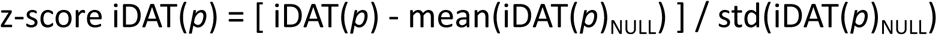

Where iDAT(*p*)_NULL_ is an array with the iDAT(*p*) of the ‘null’ sites used for Fadm p-value calculation from file ‘vars_for_Fadm_null.tab’, defined above. We could calculate the z-score iDAT(*p*) values for 3,794,425 (Germany), 4,405,969 (Ireland) and 1,307,174 (Portugal) sites, which are a subset of those where Fadm was run due to the n(*p, pop*) threshold. Out of these, we defined as significant sites those with a z-score iDAT(*p*) >= 2 (two standard deviations away from the genomic average) for selection of EF alleles, and z-score iDAT(*p*) <= -2 for selection of HG alleles. Also, to focus on independent positive selection events we identified clusters of at least 3 significant sites with the same selection direction (selection of HG or EF alleles) within 100kb of each other, and we kept one “leading” site per cluster, with highest absolute z-score iDAT(*p*) value. Then, we kept the top 10 “leading sites”, with highest absolute z-score iDAT(*p*) values, 30 in total across all three populations, as the iDAT top hits mentioned through this work. We used the 2 standard deviations as a threshold for significance as the original iDAT paper^135^ used it to find outlier sites under adaptive admixture. The code to run the iDAT scans and define significant positions is in IGfun.run_SNP_selection_analyses, and the one for defining the top hits in IGfun.get_unique_SNPselection_hits.

#### Local Ancestry Deviation (LAD) scans and definition of Fadm top hits

To infer the directionality of selection on significant sites of the Fadm scan, and to improve its robustness^13,34^, we calculated the EF Local Ancestry Deviation (EF LAD) for all sites where Fadm was calculated (described above). For this, we used the RFMix2 LAT inference from file ‘results.msp.tsv’, described above, to calculate the fraction of EF LATs at each site *p,* called fLAT(EF, *p)*, in each admixed population (Germany, Ireland, Portugal). Then, we calculated the EF LAD as a z-score (using numpy.mean and numpy.std methods) following this definition:

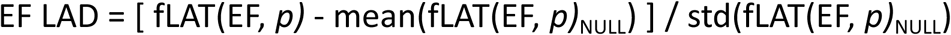

Where LAT(EF, *p)*_NULL_ is an array with the fLAT(EF, *p)* of the ‘null’ sites used for Fadm p-value calculation from file ‘vars_for_Fadm_null.tab’, defined above.

We used these EF LAD measurements to define the final Fadm top hits and infer their selection direction. Specifically, to focus on independent positive selection events we identified clusters of at least 3 significant sites with the same selection direction (EF LAD>0 or EF LAD<0) within 100kb of each other, and we kept one “leading” site per cluster, with highest Fadm value. These “leading sites”, 27 in total, are the Fadm top hits mentioned through this work. Also, we used the EF LAD to infer whether selection favoured EF alleles (EF LAD>=2), HG alleles (EF LAD<=-2), or population-independent ones (EF LAD between -2 and 2), following the approach used in previous work^13^. The code to run the Fadm-LAD scan and define significant positions is in IGfun.run_SNP_selection_analyses, and the one for defining the top hits in IGfun.get_unique_SNPselection_hits.

#### Comparison to previous studies

To enable comparisons of our top hits to previous studies of selection during the Mesolithic-Neolithic transition^6,12–14^ we integrated the regions under selection identified in such studies, as implemented in fun.get_df_regions_SNPselection_other_studies and others. For instance, we got 10 SNPs under adaptive admixture from Le et. al.^6^ from the ‘Neolithic’ hits in their Table 1, which were found through a method equivalent to our Fadm scan but treating all late Neolithic individuals as a single admixed population. These SNPs overlap the genes BDNF, PTPRV, ENSA, MAF, FTO/IRX3, FUT2, IL-1R, FAM49B, CACNB1 and PPIL2. To infer selection direction for each SNP we i) got its AF_HG_, AF_EF_, AF_ADMIXED_ (defined above) from Supplementary Data 1, ii) calculated the AF_EXP_ADMIXED_ as AF_EF_ · 0.84 + AF_HG_ · 0.16 as done in the paper, and iii) called ‘selection of EF alleles’ if the AF_EF_ - AF_HG_ of the allele increasing AF in the Neolithic admixed was >= 0.25, or ‘selection of HG alleles’ if this difference was <=-0.25. Specifically, we could find only one SNP, close to the MAF gene, with such selection of HG alleles, while the others had a small AF difference (<0.25) between the ancestral populations.

In addition, we obtained 19 genomic regions with PCAdapt hits from Irving-Pease et. al.^14^ from their Supplementary Table S2.4.3. These include the genes SLC24A5, SLC2A12, FTH1, SLC38A10, CD6, PPIAL4A, ADH1B, NYAP2, DPF3, BDNF, KCNC4, CLDND1, PIK3R5, LRRC6, SENP7, SMAD7, PLA2G3, NFKBIZ and PPFIBP1. Furthermore, we got the regions including the three genes with strong signs of adaptive admixture by the Fadm-LAD scan from Davy et. al.^13^ (HLA-E, HLA-DQB1 and SLC24A5), using their coordinates from Ensembl GRCh37.p13 matching our hg37 reference. Also, following the results of this study, we assigned selection of HG alleles for HLA-E and HLA-DQB1 regions, and selection of EF alleles for the SLC24A5 region. Note that this Fadm-LAD scan is conceptually similar to ours, but treating all late Neolithic individuals as a single admixed population.

Finally, we considered 22 SNPs that were highlighted as relevant among the results of various selection scans performed by Childebayeva et. al.^12^. For instance, this study performed three locus-specific branch length (LSBL) scans, yielding SNPs with high Fst between a target population and two background ones, conceptually similar to our PCAdapt scan. These included comparisons between i) *Linearbandkeramik* admixed German individuals (LBK) vs Anatolian Neolithic (AN) and autochthonous German Hunter Gatherers (LBK LSBL scan), ii) merged AN + LBK vs Western HG (WHG) and modern Esan individuals (EsanOut LSBL scan), and iii) merged AN + LBK vs WHG and African HG (AfricaOut LSBL scan). Also, they performed an adaptive admixture scan with the AIMLESS method, conceptually similar to our Fadm scan, yielding SNPs with an unexpected AF in LBK as compared to the expectation from the mixture between AN and WHG (AIMLESS scan). Furthermore, they performed an XP-EHH scan comparing LBK to WHG, which is similar to our XP-EHH scan but may also reflect adaptive admixture signs given that the comparison includes LBK (admixed population) instead of EF.

Following the rationale of this study^12^, we considered as highlighted SNPs those that matched one of six different criteria. First, we considered 7 SNPs significant in all three LSBL scans, including those overlapping APOM, BDNF, IGSF21, ILDR2, LMF1, MHC and PDE2A; as provided in their Supplementary Table 11. Second, we included 3 SNPs found in the LSBL AfricaOut scan, overlapping CD82, MTHFR and NBPF3; as provided in their Supplementary Table 10. Third, we included 1 SNP found in the LSBL EsanOut scan, overlapping SLC24A5; as provided in their Supplementary Table 9. This SNP also had an XP-EHH>0 in their XP-EHH scan, so we assigned ‘selection of EF alleles’ to it given the predominant EF ancestry of LBK individuals^12^. Fourth, we considered 1 SNP found significant by both LBK LSBL and XP-EHH scans, overlapping FADS1; as provided in their Table 1. This SNP had an XP-EHH<0 in their XP-EHH scan, so we assigned ‘selection of HG alleles’. Fifth, we considered 1 SNP found by one of the LSBL scans (unclear from the text) and the XP-EHH scan, overlapping JAK1; with coordinates provided the legend of their Figure 4. Sixth, we included 9 SNPs found by all their scans (LSBL, AIMLESS and XP-EHH ones), overlapping LEPR, IL13, HLA-A, HLA-B, HLA-DQB2, CHRAC1, BMP1, SORBS1 and ELMO1; as provided in their Supplementary Table 12. For these, we assigned ‘selection of EF alleles’ if XP-EHH>0 and ‘selection of HG alleles’ if XP-EHH<0. All these regions under selection in previous studies can be found in **Supplementary Table 3**.

To define regions of interest for validating our selection scans we identified windows of the genome including at least two of these regions under selection (i.e. identified by either different studies or by different scans of the same study) within 1 Mb of each other. We found six of them, available at **Extended Data Fig. 14,15**. Also, we added the coordinates of these six windows to our main Manhattan plots (**Fig. 3**), highlighting one specific gene per window (SLC24A5, NFKBIZ, JAK1, FADS1, BDNF and HLA-B, respectively).

### Copy Number Variant analysis

#### Per sample CNV calling

To perform CNV analyses we first inferred CNVs per sample with a custom, python-based pipeline, run on the 10, 20, 50 and 100kb per-window coverage calculations for chromosomes 1-22 and X, generated as described in “Coverage calculations” above. For each window size and analyzed sample (i.e. belonging to the EF/HG ancestral or one of the admixed populations), we calculated the per-window mean coverage normalized by the mean autosomal coverage, hereafter referred to as “rel_cov”. Then, we assigned to each sample, in each window, a copy number (CN) of 0,1,2,3,4 depending on its rel_cov, individual’s sex and chromosome. For chromosomes 1-22 and X chromosome in females, CN=0 was rel_cov in [0, 0.1), CN=1 was rel_cov in [0.1, 0.7), CN=2 was rel_cov in [0.7, 1.3], CN=3 was rel_cov in (1.3, 1.8] and CN=4 was rel_cov in (1.8, inf]. Thus, DEL CNVs would be windows with CN=0,1, and DUP CNVs would be windows with CN=3,4. Conversely, for X chromosome in males CN=0 was rel_cov in [0, 0.1), CN=1 was rel_cov in [0.1, 0.8], and CN=2 was rel_cov in (0.8, inf]. Accordingly, DEL CNVs would be windows with CN=0, and DUP CNVs would be windows with CN=2. Next, we filtered out “non-biallelic” windows, with more than 1 type of CNV (DEL or DUP). Furthermore, we grouped adjacent windows with the exact same CN patterns across all samples. Each of these grouped windows with equal patterns was defined as a CNV in a given sample.

To choose an adequate window size for subsequent analyses we ran the same CNV calling pipeline on each set of sample-tailored simulated datasets (based on the variants of “VLASA7”, “KK1” and “LEPE48”, described above), where no CNVs are expected. Then, to choose a window size with a reduced false positive rate we visualized the distribution of the number of samples with a given CNV across different simulations and real samples (**Extended Data Fig. 16**). This analysis suggested that 20kb is the lowest window size with an acceptable performance, for which there are very few CNVs called in simulated datasets, with almost all found in a single simulated sample. Thus, we used CNV calls on 20kb windows for downstream analyses. All these calculations were performed as implemented in IGfun.get_called_CNVs_all_samples.

#### Comparison of CNVs in ancestral EF and HG populations

To identify CNVs with the highest differentiation between the ancestral populations (EF vs HG) we calculated Wright’s F_ST_ (fixation index) between HG and EF for each CNV as specified in https://reich.hms.harvard.edu/sites/reich.hms.harvard.edu/files/inline-files/fstnote_1.pdf. This is a value between 0 and 1 that indicates whether the AF of the CNV is highly differentiated between the two populations. For each CNV, if *x* is the AF of the variant in the HG, and *y* in the AF in EF, we calculated *F_ST_ = (x - y)*^2^ */ (x + y - 2xy)*. The distribution of F_ST_ values suggested various interesting CNVs with high F_ST_ values (e.g. F_ST_ > 0.2, **Fig. 4b**), but this might be due the ‘multiple testing’ bias of comparing thousands of CNVs. For instance, by chance we may find such high F_ST_ values for a few windows when comparing individuals from the same population. To address this we calculated an empirical p value based on resampling our data, reflecting the probability to find an F_ST_ >= observed F_ST_ , when randomly comparing samples from the same population. We generated 100 sets of equivalent A/B populations, where both A and B have a mix of randomly picked HG (8 of a total of 16) and EF (25 or 26 of a total of 51). Then, we calculated the following p value for each real CNV:

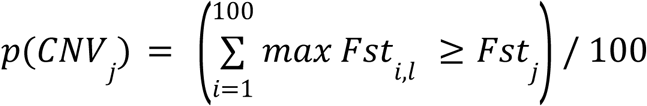

Where *j* represents the CNV, *l* is the length of the CNV*_j_*, *max Fst_i,l_* is the maximum F_ST_ observed for a CNV with length >= *l* in random set *i* and *Fst_j_* is the observed value for *CNV_j_*. This calculation is inspired by the maxT GWAS method^167,168^, and it yields a p value for each CNV corrected for multiple testing biases, as it relies on maximum intra-population F_ST_ measurements.

To identify the final set of significantly differentiated CNVs we kept those that had an AF across all EF and HG samples >= 0.05, a maxT p < 0.01, and an F_ST_ >= 0.2 (**Fig. 4c**). This yielded thirteen 20kb-windows with differential deletions, and three 20kb-windows with differential duplications. Since some of these regions may reflect the same CNV event, we clustered those that were within 10 windows (for which coverage was calculated) of each other, had the same type of CNV and direction (i.e. AF higher in EF or *vice versa*). This resulted in eight clusters for deletions, and three for duplications, 11 clusters. To get the final set of CNVs we defined one CNV region for each of these 11 clusters, where we had consecutive windows with the same CNV type and direction, and an F_ST_ >= 0.05. Also, after manual inspection of their coverage profiles (**Extended Data Fig. 17**), we decided to discard a 820kb duplication on chromosome 16, since it was driven by only a few HG individuals with noisy coverage. This resulted in a final set of 10 highly differentiated CNVs. All these calculations were performed as implemented in IGfun.generate_CNV_comparisons_ancestral_pops and IGfun.get_df_CNVs_ancestralPops_filt.

#### CNV analysis in admixed populations

To detect CNVs under adaptive admixture in the late Neolithic admixed populations (from present-day Germany, Ireland, Portugal, referred to as nGer, nIre and nPor, respectively, in the **Main text**) we performed an Fadm scan for the CNVs of each of them, equivalent to the one performed for SNPs (see above). For this, we included the 20 kb CNVs with a MAF > 0.05 in both ancestral HG/EF populations into a ‘vars_for_Fadm.tab’ file, similar to the one generated for SNPs. Also, we gathered the ‘vars_for_Fadm_null.tab’ null file from the SNP Fadm scans, defined above, to enable p-value calculations. Note that we tried to generate a null set of CNVs, instead of SNPs, but it was deemed unfeasible due to a low number of CNVs. For instance, taking CNVs 5Mb away from each other would total a maximum of 452 CNVs across populations, covering 15.67% of the genome. We deemed this problematic due to both i) the low number of CNVs considered, which may be insufficient for the gamma distribution fit, and ii) the large fraction of the genome covered, perhaps leading to a biased null distribution. Thus, we used the null SNP distribution for p-value calculations. Finally, we ran calculate_Fadm_stats.R, defined above, to get Fadm and its related p-values for all CNVs in each admixed population. Also, we used statsmodels.stats.multitest.multipletests to calculate bonferroni-corrected p values, and we kept as significant p-values those with bonferroni-corrected p<0.05. This yielded one duplication with unexpectedly low AF in the German population. All of these calculations were performed as implemented in IGfun.generate_CNV_comparisons_admixed_pops.

### Functional variant annotations

For functional variant annotations we i) downloaded the relevant annotation databases, ii) identified the affected genes and phenotypes with IGfun.get_df_sig_vars_annotated, iii) annotated the genes with IGfun.get_df_merged_vars_annotations_with_Gprofiler_info, and iv) integrated all the relevant information into **Supplementary Table 2** with IGfun.get_df_top_hits_annotated_with_popGen_stats. In the following sections we describe in detail how we performed all these processes.

#### Preparation of annotation databases

To enable functional annotations of variants we downloaded various databases and files. To get variant-phenotype mappings from the GWAS catalogue (v1.0.2) we downloaded all associations from https://www.ebi.ac.uk/gwas/api/search/downloads/alternative at 11-March-2024. Also, to map variants to transcriptomic effects we used the *cis* expression Quantitative Trait Locus (eQTL) annotations per tissue from the GTEx database (v8). We obtained all cis-eQTL annotations, which are SNP-gene pairs in specific tissues, from https://storage.googleapis.com/adult-gtex/bulk-qtl/v8/single-tissue-cis-qtl/GTEx_Analysis_v8_eQTL.tar. In addition, we obtained fine-mapped eQTL annotations, which include less redundancy from Linkage Disequilbrium (LD) between SNPs, from https://storage.googleapis.com/adult-gtex/bulk-qtl/v8/single-tissue-cis-qtl/GTEx_Analysis_v8_eQTL_independent.tar.

Given that the most available functional annotations (including those from GWAS catalogue and GTEx) use GRCh38-based coordinates, and not GRCh37 as we did, we downloaded various files for this assembly (version GRCh38.p14) from the Ensembl Build 111^169^, available at https://ftp.ensembl.org/pub/release-111. Specifically, we got the reference sequence fasta (“fasta/homo_sapiens/dna/Homo_sapiens.GRCh38.dna.primary_assembly.fa.gz”) and gene annotation files (“gff3/homo_sapiens/Homo_sapiens.GRCh38.111.gff3.gz”, “gtf/homo_sapiens/Homo_sapiens.GRCh38.111.gtf.gz”). Furthermore, to lift the variants between the GRCh37 and GRCh38 builds and vice-versa we obtained the corresponding chain files from https://ftp.ensembl.org/pub/release-111/assembly_chain/homo_sapiens/ (“GRCh37_to_GRCh38.chain.gz” and “GRCh38_to_GRCh37.chain.gz”). Finally, to be able to re-map gene identifiers to the Ensembl canonical set we downloaded https://ftp.ensembl.org/pub/release-111/tsv/homo_sapiens/Homo_sapiens.GRCh38.111.canonical.tsv. Note that the GRCh38.p14 was also used by the GWAS catalogue, which ensured proper mappings.

To get predictions of variant impacts on nearby genes and phenotypes (e.g. from COSMIC or ClinVar) we used the Ensembl Variant Effect Predictor (VEP), which relies on the index caches with the Ensembl feature annotations, downloaded from https://ftp.ensembl.org/pub/release-111/variation/indexed_vep_cache/homo_sapiens_vep_111_GRCh38.tar.gz. Also, to enable the usage of various VEP plugins (AlphaMissense, GO, Phenotypes, Enformer, StructuralVariantOverlap) we downloaded various files as specified in http://www.ensembl.org/info/docs/tools/vep/script/vep_plugins.html, clarified next.

First, for the plugin ‘AlphaMissense’ we i) downloaded pre-computed annotations from https://storage.googleapis.com/dm_alphamissense/AlphaMissense_hg38.tsv.gz on 4-April-2024 and ii) indexed the file with ‘tabix -s 1 -b 2 -e 2 -f -S 1 AlphaMissense_hg38.tsv.gz’ (env:perSVade_env). Second, for the plugin ‘GO’ we downloaded the related database by running VEP with arguments ‘--plugin GO,file=./go_terms_GRCh38.gff.gz --assembly GRCh38’ on 4-April-2024. Third, for the plugin ‘Phenotypes’, we downloaded the related database by running VEP with arguments ‘--plugin Phenotypes,file=./phenotypes_vep_GRCh38.gff.gz’ on 4-April-2024. Fourth, for the plugin ‘Enformer’ we downloaded annotations and related .tbi file from https://ftp.ensembl.org/pub/current_variation/Enformer/enformer_grch38.vcf.gz on 3-October-2024. Fifth, for the plugin ‘StructuralVariantOverlap’ we obtained the variants and related .tbi file from https://ftp.1000genomes.ebi.ac.uk/vol1/ftp/phase3/integrated_sv_map/ALL.wgs.mergedSV.v8.20130502.svs.genotypes.vcf.gz.

On another note, to map GRCh38-lifted SNPs to previously-defined variant IDs we downloaded the dbSNP vcf (build 156)^170^ at https://ftp.ncbi.nih.gov/snp/archive/b156/VCF/GCF_000001405.40.gz. Also, to enable fast access we downloaded its matching index file from https://ftp.ncbi.nih.gov/snp/archive/b156/VCF/GCF_000001405.40.gz.tbi. All of these databases were used for the functional annotations mentioned below.

#### Functional annotation of CNVs

To infer the effect of 11 CNVs with differential AF between populations, described above, on nearby genes and their previously-linked phenotypes, we used IGfun.get_df_sig_vars_annotated. This function first uses custom python code and GATK LiftoverVcf (env:perSVade_env) to lift the CNV GRCh37 coordinates to GRCh38 ones (see IGfun.get_df_vars_with_GRCh38_coordinates), as this assembly build had more available functional annotations. For this, we used the corresponding GRCh37-to-GRCh38 chain and reference genome builds described above. Next, it validates that none of the 11 CNVs changed its length upon lifting by >10kb. Furthermore, it runs IGfun.run_vep_df_vars on the lifted CNVs to i) check their overlap with previously defined CNV IDs, ii) annotate their effect on nearby genomic features and iii) annotate the phenotypes or functions related to such features.

More specifically, this function first runs VEP (singularity image ensembl-vep:release_111.0) on the GRCh38-coordinated CNVs with various custom arguments. First, to be able to run offline we provide the Ensemble annotations with ‘--offline --cache --dir_cache <annotations path>/homo_sapiens_vep_111_GRCh38_indexed_cache’, generated as described above. Second, to limit the analysis to transcripts belonging to the GENCODE basic set, without fragmented or problematic transcripts, we set ‘--gencode_basic’. Third, to retrieve information about the closest genes we used ‘--nearest gene’. Fourth, to output only one, most severe consequence per gene we set ‘--per_gene’. Fifth, to map the affected genes and variants to further functional annotations we ran various VEP plugins with ‘--plugin <plugin name>,<plugin parameters>’, explained next.

For these CNV annotations, we ran IGfun.run_vep_df_vars with plugins=[“GO”, “StructuralVariantOverlap”, “Phenotypes”], types_phenotypes_list=[“Gene”, “RegulatoryFeature”] and sources_phenotypes_list=[“G2P”, “MIM_morbid”, “Orphanet”, “Cancer_Gene_Census”], which parametrize the used plugins. This results in the setting of various arguments in VEP to retrieve i) Gene Ontology annotations for affected genes (--plugin GO,file=go_terms_GRCh38.gff.gz), ii) information about overlapping previously-defined structural variants (--plugin StructuralVariantOverlap,file=ALL.wgs.mergedSV.v8.20130502.svs.genotypes.vcf.gz,percentage=80, reciprocal=1,cols=’SVTYPE;MSTART;MLEN;MEND;MEINFO;SVLEN;END;CS;MC;AF;NS;EAS_AF;EUR_A F;AFR_AF;AMR_AF;SAS_AF;SITEPOST’) and iii) phenotypic information about the affected genes from Cancer Gene Census, G2P, MIM morbid and Orphanet (--plugin Phenotypes,file=phenotypes_vep_GRCh38.gff.gz,phenotype_feature=1,include_types=’Gene&RegulatoryFeature’,include_sources=’G2P&MIM_morbid&Orphanet&Cancer_Gene_Census’).

After the VEP run, IGfun.run_vep_df_vars runs various custom python functions to add further information and ensure the consistency in the obtained annotations. For instance, it creates a column reflecting the gene to which a variant is mapped (column ‘mapped_Gene’), which may be one set by VEP (column ‘Gene’) or the nearest one (column ‘NEAREST’) for variants affecting intergenic regions. In addition, it runs IGfun.get_df_vep_with_checked_Phenotypes to format and debug the annotations provided by the ‘Phenotypes’ plugin. For instance, this function verifies that all annotated genes correspond to the Ensembl canonical ones (defined above). Also, it uses IGfun.get_phenotypes_one_source_one_r to add, for each source database (e.g. “G2P” or “MIM_morbid”), one column to the VEP output with all the information from phenotypes_vep_GRCh38.gff.gz about that source.

Note that the VEP StructuralVariantOverlap plugin yielded no overlaps, which may be due to i) the fact that these are truly novel CNVs or ii) inaccurate mapping of structural variants between databases due to uncertain definition of their boundaries.

#### Functional annotation of SNPs

After the CNV annotation, the function IGfun.get_df_sig_vars_annotated maps each of the 85 top hit SNPs to nearby genes, previously-linked phenotypes and transcriptional effects. For this, the function firstly uses IGfun.get_df_vars_with_GRCh38_coordinates, described above, to lift the SNP GRCh37 coordinates to GRCh38 ones. Then, it uses IGfun.get_df_SNPs_with_functional_annotations on these GRCh38-based variants to obtain annotations from VEP (and related databases), dbSNP, GWAS catalogue and GTEx, described next.

Specifically, this function first runs our custom VEP pipeline IGfun.run_vep_df_vars as described above for CNVs, but with different arguments: plugins=[“GO”, “AlphaMissense”, “Enformer”, “Phenotypes”], types_phenotypes_list=[“Gene”, “Variation”, “RegulatoryFeature”] and sources_phenotypes_list=[“G2P”, “MIM_morbid”, “Orphanet”, “Cancer_Gene_Census”, “ClinVar”, “GEFOS”, “GIANT”, “MAGIC”, “COSMIC”, “dbGaP”]. This results in the setting of various arguments in VEP to retrieve i) Gene Ontology annotations for affected genes (--plugin GO,file=go_terms_GRCh38.gff.gz), ii) predictions of pathogenicity in missense SNPs from AlphaMissense (--plugin AlphaMissense,file=AlphaMissense_hg38.tsv.gz), iii) predictions of variant impacts on chromatin and gene expression from Enformer (--plugin Enformer,file=enformer_grch38.vcf.gz) and iv) phenotypic information about the affected genes from Cancer Gene Census, G2P, MIM morbid and Orphanet, as well as SNP-phenotype associations from ClinVar, GEFOS, GIANT, MAGIC, COSMIC and dbGaP (--plugin Phenotypes,file=phenotypes_vep_GRCh38.gff.gz,phenotype_feature=1,include_types=’Gene&Vari ation&RegulatoryFeature’,include_sources=’ClinVar&dbGaP&G2P&GEFOS&GIANT&MAGIC&MIM_ morbid&Orphanet&Cancer_Gene_Census&COSMIC’).

Next, to obtain dbSNP identifiers and AF in modern human populations for each of the SNPs, IGfun.get_df_SNPs_with_functional_annotations runs ‘bcftools view’ on the dbSNP vcf (defined above) (env:perSVade_env) and custom python code, through IGfun.get_df_vars_with_dbSNP_ids. We could find dbSNP identifiers for 85/85 SNPs, which validates our approach of further using these identifiers to link SNPs to GWAS catalogue associations, as described below.

To infer whether some of the SNPs affect gene expression, IGfun.get_df_SNPs_with_functional_annotations annotates if they have been described as cis-eQTLs in the GTEx database (described above). Such eQTLs are SNP-to-gene mappings for various human tissues, where it has been described that the SNP alternative allele is associated with a higher or lower expression of the mapped gene in that tissue. We considered two types of eQTLs, depending on the level of confidence of the association. On the one hand, we considered associations from “GTEx_Analysis_v8_eQTL/<tissue name>.v8.signif_variant_gene_pairs.txt.gz” (described above), which include all eQTLs with no fine-mapping to remove the redundancy between linked SNPs, corresponding to the column ‘eQTL_all’ in **Supplementary Table 2**. On the other hand, as mappings of higher confidence we considered those in “GTEx_Analysis_v8_eQTL_independent/<tissue name>.v8.independent_eqtls.txt.gz” (described above), which are based on a stepwise regression algorithm that ensures that there are no redundant eQTLs coming from LD between close SNPs. These correspond to the column ‘eQTL_indep’ in **Supplementary Table 2**.

To enable the mapping between our SNPs and those in GTEx, IGfun.get_df_SNPs_with_functional_annotations first runs IGfun.get_df_pcadapt_pvalues_filt_eQTL_data. This is a python-based pipeline that integrates all eQTLs (“all” and “indep”) into a single table, and keeps only those that involve i) some SNPs and ii) genes that have a qval<0.05 in “GTEx_Analysis_v8_eQTL/<tissue name>.v8.egenes.txt.gz”, so they have some significant associations. Then, it uses the python-based function IGfun.get_df_pcadapt_pvalues_filt_with_added_eQTL_data to merge these eQTL annotations with the VEP output in a way that each row is a unique combination between SNP and gene, as in **Supplementary Table 2**. Specifically, this function first records, for a given SNP and gene combination, all eQTLs on different tissues, resulting in strings like ‘Testis, slope=0.80 | Thyroid, slope=-0.28’ where positive / negative slopes indicate predicted up/downregulation. Then it adds these annotations to the VEP table in two ways. If the SNP-gene involved in a given eQTL matches some VEP SNP-gene mapping, the eQTL is added to the equivalent VEP row, matching the ‘mapped_Gene’ set by VEP. Else, the eQTL annotation is added as an additional row, with a new ‘mapped_Gene’.

To gather additional information about the effects on human phenotypes IGfun.get_df_SNPs_with_functional_annotations also links the SNPs to reported SNP-phenotype associations from the GWAS catalogue (described above). First, it uses IGfun.get_df_GWAS_catalog_hits, a python-based pipeline, to process the GWAS catalogue datasets. This function first obtains the SNP-phenotype associations for the SNPs, mapped though the dbSNP identifiers. Then, it keeps only those associations with a “STRONGEST SNP-RISK ALLELE” that is either i) the reference genome one, ii) the alternative allele from the SNP or iii) uncertain “?”. Furthermore, our pipeline uses IGfun.get_df_pcadapt_pvalues_filt_with_added_GWAS_hits to format the GWAS hits in a way that each hit is related a single gene, obtained from the SNP-gene VEP mappings, resembling the eQTL formatting. Also, this function formats hits like “Neuroticism, risk_allele=REF, risk=increase (B=8.40)”, indicating the phenotype, risk allele, and whether there is an increase or decrease in risk. These hits can be found in the column “GWAScatalog_hits” in **Supplementary Table 2**.

In summary, for each SNP we obtained predictions about their impact on i) nearby genes, as provided by VEP, ii) pathogenicity according to AlphaMissense, iii) human phenotypes (from COSMIC, dbGAP, ClinVar, GEFOS, GIANT, MAGIC and GWAS catalogue) and iv) gene expression (from GTEx cis-eQTLs and Enformer). Also, for all of the impacted genes, as predicted by VEP, we obtained functional / phenotypic information from Cancer Gene Census, G2P, MIM morbid and Orphanet. However, note that, due to lack of overlaps, in **Supplementary Table 2** we do not report annotations from AlphaMissense, dbGAP, ClinVar, GEFOS, GIANT and MAGIC. Also, we omitted the annotations from Enformer due to a lower interpretability as compared to GTEx eQTL annotations.

#### Integration of SNPs and CNVs

After annotating SNPs and CNVs, the function IGfun.get_df_sig_vars_annotated integrates all annotations and adds extra relevant fields using custom python code, which are also in **Supplementary Table 2**. For instance, the field “mapped_Gene_type” indicates the source of the variant-gene mapping, which can come from either i) a predicted impact (e.g. missense, synonymous, upstream…) of the variant on the gene (mapped_Gene_type=”Gene”), ii) the nearest gene, for intergenic variants (mapped_Gene_type=”Nearest_Gene”) or iii) eQTL predictions (mapped_Gene_type=”eQTL_Gene”). Also, the “mapped_Gene_name” indicates the common name for the mapped gene, parsed from either the GRCh38 gtf annotations (described above), or the GTEx files, if mapped_Gene_type=”eQTL_Gene” and the gene was not found in such gtf. This generates a ‘merged variants table’, including most of the information found in **Supplementary Table 2**.

#### Functional annotation of genes affected by SNPs and CNVs

To obtain functional annotations for the mapped genes, we used IGfun.get_df_merged_vars_annotations_with_Gprofiler_info. This function uses the object Gprofiler from the gprofiler package, which is a python-based API to the g:Profiler database. Initially, it obtains gene names, descriptions and ID mappings for each gene with Gprofiler.convert, run with organism=“hsapiens”. Then it gets annotations with Gprofiler.profile, run with sources=[“GO:MF”, “GO:CC”, “GO:BP”, “KEGG”, “REAC”, “WP”, “TF”, “MIRNA”, “HPA”, “CORUM”, “HP”], organism=“hsapiens”. This obtains annotations from Gene Ontology (GO), KEGG, Reactome, WikiPathways (WP), TRANSFAC (TF), miRTarBase (MIRNA), Human Protein Atlas (HPA), CORUM protein complexes and Human Phenotype Ontology (HP).

Furthermore, this function adds these gene names, descriptions and functional annotations to the ‘merged variants table’ described above. To filter out excessively general annotations we kept those with a term_size_fraction<=0.25, calculated as the ratio between “term_size” (number of genes that are mapped to a given annotation) and “effective_domain_size” (total number of genes considered). Both “term_size” and “effective_domain_size” are provided by Gprofiler.profile. These can be found in the fields “description” and “gprofiler_<database name>” from **Supplementary Table 2**.

#### Generation of final functional annotation table

To write the actual **Supplementary Table 2**, including population genetic statistics and relevant functional annotations, we ran IGfun.get_df_top_hits_annotated_with_popGen_stats. This function takes the annotated variants (described above) and the selection scan results, and integrates them into a single table. Also, note that this function re-writes the GWAS risk allele for rs6476690 (see **Main text**), which was undetermined in the GWAS catalogue, but according to the related study^171^ it was the alternative allele (rs6476690-A).

### Functional enrichments

#### Preparation of eQTL database

To run functional and tissue enrichment analyses related to eQTL annotations we first gathered all of such annotations from GTEx into a single dataset. For this we ran IGfun.get_df_all_eQTLs_file, which integrates all eQTLs (“all” and “indep”, defined above) and keeps only those that involve genes that have a qval<0.05 in “GTEx_Analysis_v8_eQTL/<tissue name>.v8.egenes.txt.gz”, so they have some significant associations. This generated a table where each row is one combination of gene, SNP, type of eQTL (“all” or “indep”), tissue, and eQTL direction (upregulation or downregulation).

#### Functional enrichments

To find functions enriched in the 18 gene sets mentioned in the **Main text** (PCAdapt_overExp_<POP>, XPEHH_diffExp_<POP>, Fadm_<direction de>_<adm pop>, iDAT_diffExp_<POP>_<adm pop>, <POP>_DEL), we ran the following steps. Given that most of the eQTLs mapped to the SNPs under selection are of the “all” type (involving 173 genes, while “indep” eQTLs only involved 8 genes), we filtered the database of all eQTLs, defined above, to only include “all” eQTLs. Also, we only kept eQTLs of tissues with eQTLs mapped to our SNPs under selection. Then, for each of the 18 gene sets (defined with IGfun.get_list_type_genes_enrichments and IGfun.get_df_vars_only_certain_genes based on the table of SNP / CNV hits), we ran a functional enrichment analysis with GProfiler.profile from the gprofiler package, with the arguments organism = “hsapiens”, sources = [“GO:MF”, “GO:CC”, “GO:BP”, “KEGG”, “REAC”, “WP”, “TF”, “MIRNA”, “HPA”, “CORUM”, “HP”], significance_threshold_method = “g_SCS”, user_threshold = 0.05. Also, we provided as gene universe (background argument) either i) all genes mapped to eQTLs in our filtered database (for the gene sets PCAdapt_overExp_<POP>, XPEHH_diffExp_<POP>, Fadm_<direction DE>_<adm pop> and iDAT_diffExp_<POP>_<adm pop>), or ii) the union of all genes mapped to eQTLs in our filtered database and those affected by some CNV (see **Supplementary Table 2**) (for gene sets <pop_del>). Next, we calculated the odds-ratio of the enrichment with IGfun.get_OR_enrichments, and the ‘term size fraction’ by dividing the term_size by the effective_domain_size as provided by GProfiler.profile. All these processes were performed as implemented in IGfun.get_df_enrichments_all.

To focus on the most relevant enrichments we kept those with ‘term size fraction’ <= 0.05, OR >= 1, and discarded those related to the TF and MIRNA databases. Then, to facilitate redundancy reduction and visualization we calculated a distance metric between all pairs of enriched terms (t_1_, t_2_), calculated as 1 - G_1,2_, where G_1,2_ is the Jaccard index between the set of genes included in any of the 18 gene sets with t_1_ or t_2_ annotations. Next, to get a set of representative terms for each gene set we ran IGfun.get_df_enrichments_with_non_redundant_terms, which runs a REVIGO-like pipeline for redundancy reduction, but adapted to filtering terms from different database sources.

Specifically, this function iterates through pairs of terms (t_1_, t_2_) with G_1,2_ >= 0.1 and term description text similarity (D_1,2_) >= 0.1 (calculated with sklearn.feature_extraction.text.TfidfVectorizer and sklearn.metrics.pairwise.cosine_similarity, see IGfun.get_text_similarity), sorted hierarchically by i) descending G_1,2_, ii) descending D_1,2_, iii) ascending t_1_ ID alphabetical order, and iv) ascending t_2_ ID alphabetical order. In each iteration, it defines a term (t_1_ or t_2_) as ‘redundant’, following various criteria defined next. First, if one term is significantly more general than the other (i.e. ‘term size fraction’ difference >= 0.025), the former is defined as ‘redundant’. Second, if the terms come from different source databases, we prioritize sources following this order: [’HP’, ‘KEGG’, ‘WP’, ‘REAC’, ‘GO:BP’, ‘GO:CC’, ‘GO:MF’, ‘HPA’, ‘CORUM’, ‘TF’, ‘MIRNA’]. Third, if one term has a much lower average corrected p-value than the other across significant enrichments, i.e. p_1_< (p_2_*0.5) or *vice versa*, we keep it. Fourth, if none of the other criteria are met, we keep the term which is first in the alphabetical sorting of the t_1_ and t_2_ IDs. Using this redundancy reduction pipeline we could reduce the number of enrichments from 213 to 51.

All of these enrichments can be found in **Supplementary Table 6** and **Fig. 7d**, where the field “non_redundant” indicates terms defined as non-redundant. Also, note that we chose the thresholds on G_1,2_ (0.1), D_1,2_ (0.1) and ‘term size fraction’ difference (0.025) based on manual inspection of the redundancy-reduction process to verify biologically meaningful results. All of these calculations were performed as implemented in IGfun.plot_heatmap_enrichments.

#### Tissue enrichments

To understand which tissues were mostly affected by differential expression we evaluated whether different gene sets (PCAdapt_overExp_<POP>, XPEHH_diffExp_<POP>, Fadm_<direction DE>_<adm pop>, iDAT_diffExp_<POP>_<adm pop>, see **Main text** and **Fig. 7**) tended to involve eQTLs from specific tissues over others. For this, for each gene set we calculated the number of predicted differentially expressed (DE) genes (n_DE_) in each tissue, according to their eQTLs, and also the empirical probability of observing n_DE_ or higher by chance p(n_DE_). To calculate this probability, we first generated a filtered eQTL database including “all”-type eQTLs for tissues with eQTLs mapped to our SNPs under selection. Then, for each gene set and tissue, associated with a certain number of eQTLs (n_eQTL_), we randomly picked 2,000 times n_eQTL_ from the filtered eQTL database and calculated the number of predicted DE genes obtained for that tissue (n_DE,RANDOM_). Thus, we calculated p(n_DE_) as the fraction of these 2,000 resamples that have n_DE,RANDOM_ >= n_DE_, setting it to 0.00025 (0.5/2000) if there were none. Also, we calculated the bonferroni-corrected p(n_DE_) with statsmodels.stats.multitest.multipletests. Note that for computational efficiency we performed the p(n_DE_) calculations based on a random subset of 610,000 filtered eQTLs instead of the whole filtered database, i.e. 10,000 times the maximum number of n_eQTL_ to draw (61), including 2382-9924 eQTLs for all considered tissues.

In addition, to provide a more general view of the affected tissues we performed an equivalent analysis but grouping tissues into tissue groups of the same system, e.g. grouping all central nervous system (CNS) or circulatory system tissues. For this, we used IGfun.get_general_tissue_from_eQTL_tissue to define the mapping between each tissue with eQTLs mapped to SNPs under selection and the following general tissue groups: fibroblasts, EBV lymphocytes, circulatory, digestive, skin, reproductive_female, reproductive_male, endocrine, adipose, CNS, Breast_Mammary_Tissue, Nerve_Tibial, Kidney_Cortex, Pancreas, Liver, Muscle_Skeletal and Lung. Then, we re-calculated the n_DE_ and bonferroni-corrected p(n_DE_) but considering tissue groups instead of single tissues. These are the numbers of genes and p values mentioned in the **Main text** and **Fig. 7c**. All these processes were performed as implemented in IGfun.get_df_enrichments_tissues.

### Pathogen detection

To assess whether an agricultural lifestyle led to increased pathogen exposure, we compared the content in microbial or viral sequences of samples with unadmixed HG ancestry vs those with unadmixed EF ancestry, inferred from the unmapped reads. To minimize the effect of the pathogen survivor bias on the EF vs HG comparison, we did not analyze samples where the aDNA was obtained from petrous bones, as these are unlikely to have been colonized by ancient pathogens^7^. Furthermore, to avoid unreliable results, we filtered out one AADR EF sample where the sequencing data comes from different library preparation types (e.g. with and without UDG treatment and/or with double or single-stranded preparations) and its metadata did not allow to link the libraries to the preparation method. These filterings resulted in a set of 5 HG and 1 EF, including i) some of the newly reported EF and HG Iron Gates individuals, and ii) three individuals contained in the AADR HG (I0017.SG, SF9.SG, Syltholm1.SG) from present-day Northern-European countries (Sweden, Sweden, Denmark, respectively). Given that geographical distance may influence pathogen dispersal dynamics, we considered that the absence of EF samples geographically close to the Northern-European HG could be a biasing factor in our comparison. Thus, to ensure a geographically balanced comparison we only analyzed the Iron Gates individuals (2 HG, 1 EF) sequenced here.

To examine whether these ancient individuals presented any pathogen signal, the unmapped portion of the genomic data was taxonomically classified using the HOPS pipeline in nf-core/eager (Hübler et al., 2019; Fellows Yates et al., 2021). We only found a significant signal for the one EF individual (i701-i501), whose sequenced libraries had a substantial proportion of non-human reads with hits to the human Hepatitis B Virus (HBV) clade (see **Main text**). To validate this finding, we mapped the non-human portion of the generated genomic data to Hepatitis B virus reference (NC_003977) with nf-core/eager (v2.4.7). In short, unmapped reads were aligned with BWA (v0.7.17-r1188v.0.1.17) aln -l 16 -t 8 -n 0.01 -o 2 -q 0. Resulting bam files were deduplicated with Picard MarkDuplicates (v2.26.0) and filtered for mapped reads with a minimum mapping quality of 37. Genotyping was done with GATK UnifiedGenotyper (v.3.5-0-g36282e4) with the options EMIT_ALL_SITES. We built a consensus sequence for the HBV genome with the GenConS module of the TOPAS toolkit^172^ requiring a minimal coverage of the position for the major allele to be 3x. The newly reconstructed HBV consensus genome was then added to the multiple sequence alignment including the HBV datasets described in ^19^ with mafft^81^ (v7.305b). The resulting multiple sequence alignment was used to build a Maximum Likelihood tree with RaxML-NG^173^ (v0.9.0git) with the following parameters: --all --model GTR+G --bs-trees autoMRE; to assess the phylogenetic placement of new HBV genome.

### Transposable Element (TE) analysis

#### TE comparative analysis for ancestral (HG vs EF) populations

To enable comparative analyses of Transposable Element (TE) family content between the ancestral populations we first retrievered a curated primate TE library from Dfam^174^. Non-TE sequences (i.e. satellites) and elements with “unknown” classification were removed, based on the Dfam sequence identifiers. The final reference library (TE lib) contained 1595 curated TE sequences, each representing a distinct TE family from multiple primate species. Also, to enable mapping we generated various files for this final TE library. First, we retrieved the .fai index file with samtools faidx (env:eager_env). Second, we used picard CreateSequenceDictionary (env:perSVade_env_picard_env) to obtain sequence dictionaries from the genome fasta file. Third, we ran bwa index (env:perSVade_env) to index the genome for bwa-based mapping.

Then, trimmed sequencing reads from all the HG and EF individuals were mapped against the TE library using a pipeline that is equivalent to the one used for the hg37 reference (defined above). For this, we ran the script map_reads_TEs.py on the trimmed 500,000 read chunks of each individual (defined above). Specifically, this script first runs on each chunk the eager nextflow pipeline (env:eager_env) for i) read mapping with bwa aln (arguments -mapper bwaaln --bwaalnn 0.01 --bwaalnl 16500 --bwaalno 2), and ii) filtering out reads with mapping quality < 30 (arguments --run_bam_filtering --bam_mapping_quality_threshold 30). Then, it merges all mapped reads with samtools view (env:eager_env), as implemented in IGfun.get_merged_bam_from_list. Finally, map_reads_TEs.py runs IGfun.generate_bam_one_chrom_dedupped_and_long_reads_corrected to i) discard reads longer than 90 bp with samtools and awk (env:eager_env) (as they could be modern human contaminants), ii) remove duplicates with eager (argument --dedupper markduplicates) (env:eager_env), and iii) calculate damage statistics with eager (arguments ‘--damage_calculation_tool mapdamage --mapdamage_downsample 1000000’) (env:eager_env). The resulting filtered and de-dedupped reads were further used for TE abundance estimation. Also, the damage statistics were used for validation of the mapping (**Extended Data Fig. 18, 19**)

To retain only TE families with sufficient abundance across samples, we applied three filtering criteria: i) the presence of at least four mapped reads (n=4), ii) covering at least the 50% of the TE sequence length (l=50), and iii) found in at least the 50% of individuals within the dataset (i=0.5). These thresholds were estimated empirically after parameter testing across multiple combinations (see below; **Extended Data Fig. 20c**). Because sequencing depth varied substantially among TE families due to differences in length, raw read counts were normalized using transcripts per million (TPM). The resulting normalized abundance matrices, hereafter referred to TE abundances, were subsequently compared between population groups (HG vs EF) using the Mann-Whitney U test using the “mannwhitneyu” function with alternative=’two-sided’ from the scipy library (v1.15.2). Resulting p-values were corrected for multiple testing using the Bonferroni post hoc method using the “multipletests” function with method=“bonferroni” from the library statsmodels.stats.multitest. Finally, multiple visualizations were generated to explore and summarize patterns in TE family abundance differences across populations. The complete pipeline, named AncienTE (implemented in AncienTE.py, and run on env:AncienTE_env), was developed in Python, using scipy for statistical analysis, matplotlib and seaborn for data visualization, and biopython for sequence handling. This script takes as input a directory containing mapped reads from all individuals and a CSV file with population-level metadata; and performs automated filtering, normalization of TE abundances, statistical testing, and data visualization.

For further validation, we constructed a human-specific TE library derived from Dfam by retaining only human entries. Redundancy was reduced using CD-HIT^175^ with an 80% sequence identity threshold and 80% alignment coverage^176^. For LTR retrotransposons, LTR and internal regions were merged to generate complete elements. This process resulted in a non-redundant library comprising 882 TE families. We then evaluated the performance of AncienTE (run on env:AncienTE_env) using two alternative reference libraries: the primate-based library and the human-specific library, with the aim of determining which better captured biologically meaningful signals. Each library was tested under two parameter configurations: a permissive setting (n = 1, l = 30, i = 0.3) and a more stringent setting (n = 4, l = 50, i = 0.5; **Extended Data Fig. 20d**). Based on empirical performance, the primate-based library was selected for all analyses.

#### TE adaptive admixture analysis

To infer adaptive admixture we investigated TE dynamics in each of the admixed populations (from neolithic Ireland, Germany and Portugal) (**Fig. 1g**). For this, we constructed an admixture-based expectation model, inspired by the Fadm calculations described above, using the average EF / HG ancestry proportions (HG_prop, EF_prop) calculated as explained in ‘F4 statistics broad ancestral profile modeling of publicly available and newly reported datasets’. For each TE family, in each population, the expected abundance was calculated as a weighted average of population means:

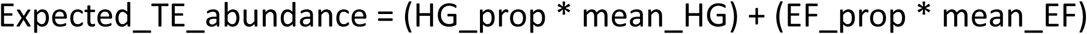

Observed TE family abundances were then estimated for each individual using map_reads_TEs.py and the initial steps (filtering and normalization) of the AncienTE pipeline (parameters n=4, l=50, i=0.5), and analysed separately by population (Germany, Ireland and Portugal). TE abundances were first transformed by log(1 - x) and then normalized using TE-wise min–max scaling, which rescales values across individuals to the [0,1] range. For each TE family, observed and expected abundances were compared using two-sided *t*-tests as implemented in “ttest_rel” from scipy (env:AncienTE_env), with multiple testing accounted for by Bonferroni correction as implemented in “multipletests” with method=“bonferroni” from Scipy (env:AncienTE_env). TE families showing significantly (bonferroni-corrected p<0.05) higher or lower abundances than predicted by the admixture model were thus identified. All these analyses were performed using the custom script admixed_analysis.py (run on env:AncienTE_env).

#### AncienTE validation using negative controls

To ensure that the variations observed by AncienTE were not driven by low or uneven coverage, post-mortem damage, or other technical artifacts, we analyzed two different negative control datasets matching HG / EF populations. First, the simulated LEPE48 chromosome 19 short-reads described at the section “Sample-tailored simulations of genomic datasets” were re-utilized, to test if AncienTE erroneously detects significantly different TE abundances between populations. Since all simulated reads originated from an identical genomic background, no genuine differences were expected. The chromosome 19 was deemed adequate for this negative control validation because of its relatively small size, its high TE proportion, and a TE proportion and copy number distribution that closely mirrors that of the remaining chromosomes in the human genome at the order classification level (**Extended Data Fig. 20a,b**). Accordingly, AncienTE should not detect statistically significant differences among TE families under this condition. To assess this, we ran map_reads_TEs.py and AncienTE on these simulated datasets when using different n,l,i parameters, and evaluated the number of false positive differences found (**Extended Data Fig. 20c**).

Second, we evaluated whether the pipeline might spuriously detect signals from non-repetitive genomic sequences (e.g. single-copy genes) in the real HG / EF samples. To this end, we constructed a control library consisting of 100 randomly selected single-copy gene sequences. Candidate genes were identified by querying the Ensembl REST API^177^ for protein-coding genes annotated as lacking paralogs in the human genome (has_paralogues = False). Then, we removed all genes categorized as pseudogenes or RNA genes. We further ensured that none of the selected genes overlapped genomic regions with significant CNVs identified in this study (**Supplementary Table 2**, column ‘mapped_Gene’). The final gene library was composed of 65 genes and can be found at **Supplementary Table 4**. Finally, the complete map_reads_TEs.py + AncienTE pipeline was executed using different n,l,i parameters, taking as a reference the single-copy-gene library instead of the TE library, and evaluated the number of false positive differences found (**Extended Data Fig. 20c**).

#### Statistical analyses on TE abundances

We assessed the association between TE abundance and genetic ancestry using multivariable linear regression on all individuals (HG, EF and admixed from the three populations). TE abundances (rows: TEs; columns: individuals) were log-transformed using a log(1 + x) transformation to reduce skewness and stabilize variance. For each TE, we fitted an ordinary least squares model of the form:

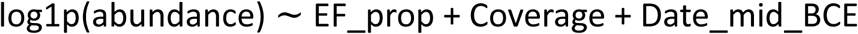

where EF_prop denotes EF ancestry proportion for each individual, Coverage accounts for its mean autosomal coverage (defined above), and Date_mid_BCE represents the midpoint of the calibrated chronological range. This approach allowed us to estimate the ancestry–abundance association while controlling for technical and temporal covariates. Regression coefficients, p-values, sample size and R² were extracted for each TE family. For visualization, predicted values were generated across the full EF_prop range (0–1), holding covariates at their median values, and plotted with 95% confidence intervals. Observed data points were coloured by population group (HG, EF, admixed). Analyses were performed in Python using NumPy, Pandas and Statsmodels (on env:AncienTE_env).

#### Validation of ancient TE signal

To validate that the reads mapping to the TE library are ancient we evaluated their MapDamage PMD profiles, as provided by map_reads_TEs.py. We found that all HG / EF mappings to the TE library had an aDNA-like PMD profile, but with a higher 5’ C>T mutation rate as compared to the mappings to the GRCh37 reference (**Extended Data Fig. 18**). We hypothesized that this may be due to the higher rate of variation within TE sequences, and to prove it we validated that the chromosome 19 simulated reads mapped to the TE library showed a similar PMD profile as the real samples (**Extended Data Fig. 19**). These findings are consistent with our expectation that reads mapped to the TE library are mostly ancient, and that we can use our datasets to infer changes in TE composition across populations.

## CONSORTIA INFORMATION

The Lepenski Vir consortium was integrated by the following authors: Aleksandra Žegarac^1^, Aleksandar Urošević^2^, Camille de Becdeliévre^3^, Quentin Cosnefroy^4^, Bojan Petrović^5^, Milica Šipovac^5^, Jelena Marković^6^, Marija Krečković Gavrilović^6^, Mihailo Radinović^6^, Marija Marin^6^, Mina Amzirkov^6^

1. Institute Biosens, University of Novi Sad
2. Institute for Biological Research “Siniša Stanković”, University of Belgrade
3. The Hebrew University, Department of Archeology
4. UMR 5199 PACEA
5. Medical faculty, University of Novi Sad
6. Laboratory for Bioarchaeology, Department of Archaeology, Faculty of Philosophy, University of Belgrade

## DATA AVAILABILITY

All of the sequencing datasets generated here are available under project code PRJNA1482402 in SRA. Also, **Supplementary Table 1** contains all individuals analyzed, with their relevant metadata. Furthermore, **Supplementary Table 2** includes all the selection signatures described in this work.

## CODE AVAILABILITY

All of the code and software environments used to generate the datasets, results, tables and figures presented here are in https://github.com/Gabaldonlab/MesoNeoSelection.

## ACKNOWLEDGEMENTS

This work was partly supported by the project EASI-Genomics – The European Union’s Horizon 2020 research and innovation programme under grant agreement No 824110. TG group acknowledges support from the Spanish Ministry of Science and Innovation, grant number PID2021-126067NB-I00 founded by MICIU/AEI /10.13039/501100011033 and by FEDER, UE and grand number PLEC2023-010225 founded by MICIU/AEI /10.13039/501100011033, as well as support from “La Caixa” foundation (grant number LCF/PR/HR21/00737, CI23-20260, LCF/PR/HR21/52410006 and LCF/PR/GN18/50310010); Fundació La Marató de TV3 (grant number 202328-31); AECC (grants PRYGN234923GABA and 290059); Instituto de Salud Carlos III (CIBERINFEC CB21/13/00061- ISCIII-SGEFI/ERDF and DTS25/00141); European Commission, Horizon Europe-HORIZON-MSCA-2023-DN-01-01 (grant number 101168618) and European Union’s Horizon 2020 research and innovation programme under the Marie Skłodowska-Curie grant agreement N° 101226544 (grant number 101227078). MAST and SOA acknowledge their AI4S fellowship within the “Generación D” initiative by Red.es, Ministerio para la Transformación Digital y de la Función Pública, for talent attraction (C005/24-ED CV1), funded by NextGenerationEU through PRTR. PC acknowledges FPI-2019 (MCIN, BDNS-ID:476421) grant and Max Planck Gesellschaft international postdoc scholarship. MGr acknowledges funding from NSERC Discovery grant (Canada) RGPIN-2018-05989. We thank Dr. Ainash Childebayeva for her feedback regarding the analysis of selection during the Mesolithic-Neolithic transition. We thank the members of the Population Genomics meeting in the Department of Archaeogenetics (Max Planck) for their feedback and evaluation of the work.

## AUTHOR CONTRIBUTIONS

MGr, CLF and TG, MAST and PC conceived and designed the study. MGr, CLF, IGG and TG secured funding. MAST, TG and PC wrote the first manuscript draft with contributions from other authors. All authors read, contributed to, and approved the final version of the manuscript. IGG, MGr, and PC assembled archaeological material and prepared the sample description. SS participated in conceiving, planning, and executing the archaeological research. MP performed morphometric analysis and archaeological description. AI performed morphometric analysis. KT, HK, CLB and MGu performed DNA extraction and library preparation. MGu and IGG planned and coordinated sequencing strategy. VZ aided in functional variant analysis. PC performed initial data quality and authenticity assessment for newly reported individuals, population genetic analyses and classification. SOA performed Transposable Elements analyses. PC and AAV performed ancient pathogen analyses. MAST performed all remaining bioinformatic analyses (integration of the whole dataset, selection scans, functional annotation, enrichments and Copy Number Variant analyses)., TG coordinated and supervised computational analyses. The members of the Lepenski Vir consortium performed morphometric analysis, (AŽ, AU, CdeB, QC, BP, MŠ, JM, MKG, MR, MM, MA), archaeological description (CdeB, QC, BP, MŠ, JM, MKG, MR, MM, MA), and aided in DNA extraction and library preparation (AŽ)

## SUPPLEMENTARY TABLES

**Supplementary Table 1**. **Metadata summary for all ancient individuals included in this study.** This table includes the summary metadata of 141 publicly available and 11 newly sequenced individuals (see column ‘source_data’) for this study. The columns including ‘qpAdm’ refer to the ancestry proportion measurements (see **Methods**). Most importantly, ‘qpAdm_proportion_EF’, ‘qpAdm_proportion_EHG’ and ‘qpAdm_proportion_WHG’ refer to the final assigned proportions of EF / WHG / EHG, from which the ‘final_profile’ (pure HG, pure EF or admixed) was inferred. Also, ‘source_qpAdm_proportions’ refers to the analysis from which these proportions were inferred, which may be ‘all_samples’ or ‘our_samples’. For source_qpAdm_proportions = ‘all_samples’, the proportions were inferred from the global qpAdm analysis applied to all samples (see **Methods**), which are reported in the columns starting with ‘qpAdm_all’. Note that for some of the samples sequenced here we could not fit such a model (see **Methods**), so the ‘qpAdm_all’ are blank. Conversely, for source_qpAdm_proportions = ‘our_samples’, the proportions were inferred from a qpAdm analysis tailored to the newly sequenced samples (see **Methods**), which are which are reported in the columns starting with ‘qpAdm_our’. Beyond the qpAdm results, the table also shows the inferred haplogroups (for mtDNA and y chromosome), the number of genotype SNPs in the 1240k array, the dating information (columns ‘Date_BCE’ and ‘RC_dating_ID’), archaeological information (‘FEMFERT_ID’, ‘Grave_ID’, ‘archeological_age’, ‘stature’, ‘body_mass’ and ‘BMI’), the source skeletal element, the publication information (year, publication abbreviation and DOI), the group ID, geographical origin (locality, latitude, longitude and present-day political entity), the sequencing source (either the sequencing project ID for samples sequenced here or the ENA run IDs for the AADR ones), the % of modern human contamination as inferred by ContamMix, the Library type (minus = no damage correction, half = damage retained at last position, plus = damage fully corrected, ds = double-stranded library preparation, ss = single-stranded library preparation), the mean autosomal depth of coverage (calculated across mappable windows, see **Methods**), the percent of the autosomal genome covered (only for mappable windows), the chromosomal sex, the admixed population (Ireland, Portugal, Germany for nIre, nPor, nGer, respectively), the downsampling ID (used for generating the ancient imputation panel, see **Methods** and **Extended Data Fig. 9**), and sequencing metadata for the newly-generated datasets (‘P7 Index Seq.’, ‘P5 Index Seq.’, ‘# Indexing PCR Cycles’, ‘Qubit (ng/μl)’). We obtained all the related AADR metadata from the file ‘AADR_v62.0_1240k_public.anno’ available at https://dataverse.harvard.edu/dataset.xhtml?persistentId=doi:10.7910/DVN/FFIDCW.

**Supplementary Table 2. Functional information for the top, non-redundant hits under selection, including SNPs and CNVs.** Each row includes a functional effect of a given variant. The columns ‘#CHROM’, ‘POS’, ‘CNV_start’, ‘CNV_end’, ‘ID’, ‘REF’ and ‘ALT’ indicate the variant coordinates with respect to GRCh37, used for calling and imputation. Also, ‘ID_GRCh38’ reflects the lifted coordinates to GRCh38, which was used for variant annotation. The ‘type_top_hit’ indicates the source of the hit, which may be CNV_Fst (CNVs with highly differentiated Fst between EF and HG populations), CNV_Fadm-Germany (CNV with reduced AF in nGer population, see **Fig. 4 h**), pcadapt (PCAdapt scan top hit), XPEHH (XP-EHH scan top hit), Fadm_<admixed population> (Fadm scan top hit) or iDAT_<admixed population> (iDAT scan top hit). Additionally, ‘direction_top_hit’ indicates the direction of selection (e.g. favouring HG or EF alleles). The ‘sig_methods_SNPs’ indicates which SNP selection scans yielded this SNP as significant, even if it did not end in the final set of non-redundant top hits. The columns ‘AF_<POPULATION>‘ indicate the observed allele frequency of the variant in each population, while the ‘AF_admixed_expected_<admixed population>’ indicates the expected AF under the mixture HG+EF model. Furthermore, there table includes the statistics (Fadm, iDAT, XP-EHH, fraction of EF/HG LATs) and p-values (also including bonferroni-corrected ones) of all selection scans (see **Methods**). Regarding functional variant annotations, the table includes the dbSNP ID, the mapped gene involved in the functional effect, the source of the variant-gene mapping (column ‘mapped_Gene_type’, which can be the nearest one (Nearest_Gene), based on eQTL annotations (eQTL_Gene) or provided by VEP (Gene)), the DNA / protein effect metadata provided by VEP (BIOTYPE, Consequence, cDNA_position, CDS_position, Codons, Protein_position, Amino_acids), predicted functional effects by VEP (from SIFT, PolyPhen, ClinVar and various databases included in the VEP ‘PHENOTYPES’ plugin), eQTL annotations (see **Methods**), hits in the GWAS catalog, and G:Profiler gene description and pathway annotations. Finally, the ‘type_genes_enrichment’ indicates the belonging of each annotation to a gene set used for functional enrichments (see **Fig. 7**), and ‘type_genes_enrichment_description’ provides a description of the gene set.

**Supplementary Table 3. Table with the regions under selection in previous studies.** Each row includes one of these regions, with the ‘study’ indicating the source of the signal. Also, the ‘favoured_allele’ indicates the directionality of selection.

**Supplementary Table 4. Single copy genes used for the negative control of AncienTE.** In total 65 single copy genes were selected as a negative control to test AncienTE performance (see **Methods**).

**Supplementary Table 5. Number of significantly changed TE families, grouped by superfamily, identified in admixed individuals in each population.**

**Supplementary Table 6. Functions enriched in the different gene sets**. The ‘non_redundant’ column indicates whether the enriched function was included among the final set of non redundant ones from **Fig. 7d**. The ‘type_genes’ column indicates the gene set where the enrichment was performed, as defined in **Supplementary Table 2.**

