## Supplementary information for "Ancient whole genomes reveal regional selection during adaptation to Neolithic lifestyle in Western Eurasia"

#### Table of contents

**1. Archaeological Information on the Newly Reported Individuals** **2**

1.1 Vlasac 2

1.2 Lepenski Vir 7

1.3 Outside the Danube Gorges: Golokut-Vizić 13

**2. Methodology on the Statistical Framework for Population Genomics tests** **18**

2.1 Proximal Demographic Modeling of Newly Reported Individuals 18

2.2 Distal Demographic Modeling of Publicly Available Individuals 21

**3. Extended Results and Discussion** **24**

3.1 Exploring the Feasibility of SNP calling 24

3.2 SNP-based Selection Scans 25

3.3 Transposable Element Analysis 28

**Supplementary References** **31**

### 1. Archaeological Information on the Newly Reported Individuals

Sites in the Danube Gorges region, occupied between the 10th and 6th millennia BC, document the Mesolithic–Neolithic transition in the Central Balkans. During the Mesolithic, fisher-forager communities established settlements along the riverbanks and produced elaborate architecture and distinctive forms of art. At the onset of the Neolithic in this region (from around 6200 BC), new populations reached the Danube Gorges. Many sites were located near whirlpools, which would have facilitated specialized fishing, particularly of large species [1–3].

Among wild game, red deer remains are the most numerous, followed by wild boar, roe deer, and aurochs[4]. Large game (especially red deer and aurochs) constituted an important component of the diet due to the substantial amount of meat they provided[5]. Fish also represented a significant dietary component, with remains of various cyprinids (*Cyprinidae*), Wels catfish, migratory sturgeons (*Acipenseridae*), and huchen (*Hucho hucho*) recovered from Lepenski Vir, Padina, and Vlasac. It is important to emphasize that the large size of some species, such as beluga sturgeon and catfish, would have provided a considerable amount of protein [6].

Regarding domesticated animals, remains of dog, pig, cattle, and sheep/goat have also been identified in the Danube Gorges. Stable isotope analyses confirm the significant role of fish in the diet, as well as an increase in the consumption of terrestrial resources at Lepenski Vir during the Neolithic period[7]. Below we provide a description of the specific archaeological sites and of the individual remains from which new genomic sequences were obtained in this study.

#### 1.1 Vlasac

Vlasac is situated in the Upper Gorge, on the right bank of the Danube River. Absolute dates for Vlasac range from 7035–6698 cal BC to 5700–5500 cal BC[8]. The first excavations at Vlasac were conducted by D. Srejović and Z. Letica in 1970–1971[9]. An area of 640 m² was investigated, yielding various findings from the Mesolithic, Neolithic, and Eneolithic periods. Subsequently, the site was flooded following the construction of the Đerdap I dam, which created a large water reservoir. However, in 2006, the water level dropped, exposing a section of the riverbank and enabling new excavations at the site[8]. We sequenced the genomes for six individuals from this site, detailed below.

**1.1.1 Vlasac, grave 29, female, around 30. Genetic Identifier: i723-i515**

Absolute chronology: BRAMS-2578, corrected for the reservoir effect, 7046–6691 cal BC [10].

Description: The skeleton was found in square A/II, at a relative depth of 1.25–1.87 m, with an orientation of south/southwest–north/northeast (head at 205°). A larger parallelepiped-shaped stone was found on the rib cage, in front of the skull, near which "hearth 3" was identified. The individual was buried in a supine position, with the lower part of the skeleton slightly turned to the right side. The pelvis was dislocated; the arms and hands were positioned in the pelvic area, while the legs were slightly flexed. Ochre was scattered over the pelvis, femora, and hands. In the pelvic area, teeth of cyprinids were found. Regarding grave goods, two fragmented tools made of deer antler were recovered.

Relative chronology: Late Mesolithic

Age: Histological study of tooth cementum annulation indicates an age of 28.72 years.

Stature: 172.71 cm

Body mass: 63.50 kg

Body mass index: 21.29 (normal)

Pathology: On the right femur, in the medial third of the shaft, near the medial lip of the linea aspera, a swelling of inflammatory origin was observed (15 × 8 mm in size and 8 mm in thickness). At its center, a fistula-like opening is present (diagnosis: status post osteomyelitis of the left femur). *Cribra orbitalia* was noted on the left orbital roof (degree 2, activity 3) and on the right orbital roof (degree 1, activity 1). Porotic hyperostosis was observed on the frontal bone (degree 1, activity 2), parietal bones (degree 2, activity 3), and occipital bone (degree 2, activity 3). A periosteal reaction was noted on the medial surfaces of both tibiae.

Dental analysis: Supragingival dental calculus (degree 1) was observed on four teeth.

Radiogenic strontium analysis: Local origin[11].

Stable isotope analysis: Not applicable.

Other observations: The epigenetic feature *perforatio fossa olecran*i was noted on the right humerus.

**1.1.2 Vlasac, grave 55, female, around 40 years. Genetic Identifier: i705-i505**

Absolute chronology: BRAMS-2583, corrected for the reservoir effect, 7531–7348 cal BC [12].

Description: The skeleton was found in square A/17, in sterile soil at a relative depth of 2.56 m (head at 50°), with a northeast–southwest orientation. The individual was buried in an extended supine position, with the hands placed on the lower abdomen. The feet of the individual were located beneath grave 54a.

Relative chronology: Late Mesolithic

Age: Histological study of tooth cementum annulation indicates an age of 42.61 years.

Stature: 166.54 cm

Body mass: 65.81 kg

Body mass index: 23.73 (normal)

Pathology: Porotic hyperostosis was observed on the frontal bone (degree 1, activity 2), parietal bones (degree 3, activity 3), and occipital bone (degree 2, activity 2). On the right side of the frontal bone, 31 mm from the coronal suture, a very shallow (0.01 mm) egg-shaped depression, measuring 17 mm in length and 12 mm in width, was noted. On the medial surfaces of both tibiae, a moderate periosteal reaction with some remodeling was observed. A periosteal reaction was also visible on the lateral surfaces of both tibiae, but to a lesser extent. A mild periosteal reaction with traces of remodeling was also noted on both femora.

Dental analysis: Supragingival dental calculus (degree 1) was observed on eight teeth, and degree 2 on three teeth. One tooth was lost *ante mortem.*

Radiogenic strontium analysis: Local origin[11].

Stable isotope analysis: Not applicable.

Other observations: The epigenetic feature *perforatio fossa olecrani* was noted on the left humerus.

**1.1.3 Vlasac, grave 56, female 40-50. Genetic Identifier: i706-i506**

Absolute chronology: BRAMS-2584, corrected for the reservoir effect, 7135–6700 cal BC [12].

Description: The skeleton was found in square D/5, at a relative depth of 2.24 m, with a west-southwest–east-southeast orientation (head at 240°). The individual was laid on her right side, with the head tilted backward. The arms were positioned alongside the body, with the hands placed on the lower abdomen. The right leg was placed over the left.

Relative chronology: Late Mesolithic

Age: Histological study of tooth cementum annulation indicates an age of 47.39 years.

Stature: Not applicable.

Pathology: On the atlas, the superior articular facet was found to be considerably enlarged and bordered by an annular osteophyte approximately 1–2 mm in height (diagnosis: spondylarthrosis atlanto-occipitalis l.s. (++)). Porotic hyperostosis was observed on the frontal bone (degree 1, activity 2), parietal bones (degree 3, activity 2), and the occipital bone (degree 2, activity 2).

Dental analysis: Supragingival dental calculus (degree 1) was observed on one tooth. One tooth was lost *ante mortem.*

**1.1.4 Vlasac, grave 67, middle aged adult. Genetic Identifier: i722-i513**

Absolute chronology: BRAMS-2585, corrected for the reservoir effect, 7319–6828 cal BC [10].

Description: The skeleton was found in square C/9, at a relative depth of 2.60 m, with a south–north orientation (head at 190°). The grave was located 0.40 m beneath hearthstone no. 23. The individual was buried in an extended position, with the legs slightly flexed to the left and the arms placed alongside the body, while the right hand rested on the lower abdomen. In the pelvic area, fetal bones were found. Ochre was present in the pelvic region. A stone was found above the head.

Relative chronology: Late Mesolithic

Stature: 163.7 cm

Pathology: Cribra orbitalia was observed on the right orbital roof (degree 2, activity 3). Porotic hyperostosis was present on the frontal bone (degree 1, activity 3), parietal bones (degree 3, activity 3), and occipital bone (degree 2, activity 3). On the right side of the frontal bone, 21 mm from the coronal suture, a small egg-shaped lesion (8 mm in length, 5 mm in width, and 0.2 mm in depth) was noted. Additionally, on the left side of the frontal bone, a 19 mm area of pronounced porotic hyperostosis was observed. On the parietal bones, three lithic fragments were found.

Dental analysis: Supragingival dental calculus (degree 1) was observed on four teeth.

**1.1.5 Vlasac, grave 74, female, around 50. Genetic Identifier: i707-i507**

Absolute chronology: BRAMS-2587, corrected for the reservoir effect, 7312–7057 cal BC [12].

Description: The skeleton was found in square E/9, at a relative depth of 1.40 m, with an east–west orientation (head at 90°). The grave was located approximately 0.50 m beneath a stone placed next to grave no. 71. The individual was buried in an extended supine position, with the arms positioned alongside the body. The right hand was placed on the lower abdomen, while the left rested on the left hip. In the pelvic area, teeth of cyprinids were found.

Relative chronology: Late Mesolithic

Age: Histological study of tooth cementum annulation indicates an age of 48.92 years.

Stature: Not applicable.

Pathology: On the proximal epiphysis of the fifth right (?) metatarsal, a bill-like bone apposition (approximately 3–5 mm) was observed (diagnosis: *arthrosis deformans articulationis tarso-metatarsalis quinti l.d.* (?) (++), osteomalacia (+)). Cribra orbitalia was noted on both orbital roofs (degree 1, activity 2). Porotic hyperostosis was observed on the frontal bone (degree 1, activity 2), parietal bones (degree 2, activity 2), and occipital bone (degree 3, activity 2).

Dental analysis: Dental calculus (degree 1) was observed on nine teeth, and degree 2 on two teeth (both supragingival and subgingival).

Radiogenic strontium analysis: Not applicable.

**1.1.6 Vlasac, grave 82, female, around 40. Genetic Identifier: i716-i512**

Absolute chronology: BRAMS-2588, corrected for the reservoir effect, 6589–6273 cal BC [10].

Description: In square D/15, at a relative depth of 2.27–2.30 m, four skulls were discovered (82, 82a, 82b, 82c). Two skulls with dislocated mandibles (82, 82a) were enclosed by two randomly placed limestone slabs, with long bones of the postcranial skeleton placed both above and below them. Further east, additional skulls (82b, 82c) were found, positioned atop long bones.

Relative chronology: Late Mesolithic

Stature: 164.60 cm

Body mass: 62.79 kg

Body mass index: 23.17 (normal)

Pathology: Porotic hyperostosis was observed on the frontal bone (degree 2, activity 2), parietal bones (degree 3, activity 2), and occipital bone (degree 2, activity 2).

Dental analysis: Supragingival dental calculus (degree 1) was noted on five teeth.

Radiogenic strontium analysis: Local origin.

##### 1.2 Lepenski Vir

The site of Lepenski Vir is situated in the Upper Gorge, between the right bank of the Danube River and the slope of Koršo Hill, near the confluence of the Boljetin River. This area of the Gorges is difficult to access due to the steep hills (up to 500 m) and the narrowing of the Danube. Lepenski Vir is located between the sites of Padina and Vlasac. The site was discovered in the 1960s, prior to the construction of a hydroelectric dam in the Danube Gorges. Excavations were conducted by D. Srejović from 1965 to 1970[13–15]. An area of 2,500 $m^2$ was excavated, with archaeological layers averaging 3.5 m in depth[6].

The stratigraphy of Lepenski Vir is very well documented, as it is among the best radiocarbon-dated sites of the period in the region. These dates indicate that the site was inhabited from the Mesolithic through the Early Neolithic, as well as during the Eneolithic, Roman, and Medieval periods[6,16]. We generated shotgun whole genomes for four individuals from this site, detailed below.

###### 1.2.1 Lepenski Vir, grave 8, female, 30–40 years. Genetic Identifier: i701-i501

Context: Square B/IV, corner D of Building 24, approximately 0.30 m above the floor.

Description: The skeleton is oriented south–north with a 55° deviation west of north. The head is positioned to the south, facing east. The individual was placed in a flexed position on the right side, with the knees drawn to chest height. The arms were bent, with the hands positioned in front of the face (diary 06/08/1967).

Relative dating: Middle Neolithic

Absolute dating: OxA-25207 (replacing OxA-11694), corrected for the reservoir effect, 5986–5783 cal BC (Bonsall et al. 2015).

Stable isotopes: $\delta^{15}N$ = 9.4‰, $\delta^{13}C$ = –20‰, $\delta^{34}S$ = 5.5‰ (de Becdelièvre 2020). These values suggest a diet largely based on terrestrial proteins, primarily from herbivores and C3 plants (LV 8, LV 88, LV 32a, LV 17, lower-left quadrant of PCA). Notably, starch grains from a Poaceae species were detected in the dental calculus of non-local individuals LV 8, LV 20, and LV 32a (Cristiani et al. 2016).

Bones present:

- Cranial skeleton: Damaged frontal bone, right sphenoid, partially damaged left parietal, damaged temporal and occipital bones. Maxillofacial bones are preserved from below the lower nasal edge, including the entire palatal surface. The right zygomatic bone is preserved, with only a narrow fragment of the left along the orbit. The mandible is complete. In the maxilla, only tooth 12 is missing ante mortem; in the mandible, teeth 41–44 and 31–34 are absent.
- Postcranial skeleton: Right humerus, left ulna and radius, both clavicles, fragmented scapulae, most hand bones, right pelvic wing, midshaft-damaged right femur, left tibia, head of the left femur, right and left fibulae (without proximal epiphysis), and individual foot bones.

Sex: Female, with pronounced sexual characteristics. The greater sciatic notch is very wide, and the preauricular groove is strongly emphasized. Gracility of postcranial bones and overall cranial morphology (gracile skull, non-prominent brow ridges, very small mastoid processes, absent external occipital protuberance) further confirm female sex.

Age: Estimated between 30–40 years. Teeth attrition (except for anterior maxillary teeth affected by activity) suggests the individual was no older than 40. Auricular surface morphology indicates

35–40 years, whereas cranial suture closure suggests around 40. Epiphyseal lines on the femoral heads suggest 25–30 years. Porous hyperostosis on the femoral heads may reflect prolonged growth due to health disorders. Overall, the age is estimated to be 30–40 years.

Musculoskeletal markers of stress: This individual exhibits unique stress markers among Mesolithic–Neolithic women of the Djerdap region. The M. latissimus dorsi attachment on the humerus is unusually pronounced, suggesting activities not observed in other individuals. Abrasion of anterior maxillary teeth indicates they were likely used as tools. Both humeral and dental markers may result from the same activity. Additionally, a 2 cm-wide depression along both parietal bones near the coronal suture is consistent with load-carrying using straps over the head.

Pathology: Strong traces of porous hyperostosis are present on the femoral necks. No cranial infections were observed. The left tibia shows minor remodeled periosteal reaction (OB1/1).

Cranial metrics: Maximum skull length 18.6 cm, maximum width 13.3 cm, cranial index 71.50, classifying the skull as hyperdolichocranial—the only such skull at Lepenski Vir.

Body height and weight: Based on humeral maximum length (32.1 cm), estimated height is 167.3 cm; based on femoral length (44.1 cm), 165.7 cm. Average height is 166.33 cm. Body weight, estimated from maximum femoral head diameter (4.2 cm), is 54.1 kg.

**1.2.2 Lepenski Vir, grave 54d, female, 40-50 years. Genetic Identifier: i702-i502**

Context: Square B/12, IX–X o.s., within stone structure XXXVI (61.543 m).

Description: The skeleton was displaced by the burial of grave 54e. Skeletal remains of grave 54d are located on both sides of skeleton 54e. The skull is located near the middle of the skeleton. The individual was laid in an extended position, oriented south–north with a 10° deviation east of south. The skeleton is at a greater depth than skeletons 54b and 54c and at the same level as skeleton 54e. A single bone awl was found as a grave good.

Relative chronology: Mesolithic–Neolithic transition

Absolute chronology: OxA-25213 (replacing OxA-11700), corrected for the reservoir effect, 6351–6013 cal BC[17].

Bones present:

- Cranial: Almost the entire frontal bone is preserved; parietals lack small fragments; occipital bone lacks the basal fragment; temporal bones partially damaged. Facial bones are highly fragmented, with only a small fragment of the maxilla preserved; the mandible is intact. In the maxilla, teeth 12, 16–18, 23–24, and 26–28 are present, while 13–15, 22, and 25 are lost ante mortem; tooth 21 is missing with part of the jaw. In the mandible, teeth 35–36, 42–43, and 45–47 are present; the rest are lost postmortem.
- Postcranial: Left and right clavicles (right without medial end), fragment of upper sternum, acromion of right scapula, 28 rib fragments, upper third of sacrum, left and right ilium with fragments of both sciatic notches, both humeri (proximal ends damaged), left ulna and radius (extremities damaged). Hand bones include 2 carpals, 2 metacarpals, 6 proximal phalanges, and 1 medial phalanx. Vertebral column includes 2 cervical, 1 thoracic, and 3 vertebral extensions. Lower limbs include left femur (missing small fragment of head), both tibiae (proximal ends damaged), left patella, right fibula (proximal end damaged), fragment of left fibula, left calcaneus, right calcaneus (missing lateral half), 2 cuneiforms, 1 cuboid, right navicular, and 5 metatarsals.

Sex: Female. Despite fragmented pelvic bones, the wide greater sciatic notch and strongly pronounced preauricular sulcus (the largest at Lepenski Vir) confirm female sex. Cranial morphology is very gracile, with unaccented supraorbital arches, parietal and frontal tubers, small mastoid processes, and frontal bone inclination supporting female characteristics. The gracile mandible with unpronounced mental protuberance, as well as femoral metrics, further confirm female sex.

Age: Estimated 45 years. High tooth wear is observed; maxillary second and third molars suggest the individual was not older than 50. Cranial suture closure suggests ~45 years, while auricular surface fragments indicate 40–45 years. Histological study of tooth cementum annulation indicates 48.89 years.

Body height: Based on maximum femoral length (42.5 cm), estimated height is 161.03 cm.

Cranial metrics: Maximum skull length 17.9 cm, maximum width 13.2 cm, cranial index 73.74, classifying the skull as long and narrow. Mandible: chin height 3.42 cm, mandibular body height 3.02 cm, body width 1.1 cm, bigonial width 9.1 cm, minimum ramus width 3.5 cm, maximum ramus width 4.5 cm.

Stable isotopes: $\delta^{13}C$ = –19.9‰, $\delta^{15}N$ = 13.4‰, $\delta^{34}S$ = 13.8‰. Elevated $\delta^{34}S$ values suggest consumption of anadromous fish, while slightly lower $\delta^{13}C$ and $\delta^{15}N$ indicate inclusion of terrestrial proteins.

Comment: LV 54D (local) and LV 54E (non-local) were found in the same location over the abandoned space of trapezoidal building 65, dated 6340–6015 cal BC and 6210–5930 cal BC, respectively (Period of Transformation – Early Neolithic; Bonsall et al. 2015). LV 54D was displaced, likely by the deposition of LV 54E, who was laid extended on her back. Despite very similar $\delta^{13}C$ and $\delta^{15}N$ values (LV 54D: –19.9‰, 13.4‰; LV 54E: –19.7‰, 13.9‰), $\delta^{34}S$ of the local female LV 54D is higher (13.7‰ vs. 10‰). This may reflect environmental differences prior to migration, with LV 54E consuming more freshwater fish and terrestrial game. Their proximity and isotopic similarity suggest potential close social or familial bonds.

Pathology: Mild porous hyperostosis on parietal, temporal, and occipital bones, with inactive areas (e.g., right parietal). Fistula below mandibular teeth 46–47 measuring 0.9 × 0.6 cm.

Trauma: Two small impact traces on the right parietal bone, 4.2 cm and 8.1 cm from the lambdoid suture (length 0.7 cm × width 0.5 cm, and length 0.5 cm × width 0.1 cm), consistent with soft tissue coverage at time of injury; unclear if perimortem or postmortem. Possible animal claw traces observed on distal medial–posterior femur.

###### 1.2.3 Lepenski Vir, grave 96, newborn, 38–40 gestational weeks. Genetic Identifier: i703-i503

Context: Below the level of building 43, in an area without a floor, along the upper narrower side of the building. No grave cut was observed; the skeleton was located at an elevation of 65.52 m.

Description: Oriented southeast–northwest, with a 5° deviation from east toward south (140° from north). The head was directed to the southeast. The skeleton's position could not be fully determined due to damage. Preserved elements include fragments of the skull, limb bones, several vertebrae, and ribs (diary 05/09/1970).

Relative dating: Transition / Early Neolithic

Absolute chronology: BRAMS-4507, corrected for the reservoir effect, 6085–5894 cal BC (unpublished BIRTH project data)

Individual age: Based on maximum long bone lengths (femur 7.8 cm, humerus 6.9 cm), the individual was a newborn of 38–40 gestational weeks. Histological analysis confirmed the presence of a neonatal line, indicating live birth. Postnatal enamel measurements suggest the infant lived approximately 18 days after birth[18].

Pathology: Distal extremity thickening is present on the tibiae, with moderately pronounced periosteal reaction (OB2) on the medial surfaces of both tibiae. A milder periosteal reaction (OB1) is observed on the femurs and upper limb bones, while the proximal end of the humerus exhibits intense reaction; only the distal end is preserved on the left humerus.

**1.2.4 Lepenski Vir, grave 113, newborn, 38-40 gestational weeks. Genetic Identifier: i704-i504**

Context: Below the floor of building 63, at an elevation of 59.40 m.

Description: The tomb is rectangular, cut into the floor of building 63. The opening is partially covered with slab stone. The skeleton is well preserved, with the head oriented north–northwest (335° from north). The individual lies supine, with arms extended along the body, legs spread and bent at the knees so that the tibiae and fibulae lie parallel. Beneath the floor of the same building lies grave 117 (diary 30/09/1970).

Relative dating: Transition / Early Neolithic

Absolute chronology: Not applicable.

Bones present: The skeleton is largely complete. All bones of the upper and lower limbs are present, except for the bony pelvic wing; the scapulae are fragmented. The skull bones are highly fragmented but mostly preserved.

Individual age: Based on maximum long bone lengths (femur 7.7 cm, humerus 6.6 cm), the newborn was 38–40 gestational weeks.

Pathology: Both tibiae exhibit moderately pronounced periosteal reaction (OB2), with slightly milder reaction on the femurs and upper limb bones. Distal femoral epiphyses are notably expanded, and the proximal thirds of the tibiae show strong arcuate curvature. Thickening of the sternal end of the clavicle indicates possible congenital infection.

##### 1.3 Outside the Danube Gorges: Golokut-Vizić

The site of Golokut is located in a glade bordered by trees, at an altitude of 195–200 m asl, on the western slopes of Fruška Gora Mountain in the Srem region[19]. It lies 2 km southwest of the small village of Vizić, 5 km from Neštin, and 6.5 km from the Danube River. The site was discovered during an archaeological field survey in 1965, and shortly thereafter, in 1973, the Museum of Vojvodina initiated excavations under the direction of Jelka Petrović. Excavations continued, with minor interruptions, until 2003, encompassing more than 50 trenches. Two occupation layers have been identified, both corresponding to a single phase of the Starčevo culture. In addition to the Starčevo phase, layers associated with the Vučedol and Vinkovci cultures have also been documented[19–21].

Radiocarbon dating places the site roughly in the middle of the sixth millennium BC (5630–5470 cal BC to 5560–5360 cal BC)[22]. Within the Starčevo layer, several pits containing pottery and osteological remains were excavated in dwellings, alongside above-ground structures, hearths, ovens, and six graves.

Two middle-aged females from Golokut-Vizić exhibit elevated body mass indices. Notably, both displayed numerous dental and skeletal pathologies. The woman from grave 2/1984, who had a predominantly terrestrial diet, showed cribra orbitalia with healed and unhealed lesions, dental caries, enamel hypoplasia, and ante-mortem tooth loss. The woman from grave 3/1984 had a low-protein diet, evidence of starch grain consumption, cribra orbitalia with healed and unhealed lesions, dental caries, and enamel hypoplasia.

Given that all overweight individuals exhibited health issues, it is plausible that their dietary patterns and mobility strategies contributed to elevated body mass indices. We sequenced the genomes for four individuals from this site, detailed below.

###### 1.3.1 Golokut Vizić, grave 3 (1984), female, 30–39 years. Genetic Identifier: i713-i509

Context: South-eastern part of the pit in trench 30.

Description: The individual was buried in an extended supine position, oriented northwest–southeast, with the head toward the northwest. The right arm was placed above the chest, palm on the left side, with the hand positioned near the left elbow. The head was bent toward the left shoulder, resting on the left hand. Legs were extended side by side. One obsidian knife was found near the right side of the skull, but it likely does not belong to the grave goods.

Relative chronology: StarČevo culture, Early–Middle Neolithic

Absolute chronology: OxA-8505, 5621–5379 cal BC[22]

Preserved bones:

- Skull: Well preserved; mandible fragmented, missing both condyloid processes.
- Postcranial skeleton: Fragmented; preserved elements include left humerus (distal end partially), right humerus (partially fragmented), both ulnae, both radii, left clavicle, right clavicle (without sternal end), scapulae (both fragmented), coracoid and acromion processes, glenoid fossa, sternum fragment, cervical (5), thoracic (10), lumbar vertebrae (5), ribs (various fragments), left and right pelvic bones (fragmented), left femur (proximal end fragmented), right femur (proximal end + diaphysis), right tibia (partially), right fibula (proximal end), right talus, navicular, medial cuneiform, four left metatarsals, one proximal and one distal foot phalanx, left hand bones including carpal, metacarpals, and phalanges.

Sex: Female, based on gracile skull (unaccented nuchal crest, mastoid process, supraorbital margin, glabella, and mandible) and pelvic morphology.

Age: Estimated 30–39 years. Suture closure suggests ~40 years, tooth wear suggests 25–35 years, and pelvic bone characteristics confirm 30–39 years.

Dental analysis:

- All teeth present except 22, 31, 41 (lost postmortem).
- Abrasion: Degree 2 on most posterior teeth; degree 3 on 11, 23, 24; degree 4 on 12, 21.
- Hypoplasia: Teeth 13, 17, 18, 27, 28 (linear or pit).
- Caries: Teeth 26, 27.
- Supragingival calculus (degree 1) on teeth 27, 28 (buccal) and 32–38, 42–48 (bucco-lingual).

Pathology:

- Cribra orbitalia: Right orbit (degree 1, activity 2), left orbit (degree 2, activity 3).
- Porotic hyperostosis: Frontal (1/2), parietals (2/2), occipital (2/3).

- Mild periosteal reaction along the right tibial diaphysis.

Stable isotope analysis: $\delta^{13}C$ and $\delta^{15}N$ indicate a primarily terrestrial diet[22].

Starch grain analysis: Two starch grains identified in dental calculus.

Metric analysis: All measurements and indices shown in Table 1 of [23].

Stature and body mass: Not applicable.

Estimated stature: Humerus 32.0 cm: 165.49 cm; femur 43.6 cm: 161.79 cm; average 163.64 cm.

Body mass: Based on left femur head (4.6 cm): 68.84 kg.

Comment: This individual had very low $\delta^{15}N$ values, indicating minimal animal protein intake, which likely impacted her health. Childhood stress events (ages 3–5) are evident from linear enamel hypoplasia. Persistent adult health issues are confirmed by cribra orbitalia and partial lesion activity at death. Her primarily terrestrial, carbohydrate-rich diet (starch grains in calculus) may have contributed to dental caries. Similar dietary patterns are observed in other Golokut-Vizić individuals, suggesting a preference for cereals, which negatively affected health.

Other observations: A few animal bones were found among the human remains, including one cattle lower incisor, sheep/goat calcaneus, and two deer medial phalanges.

**1.3.2 Golokut Vizić, grave 3 (2003), juvenile, 15–18 years. Genetic Identifier: i708-i508**

Context: Trench 72, north-western part of pit-dwelling 27.

Description: The individual was buried in a semi-extended supine position, oriented southwest–northeast, with the head toward southwest and face oriented north. The right arm lay next to the body, while the left arm and left leg were contracted. The right leg was positioned above the left leg.

Relative chronology: Starčevo culture, Early–Middle Neolithic

Absolute chronology: BRAMS-4489, corrected for reservoir effect, 5481–5373 cal BC (unpublished BIRTH project date)

Preserved bones:

- Cranial: Left frontal, left parietal, 1/5 of right parietal, left half of squama occipitalis, fragmented temporal bones, fragment of greater wings of sphenoid, alveolar fragments of maxillae, left and right mandible halves (without condyloid processes).
- Postcranial: Both humeri (proximal epiphysis not fused), both ulnae (proximal epiphysis visible, distal not fused), both radii (proximal and distal epiphyses unfused, except distal left radius partially fused), left scaphoid, left lunate, left triquetral, left capitate, left hamate, right triquetral, right trapezium, right trapezoid, 9 fragmented metacarpals (distal ends not fused), 6 proximal, 3 medial, 2 distal phalanges (proximal ends of proximal phalanges not fused), medial half of left clavicle, right clavicle (without sternal end), lateral borders of scapulae, glenoid fossa, fragmented acromion, 6 cervical, 10 thoracic, 5 lumbar vertebrae, 13 vertebral + 4 sternal ends of ribs, 20 rib body fragments, left iliac, fragmented left ischium & pubis, fragmented right iliac & pubis, fragmented sacrum, both femurs (proximal partially fused, distal not fused), left tibia (epiphyses not fused), right tibia (without distal epiphysis), left patella, left fibula, right fibula (proximal not fused).

Sex: Undetermined (morphological features not fully developed at this age).

Age: 15–18 years, based on epiphysis–diaphysis fusion, tooth eruption, tempo-sphenoid suture, and pelvic bone development.

Dental analysis:

- Teeth in alveolar sockets: 13, 14, 16–18, 26–28, 31–36, 37–38, 41–44, 45–48
- Teeth lost postmortem: 11, 12, 15, 21, 22, 24, 25, 35, 38
- Abrasion: Degree 1 (enamel) on most teeth; degree 2 (dentin exposure) on 11, 21, 22, 26, 31, 32, 36, 41, 46
- Caries: Tooth 46 (buccal)
- Supragingival calculus (degree 1) on 13–16, 31–36, 41–42
- Chipping: Teeth 18 (lingual) and 48 (buco-mesial)
- Discoloration: Teeth 16–18, 26–28, 36, 47

Pathology:

- Cribra orbitalia: Left orbit (degree 4, active at death)
- Porotic hyperostosis: Both parietals (degree 3, active), frontal absent (0)

- Increased porosity on internal surface of left parietal
- Small thickening on left humerus above nutrient foramen (1.0 × 0.1 cm)

Metric analysis: All measurements and indices shown in Table 1ab of [23].

Stable isotope analysis: $\delta^{13}C$, $\delta^{15}N$, $\delta^{34}S$ indicate primarily terrestrial diet[23].

Comment: This juvenile, along with other children from Golokut-Vizić and outside the Danube Gorges, shows significant porotic hyperostosis. Lesions were mostly active at death, indicating insufficient healing. Terrestrial dietary patterns, particularly very low animal protein intake (e.g., Grave 2/2003), suggest nutritional imbalance. The combined evidence from cribra orbitalia and hyperostosis implies limited nutrient availability, likely affecting immunity and overall health.

### 2. Methodology on the Statistical Framework for Population Genomics tests

#### 2.1 Proximal Demographic Modeling of Newly Reported Individuals

To ensure the proper categorization of the ancestral origins of the 12 newly reported individuals, we designed a set of demographic models to statistically validate their ancestry using $f_4$-statistics. This model testing was performed with the qpAdm algorithm from ADMIXTOOLS v7.0, using the 'allsnps = YES' parameter and the '1240k' SNP array dataset, which was merged with publicly available data[24] (see **Methods**).

To achieve a genuinely neutral variant dataset for this analysis, we pruned 178,342 SNPs from the array that were associated with known phenotypic traits, subject to selective pressures, or in linkage disequilibrium with other selected variants (see **Methods**). The total SNP coverage for each of the newly reported individuals in the 1240k SNP panel was between 358,829 and 1,046,832, likely sufficient for individualised ancestry profiling (**Supplementary Table 1**).

We aimed to model the ancestry of the newly reported individuals using a highly proximal framework to detect subtle differences in their ancestral components. Before constructing the $f_4$-statistics tests and performing subsequent value matrix comparisons using qpAdm, we excluded any individuals whose genomic data were generated using methods different from those applied to our dataset. This was necessary to mitigate known allelic biases associated with different capture techniques[25]. By doing so, we sought to eliminate statistical distortions that could arise when calculating $f_4$-statistics due to generation-specific allelic biases.

For the demographic model design, we selected the following outgroup populations:

- **Mbuti.DG**: Chosen as the anchor out-population, representing basal out-of-Africa divergence[26].
- **Russia_Kostenki14_UP.SG**: Selected to capture basal West Eurasian ancestry (n=1)[27].
- **Russia_MA1_UP.SG** – Included as a distal pull for general Eurasian ancestry (n=1)[28].
- **Ukraine_Mesolithic.SG**: Representing Eastern Hunter-Gatherer (EHG)-like ancestry, serving as a more proximal reference for Western Hunter-Gatherer (WHG) ancestry (n=9)[29–31].
- **Turkey_Central_Musular_PPN.SG**: Selected to represent Early Farmer (EF) ancestry (n=2)[32].

During the model design and outgroup selection process, we considered additional populations but ultimately excluded them due to statistical limitations:

1. **Caucasus_HG.SG[33]**: Initially tested to assess a potential Caucasus-related genetic signal in Neolithic migrants in the Balkans[34]. However, models incorporating this population showed significant interactions between EHG and CHG in the f4 matrix without improving statistical discrimination.
2. **Latvia_Mesolithic.SG[35]**: Tested as a potential distal WHG reference. However, models including this outgroup exhibited strong interactions with WHG populations in the left populations, leading to their exclusion.
3. **Russia_UstIshim_IUP.DG[36]**: Considered as a pull for basal out-of-Africa ancestry. However, models incorporating this outgroup did not yield significant differences compared to those without it, suggesting it did not provide additional discriminatory power. Consequently, we excluded it from the final outgroup set.

Full details on these negative results are available upon request from the corresponding authors.

In order to ensure the testing of a true proximal model, we selected possible proximal proxy source populations:

- **Iron_Gates_Mesolithic.SG**: Composed by Serbia_IronGates_Mesolithic.SG and Romania_IronGates_Mesolithic.SG to act as a very proximal proxy source of Hunter Gatherer ancestry of the region (n=6)[31,37,38].

- **Balkan_Early_Neolithic.SG**: Composed by Serbia_LepenskiVir_EN.SG, Serbia_GradStarcevo_Starcevo_EN.SG, Romania_Negrilesti_StarcevoCris_EN.SG, Hungary_EN_Koros.SG, and Romania_Baciu_StarcevoCris_EN.SG to act as proximal proxy population of the EEFs societies' ancestry in the region (n=6)[29,31,38].

- **Turkey_Marmara_Barcin_N.SG**: Composed by 3 individuals from Barcın-Höyük with early Neolithic ancestry (n = 3)[39].

- **Luxembourg_Mesolithic.DG:** Composed by a WHG individual (n=1)[40].

Using the selected right (outgroup) and left (source/target) populations, we successfully identified a fitting demographic model for all 12 newly reported individuals. Initially, we attempted to model their ancestry using a simple one-way model (uni-source model). If this approach failed to explain the individual's genetic profile, we incrementally introduced additional left populations until a suitable model was found. Through this approach, we identified nine one-way models, two two-way models, and one three-way model for each of the newly reported individuals studied.

Six individuals (i705-i505, i706-i506, i707-i507, i716-i512, i722-i513, and i723-i515) could be modeled as deriving their entire ancestry from Iron_Gates_Mesolithic.SG, aligning with their placement in the Principal Component Analysis (PCA) (**Extended Data Fig. 3a**). This further supports their classification as belonging to the hunter-gatherer population that inhabited the Iron Gates region during the Mesolithic period.

Three additional individuals, i701-i501 and i708-i508, were best modeled with Balkan_Early_Neolithic.SG as their sole ancestral source, while i713-i509 was most closely associated with Turkey_Marmara_Barcin_N.SG. Notably, despite their proximity in PCA space to i701-i501 and i708-i508 (**Extended Data Fig. 3a**), a one-way model for i713-i509 using Balkan_Early_Neolithic.SG was statistically unsupported. A possible explanation for this discrepancy is that Turkey_Marmara_Barcin_N.SG exhibits lower WHG ancestry than Balkan_Early_Neolithic.SG, making it a better fit for i713-i509 in our fine-scale ancestry modeling approach. These results confirm that these three individuals belonged to Neolithic migrant populations originating from Anatolia, which expanded across western Eurasia, introducing agriculture during the Neolithic period.

Among the newly reported individuals, we identified three admixed cases. i702-i502 and i703-i503 were best modeled with a two-way ancestry model, deriving from a combination of Iron_Gates_Mesolithic.SG and Balkan_Early_Neolithic.SG. Individual i702-i502 exhibited an almost equal split between both ancestries, a finding consistent with its PCA positioning (**Extended Data Fig. 3a.**). This balanced admixture could reflect a recent mixture event between these two populations.

In contrast, i703-i503 displayed a much lower proportion of Balkan_Early_Neolithic.SG ancestry (15.6% ± 0.25). Despite this minor admixture, the individual was included in the WHG group for adaptation analyses, as the Neolithic ancestry component was unlikely to significantly influence the results.

Individual i704-i504 required a three-way ancestry model, incorporating Balkan_Early_Neolithic.SG, Iron_Gates_Mesolithic.SG, and Luxembourg_Mesolithic.DG. While this individual primarily derived its ancestry from WHG populations, a small proportion (9.3% ± 2.8) of Neolithic-related admixture suggests that contact between these groups occurred several generations before this individual's lifetime.

Additionally, the inclusion of Luxembourg_Mesolithic.DG as an extra WHG source was necessary to achieve a statistically valid model. However, this does not necessarily imply direct admixture between Loschbour-like populations and Balkan hunter-gatherers. Instead, the need for an additional WHG component may reflect reduced EHG ancestry in i704-i504, as indicated by its lower PC2 values in PCA space (**Extended Data Fig. 3a**). Similar to i703-i503, this individual was included in the WHG group for adaptation analyses despite its minor Neolithic admixture.

The results of the f4-statistics modeling enabled us to classify the newly reported individuals into two distinct groups for subsequent selection analysis (see **Methods**). Three individuals (i701-i501, i708-i508, and i713-i509) were classified as Early European Farmers (EEF). Eight individuals were classified as Western Hunter-Gatherers (WHG). One individual, i702-i502, was excluded due to its high level of admixture. This classification ensures robust population groupings for adaptation analyses while minimizing confounding effects from genetic mixture.

#### 2.2 Distal Demographic Modeling of Publicly Available Individuals

In this section, we describe the $f_4$-statistics based modelling of the whole dataset of 201 publicly available[24] and 12 newly sequenced individuals. We designed a model to determine the ancestral composition of each individual with respect to their ancestry proportions of Early Farmer (EF), Western Hunter-Gatherer (WHG), and Eastern Hunter-Gatherer (EHG) ancestries.

We performed $f_4$-statistics using the *qpAdm* module in ADMIXTOOLS v.6.0 with 'allsnps: YES'. For this set of analyses, we based the analyses on the neutral variant '1240k' SNP panel, mentioned above, also pruning the same 178,342 SNPs from the array that were associated with known phenotypic traits, subject to selective pressures, or in linkage disequilibrium with other selected variants.

For each target individual, we tested a one-way, a two-way, and a three-way model. This was a necessary prerequisite for filtering the dataset to a high-confidence set of samples for subsequent adaptive variant analysis (see below), to ensure a correct categorization of the ancestral origin for comparison.

For the demographic model design, we selected the following outgroup populations, selected to represent deep, unadmixed lineages that provide a stable background for estimating the proportions of three primary source ancestries:

- **Mbuti.DG**: Chosen as the anchor outgroup, representing a basal out-of-Africa divergence[26].
- **Russia_UstIshim_IUP.DG**: Selected to capture an ancient Eurasian lineage that predates the diversification of subsequent Eurasian populations[36].
- **Russia_Kostenki14_UP.SG**: Included to represent basal West Eurasian ancestry[27].
- **Russia_MA1_UP.SG**: Selected as a distal pull for ancient North Eurasian (ANE) ancestry, which helps to differentiate between WHG and EHG components[28].
- **Italy_Epigravettian.AG**: Incorporated to represent a southern European hunter-gatherer lineage, providing further resolution in the outgroup set[41].
- **Turkey_Central_Musular_PPN.SG**: Selected as a distal outgroup related to, but predating, the early farmers of Europe[32].

Using this stable set of right outgroup populations, we modeled each target individual as a mixture of three possible proximal source populations in a one-way, two-way, and three-way *qpAdm* model. These sources were chosen to represent the proxy primary ancestries known to have contributed to Mesolithic and post-Mesolithic European populations:

- **Russia_YuzhniyOleniyOstrov_Mesolithic.AG**: Selected as a proxy source for Eastern Hunter-Gatherer (EHG) ancestry[35,41,42].
- **Turkey_Marmara_Barcin_N.AG:** Selected as a proxy source for Early European Farmer (EF) ancestry[42,43].
- **W_Mesolithic.AG:** A composite population selected as a proxy source for Western Hunter-Gatherer (WHG) ancestry[44–46].

This initial qpAdm-based classification was applied to all 201 public individuals and the 12 newly reported ones. After discarding individuals without a fitting model (by means of p-value, negative ancestry proportion, or high standard errors), and also discarding individuals with more than one fitting model, this process resulted in a set of 189 public and 12 newly sequenced individuals with successfully attributed ancestry profiles. The results of this distant ancestry proportions modelling can be found in **Supplementary Table 1**.

Through a sequential process of quality control for the dataset used in the adaptive variants analysis (see methods), we derived a final, high-confidence dataset of 152 individuals (141 public and 11 newly sequenced) for subsequent analyses (**Supplementary Table 1**). This set comprises 48 individuals of EF ancestry, 11 of HG ancestry, and 82 with admixed ancestry, achieving robust population characterizations for further downstream adaptation analyses.

#### 3. Extended Results and Discussion

##### 3.1 Exploring the Feasibility of SNP calling

Accurate SNP discovery (i.e. finding all SNPs across the whole genome of an individual, as opposed to only those within predefined locations) is not trivial for aDNA samples. There are no clearly established pipelines to do so, and most studies used algorithms designed for and tested on high-coverage modern DNA datasets. Although it could have been possible to just use some available pipelines (e.g. the Atlas pipeline used in Marchi et. al.[38], or AriaDNA[47]), we considered that our research aim required a custom pipeline ensuring high SNP calling accuracy in our samples. Note that we were not only interested in the overall genetic structure of the analyzed populations, but we also wanted to characterize specific SNPs. Thus, we needed to be highly confident about all called variants, and not only a fraction of them. In addition, the highly heterogeneous nature of our datasets (see **Supplementary Table 1**) made the task of SNP calling and filtering particularly complex, as it was unclear whether we could identify SNPs using the same algorithms/parameters for all samples.

To ensure accurate SNP calling in all samples, we implemented a custom pipeline that tailored SNP calling algorithms and filtering parameters towards each sample, by using sample-specific simulations (see **Methods**). This was inspired by our previous work on structural variant calling[48]. For this, we first generated simulated reads matching the technical properties of the real samples, including their read length, post-mortem damage (PMD) profile, coverage, and contaminant DNA proportions. We generated three simulations for a given sample, each containing a realistic set of SNPs based on the ancient individuals KK1, LEPE48 and VLASA7[38]. To validate the simulations, we empirically verified that their coverage, read length and PMD profile matched those of the real samples (see **Methods**, **Extended Data Fig. 4b**,**5**,**6**).

Next, we evaluated the effect of various parameters on the SNP calling accuracy, including i) the minimum average coverage that a sample should have to be considered (cov_s), ii) the minimum number of reads that a position should have in all considered samples (depending on cov_s) to be considered (cov_p), iii) the calling algorithm (including GATK HaplotypeCaller, freebayes, bcftools, freebayes in pooled mode and AriaDNA), iv) the set of samples considered (sample_set, all datasets or various downsampled datasets) and v) various SNP filtering settings. Note that cov_s and cov_p were used to constrain the calling procedure towards genomic positions with sufficient coverage, as coverage is likely the main predictor of calling performance (**Extended Data Fig. 7**). We also tested various downsampled subsets of the data (sample_set parameter). We performed this step because considering all samples with certain cov_p thresholds may lead to insufficient numbers of positions with enough coverage. This approach allowed us to evaluate whether we could perform reliable SNP calling at least on a subset of the data. To define such sample sets, we first distributed all samples into seven chunks, each containing samples covering the coverage and genetic range in all populations of interest (EF, HG, and the three admixed ones) (**Extended Data Fig. 9**). Then, we defined various sample sets, each including various combinations of these chunks, resulting in sets with a range of sample sizes (see **Extended Data Fig. 8a**). In total, for a given combination of cov_p, cov_s, and sample_set, we benchmarked the performance of 695,592 combinations of algorithms and filters.

For each of these combinations, we inferred the calling performance from the F-value (harmonic mean between precision and recall), calculated from the overlap between called and expected (inserted during read simulation) SNPs (see **Methods**). Also, to ensure that our cov_p settings are not too conservative we calculated the percent of the genome covered by each combination. Our objective was to find a combination of parameters (cov_s, cov_p, sample_set) yielding sufficient genomic coverage (at least 20%), acceptable calling performance (at least 0.65), while considering a sufficient number of samples to run selection analyses. We could not find any parameters matching these criteria (**Extended Data Fig. 8**). There was one combination that gave almost 20% coverage and >0.7 F-value (cov_s=3, cov_p=2), but it was on a sample_set that only included a small fraction of our dataset (HG, admixed individuals with >0.45 HG ancestry (admHG), and one chunk of downsampled EF samples (EF_down1)).

Based on this exploration, we conclude that we have insufficient coverage to perform accurate SNP calling on our data. Still, our sample-tailored simulation-based approach to benchmark the performance of various parameters may serve as a useful starting point for future efforts to call variants in aDNA data.

##### 3.2 SNP-based Selection Scans

To infer differential selective pressures between ancestral HG/EF populations from the imputed SNPs, we used two complementary approaches. First, we used PCAdapt, as in previous studies of adaptation during the Mesolithic-Neolithic transition[49] to detect SNPs driving the separation between the HG/EF populations along the major PCA axis (**Fig. 2a,b**). We defined as 'significant' the 300 SNPs with the lowest p values, all of which had a $p < 4.42 \cdot 10^{-77}$ (**Fig. 2c,d**). To focus on the most reliable, non-redundant signals, likely related to selective sweeps, we selected 8 'PCAdapt hits', each representing a genomic cluster with >2 significant SNPs within <100 kb. (**Fig. 3, Supplementary Table 2**). Second, we performed a haplotype-based XP-EHH scan to identify more recent selective sweeps and their direction in each of the EF and HG populations, as in previous similar studies[50]. This approach identifies selection by finding positions with longer Extended Haplotype Homozygosity (EHH) in a 'target' population vs a 'background' one, with positive values reflecting selection in EF, while negative values reflect selection in HG in our case (**Fig. 2 e**). We identified 20 non-redundant XP-EHH hits, each representing a cluster with >2 significant positions (Bonferroni-corrected p<0.05) within <100 kb (**Fig. 2f,g, 3; Supplementary Table 2**), of which 12 indicate selection in EF and 8 indicate selection in HG, supporting lineage-specific sweeps in both ancestral populations.

To infer signs of adaptive admixture in each of the three regional admixed populations (nIre, nGer, nPor), we again used two complementary approaches. First, as done in similar previous studies[42,51], we calculated Fadm statistics and associated p values, which measure deviations between observed allele frequencies and those expected under an admixture model parameterized from ancestry proportions inferred by qpAdm (**Fig. 2h**). To infer selection direction we calculated EF Local Ancestry Deviation (EF LAD) for significant SNPs (Bonferroni-corrected p<0.05, **Fig. 2i**), as done in [51,52], a haplotype-based z-score quantifying how much the proportion of EF Local Ancestry Tracts (LATs) at a given SNP deviates from the genomic average (see **Methods**, **Extended Data Fig. 10**). Thus, SNPs with high EF LAD (>=2, i.e. two standard deviations away from the mean) were considered to reflect selection favouring EF alleles, and *vice versa* (EF LAD <= -2) for HG alleles (**Fig. 2j**). We finally kept 27 non-redundant 'Fadm hits' (12 from nPor, 9 from nIre, 6 from nGer), each representing a cluster with >2 significant SNPs with equivalent directionality within <100 kb (**Fig. 3, Supplementary Table 2**). All but one of these hits had no clear directionality (EF LAD between -2 and 2), suggesting that most allele-frequency deviations cannot be straightforwardly attributed to preferential increases of either EF- or HG-derived local ancestry.

Second, to find signs of adaptive admixture specifically favouring EF or HG alleles, we adapted the recently proposed haplotype-based iDAT (integrated Decay in Ancestry Tracts) method, to our dataset. iDAT measures whether EF LATs overlapping a given position are, on average, longer than the corresponding HG LATs, as expected under a recent selective sweep favouring local EF ancestry, or *vice versa* (**Fig. 2k, Extended Data Fig. 11**). From 'significant' positions having iDAT ≥ 2 or iDAT ≤ -2 for EF- or HG-favouring selection, respectively (**Fig. 2j**), we selected 30 'iDAT hits' (10 from nPor, 10 from nIre, 10 from nGer) corresponding to the top 10 hits per population and each representing a cluster with >2 significant hits with equivalent directionality within <100 kb (**Fig. 3, Supplementary Table 2**). Of these, 19 favour EF ancestry and 11 favour HG ancestry, again indicating contributions from both ancestral backgrounds to post-admixture adaptation.

The above selection scans defined a total of 85 hits. To validate them and focus on the most relevant candidates, we used two strategies. First, since different methods may capture equivalent selective processes (e.g. a given selective sweep in EF captured by PCAdapt and XP-EHH, or another in nGer captured by Fadm and iDAT), we tried to prioritize SNPs shared among the top hits of different scans (PCAdapt, XP-EHH, Fadm nIre, Fadm nPor, Fadm nGer, iDAT nIre, iDAT nPor or iDAT nGer). While each scan yielded non-overlapping top hits, we found 20 reaching significance in at least one additional scan. Closer inspection revealed that 9 of these 20 overlaps were consistent with the same ancestry-favouring iDAT pattern across multiple admixed populations, potentially reflecting selective processes already present in their shared ancestral population (**Extended Data Fig. 12, 13**). In contrast, the remaining 11 overlaps reflected more complex relationships, including i) selection in one of the ancestral HG/EF populations, and consistent favouring of the alleles of the same population in the admixed ones (1), ii) discordant ancestry directions across admixed populations (2), iii) combinations of ancestry-specific and ancestry-independent signals (2), or iv) apparent opposite direction of selection in ancestral and admixed populations (6). These results indicate that overlap across methods does not often reflect a single shared sweep. Rather, the limited agreement between scans suggests that the different approaches capture complementary selection impacts acting at different evolutionary timescales. Similarly, the low overlap across admixed populations points to substantial regional specificity, consistent with region-specific adaptive processes during the Mesolithic–Neolithic transition, which we could characterize thanks to our geographically-structured adaptive admixture scan.

Second, we compared our top hits to loci reported in previous similar selection scans across ancient Eurasian individuals[49–51,53] (see **Methods, Extended Data Fig. 14,15, Supplementary Table 3**). Out of six 1 Mb regions including variants reported in more than one previous study or by more than one selection detection method, only two SNPs overlapped with our hits, including *FADS1* and *HLA* genes (**Extended Data Fig. 14**). However, we could not find equivalent evidence for the remaining regions, overlapping *JAK1*, *NFKBIZ*, *SLC24A5* and *BDNF* (**Extended Data Fig. 15**). This may result from differences in methodologies, populations under study or significance thresholds, such as the ones used for PCadapt scans ($10^{-30}$-$10^{-40}$ in [49], versus our $10^{-77}$) or XP-EHH analyses (XP-EHH threshold of 2.0 in [50], versus our 5.78). More broadly, previous studies also exhibit very little overlap among them (**Extended Data Fig. 14,15**), suggesting that overlap with prior studies provides a weak basis for prioritization. Hence, we retained all selected 85 top-hits for downstream analyses, treating them as largely independent signals of past selective sweeps in HG, EF and admixed populations during the Mesolithic-Neolithic transition.

##### 3.3 Transposable Element Analysis

We developed an aDNA-specific analytical framework to specifically assess the contribution of TEs (AncienTE, see **Methods, Fig. 5a, Extended Data Fig. 18**,**19,20**). This analysis identified 47 TE families with significantly different abundances between HG and EF populations after Bonferroni correction ($p<0.05$) (22 enriched in HG and 25 in EF; **Fig. 5b-c.**, **Extended Data Fig. 21**). To test whether TE abundance variation was associated with ancestry independently of sequencing depth or sample age, we fitted a linear model of the form y ~ ancestry_prop + Coverage + Date_BCE, where *y* corresponds to TE family abundance. Ancestry proportion remained a significant predictor for the TE families identified above (**Extended Data Fig. 22**). These families comprised 18 LTR/ERV1, 12 LINE/L1, 9 LTR/ERVK, 6 SINE/Alu, 1 SVA, and 1 DNA/TcMar-Tigger. Notably, most belong to evolutionarily young retrotransposon groups with documented recent or ongoing activity in modern humans, such as LINE/L1 and the non-autonomous Alu and SVA elements mobilized by L1 machinery[54], as well as ERVK/HERV-K (HML-2), the youngest endogenous retrovirus lineage[55]. The direction of enrichment differed across superfamilies. Among HG-enriched TE families, the most represented were LTR/ERVK (9 families), and SINE/Alu (6 families), followed by LTR/ERV1 (4 families). This is consistent with evidence that SINE/Alus are known for playing a major role in the evolution of primate-specific enhancers and gene regulatory networks[56,57]. By contrast, EF-enriched TE families were predominantly LTR/ERV1 (13 families), and LINE/L1 (11 families), revealing a marked shift in the superfamily composition of differentially abundant elements between populations.

The predominance of evolutionarily young retrotransposon families among the differentially abundant elements suggests a potential link with genomic plasticity. LINE/L1, SINE/Alu and SVA elements retain mobilization capacity in modern humans, and their activity has been associated with insertional mutagenesis, structural variation, and altered gene regulation[58–60]. Similarly, ERVK (HML-2), preserves coding potential and transcriptional competence at some loci in modern humans[55]. Given the highly degraded nature of aDNA, our data do not allow precise genomic localization of these elements or identification of insertion polymorphisms, so the specific functional consequences remain unresolved. Although it is hard to link differences in germline TE abundance with phenotypic consequences, increased representation of mobilizable elements could result in increased genomic instability in somatic tissues, driving changes in epigenetic regulation or resulting in higher susceptibility to somatic retrotransposition[61]. Finally, it is plausible that differences in TE content between populations may influence the mutational landscape over evolutionary timescales, providing a mechanistic framework through which ancestry-associated TE variation could contribute to long-term phenotypic differences.

We next investigated TE dynamics in admixed individuals by constructing an admixture-based expectation model using inferred ancestry proportions (see **Methods**) to identify TE families deviating from neutral ancestry-driven expectations. To assess local admixture dynamics, we separately analyzed admixed populations of nGer, nIre and nPor **(Fig. 5d-f**), which yielded 78, 171, and 2, TE families with significant deviations (Bonferroni-corrected $p<0.05$), respectively (**Fig. 5g-i, Extended Data Fig. 23,24**). In the nGer population, most enriched TE families (87%) corresponded to endogenous retroviruses, predominantly LTR/ERV1 (45/78), followed by LTR/ERVL (12/78) and LTR/ERVK (11/78). A similar pattern was observed in nIre, where 73% of enriched TE families were retroviral, including LTR/ERV1 (84/171), LTR/ERVK (28/171) and LTR/ERVL 16/171). Both TE families identified in the nPor population belonged to the LTR/ERVK superfamily (**Supplementary Table 5**).

Together, these results show that TE abundance profiles capture ancestry-associated genomic variation in ancient individuals and reveal structured shifts in evolutionarily young TE superfamilies. Despite the limitations imposed by short, degraded aDNA reads, TE abundance patterns vary across HG/EF populations, track admixture gradients and uncover regional deviations from neutral ancestry-driven expectations. The enrichment of distinct retrotransposon lineages across populations suggests that recently active elements followed different evolutionary trajectories in prehistoric Eurasia. Although the precise genomic locations and functional consequences of these shifts remain unresolved, our findings demonstrate that repeatome dynamics can be interrogated in ancient genomes and provide complementary insights to SNP and CNV-based analyses.
