## Extended figures 1-12 for "Ancient whole genomes reveal regional selection during adaptation to Neolithic lifestyle in Western Eurasia"

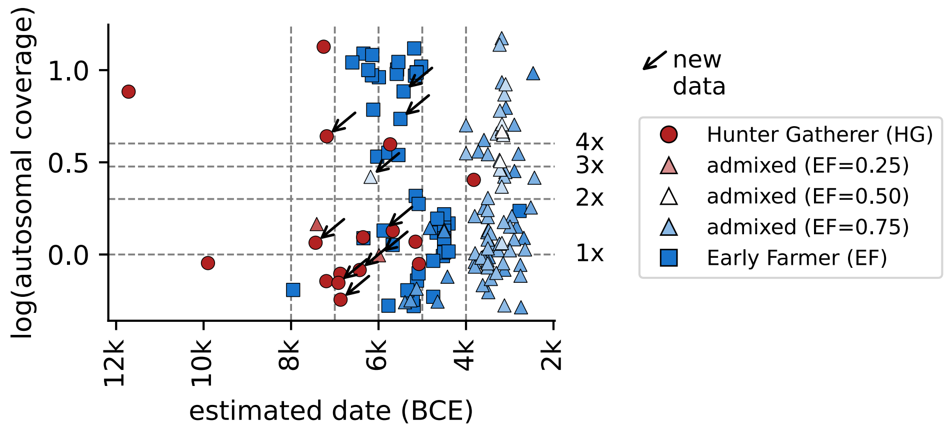

log(autosomal coverage)
1.0
0.5
0.0
4x
3x
2x
1x
12k
10k
8k
6k
4k
2k
estimated date (BCE)
new
data
Hunter Gatherer (HG)
admixed (EF=0.25)
admixed (EF=0.50)
admixed (EF=0.75)
Early Farmer (EF)

**Extended Data Fig. 1. Coverage of analyzed individuals.** Log-scaled average depth for all individuals analyzed in this study, plotted against the estimated chronological date. The color represents the EF proportion, as in **Fig. 1**. Note that we only included genomes with coverage < 0.5x.

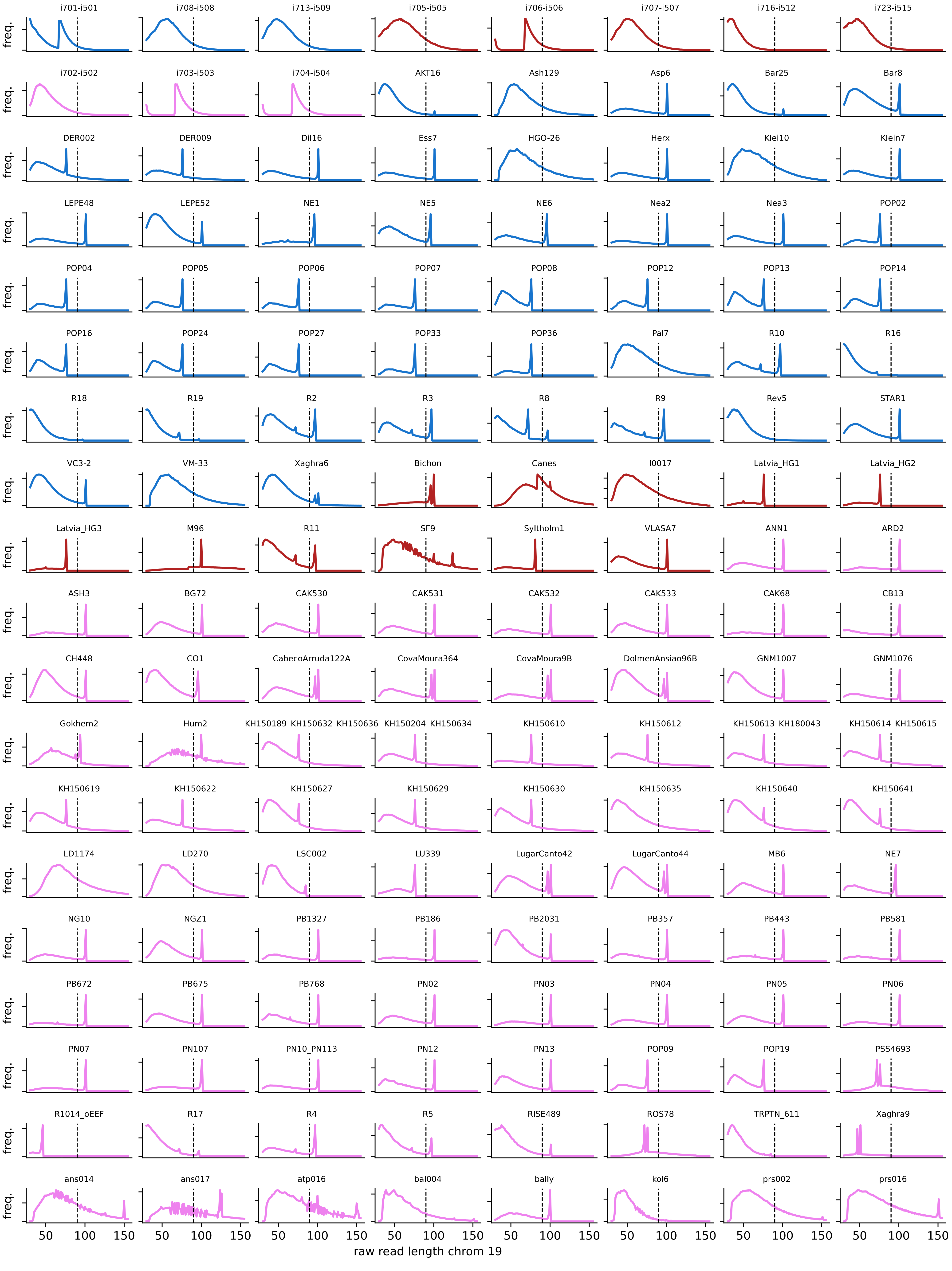
i701-i501
i708-i508
i713-i509
i705-i505
i706-i506
i707-i507
i716-i512
i723-i515
i702-i502
i703-i503
i704-i504
AKT16
Ash129
Asp6
Bar25
Bar8
DER002
DER009
Dil16
Ess7
HGO-26
Herx
Klei10
Klein7
LEPE48
LEPE52
NE1
NE5
NE6
Nea2
Nea3
POP02
POP04
POP05
POP06
POP07
POP08
POP12
POP13
POP14
POP16
POP24
POP27
POP33
POP36
Pal7
R10
R16
R18
R19
R2
R3
R8
R9
Rev5
STAR1
VC3-2
VM-33
Xaghra6
Bichon
Canes
I0017
Latvia_HG1
Latvia_HG2
Latvia_HG3
M96
R11
SF9
Syltholm1
VLASA7
ANN1
ARD2
ASH3
BG72
CAK530
CAK531
CAK532
CAK533
CAK68
CB13
CH448
CO1
CabecoArruda122A
CovaMoura364
CovaMoura9B
DolmenAnsiao96B
GNM1007
GNM1076
Gokhem2
Hum2
KH150189_KH150632_KH150636
KH150204_KH150634
KH150610
KH150612
KH150613_KH180043
KH150614_KH150615
KH150619
KH150622
KH150627
KH150629
KH150630
KH150635
KH150640
KH150641
LD1174
LD270
LSC002
LU339
LugarCanto42
LugarCanto44
MB6
NE7
NG10
NGZ1
PB1327
PB186
PB2031
PB357
PB443
PB581
PB672
PB675
PB768
PN02
PN03
PN04
PN05
PN06
PN07
PN107
PN10_PN113
PN12
PN13
POP09
POP19
PSS4693
R1014_oEEF
R17
R4
R5
RISE489
ROS78
TRPTN_611
Xaghra9
ans014
ans017
atp016
bal004
bally
kol6
prs002
prs016
freq.
50
100
150
raw read length chrom 19

**Extended Data Fig. 2. Distribution of raw read lengths.** Only for chromosome 19-mapped reads after read de-duplication. The line at 90 bp reflects the threshold used, discarding longer reads to avoid modern human contamination.

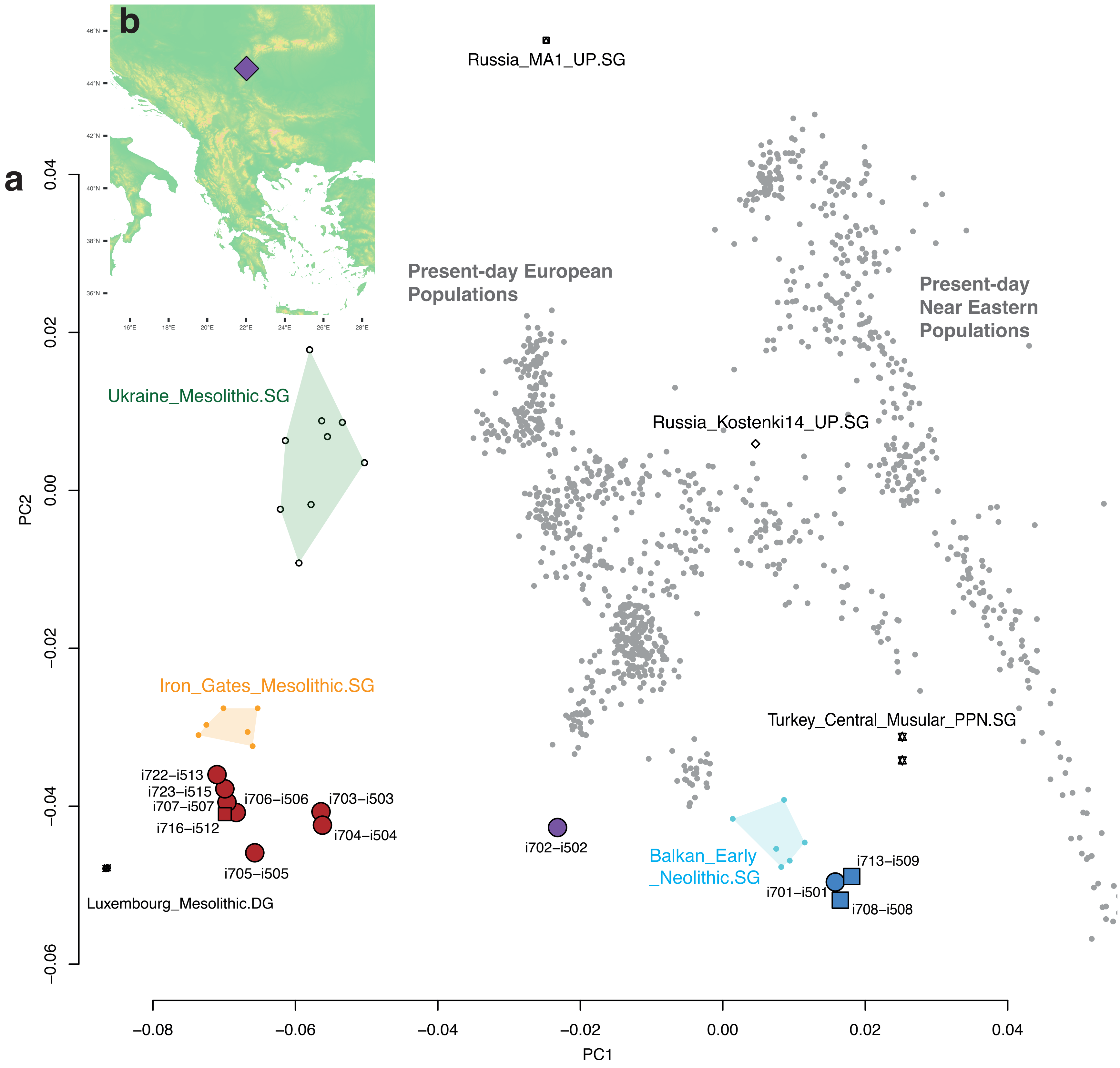

a
b
Russia_MA1_UP.SG
Present-day European Populations
Present-day Near Eastern Populations
Ukraine_Mesolithic.SG
Russia_Kostenki14_UP.SG
Iron_Gates_Mesolithic.SG
Turkey_Central_Musular_PPN.SG
i722–i513
i723–i515
i707–i507
i716–i512
i706–i506
i703–i503
i704–i504
i705–i505
i702–i502
Balkan_Early_Neolithic.SG
i713–i509
i701–i501
i708–i508
Luxembourg_Mesolithic.DG
PC1
PC2
0.04
0.02
0.00
−0.02
−0.04
−0.06
−0.08
−0.06
−0.04
−0.02
0.00
0.02
0.04

**Extended Data Fig. 3. Summary of the archaeological context and ancestral background of the 12 newly reported ancient individuals**. **a.** Principal Component Analysis plot of present-day Western Eurasian genetic variability for the Human Origins SNP panel with projected relevant ancient individuals. Note that there are 12 individuals sequenced here (red, blue, purple dots), not 11 as indicated in **Fig. 1a**, since we discarded one low coverage HG due to the inclusion criteria of the final dataset. **b.** Geographical location of Lepenski Vir archaeological site.

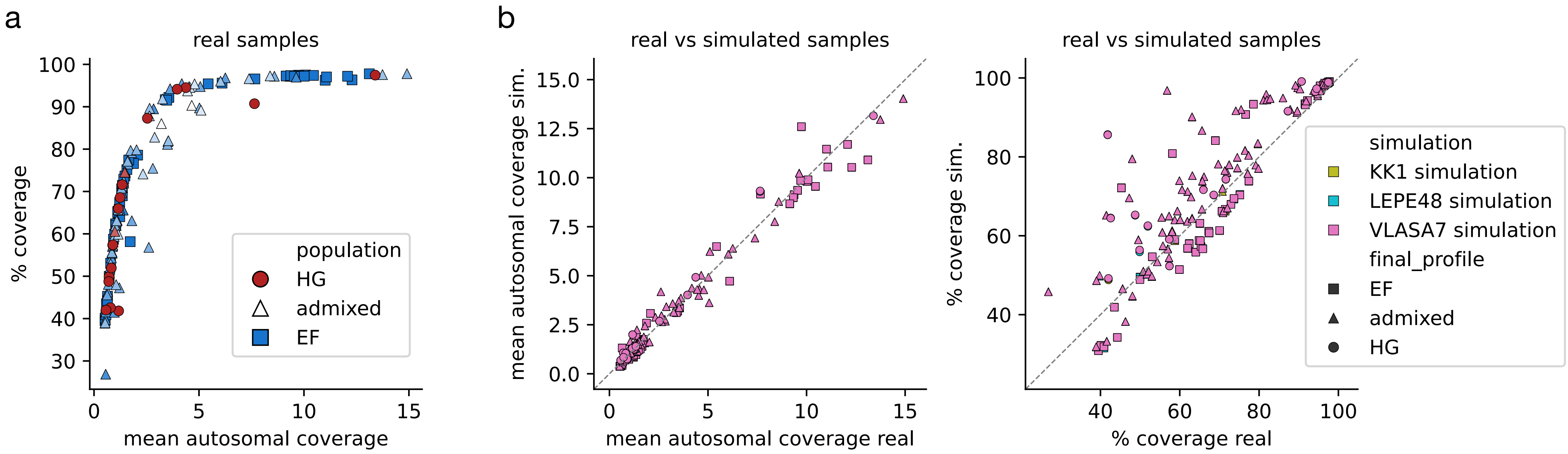

a
real samples
% coverage
mean autosomal coverage
population
HG
admixed
EF
b
real vs simulated samples
mean autosomal coverage sim.
mean autosomal coverage real
real vs simulated samples
% coverage sim.
% coverage real
simulation
KK1 simulation
LEPE48 simulation
VLASA7 simulation
final_profile
EF
admixed
HG

**Extended Data Fig. 4. Coverage validations for simulations. a.** Correlation between the mean autosomal coverage depth vs the percent of the genome covered by at least one read, in real samples. The colors are equivalent to those in **Fig. 1b**. **b.** Correlation between the coverage statistics (mean autosomal depth on the left, percent genome covered on the right) of real samples vs those in the simulated data.

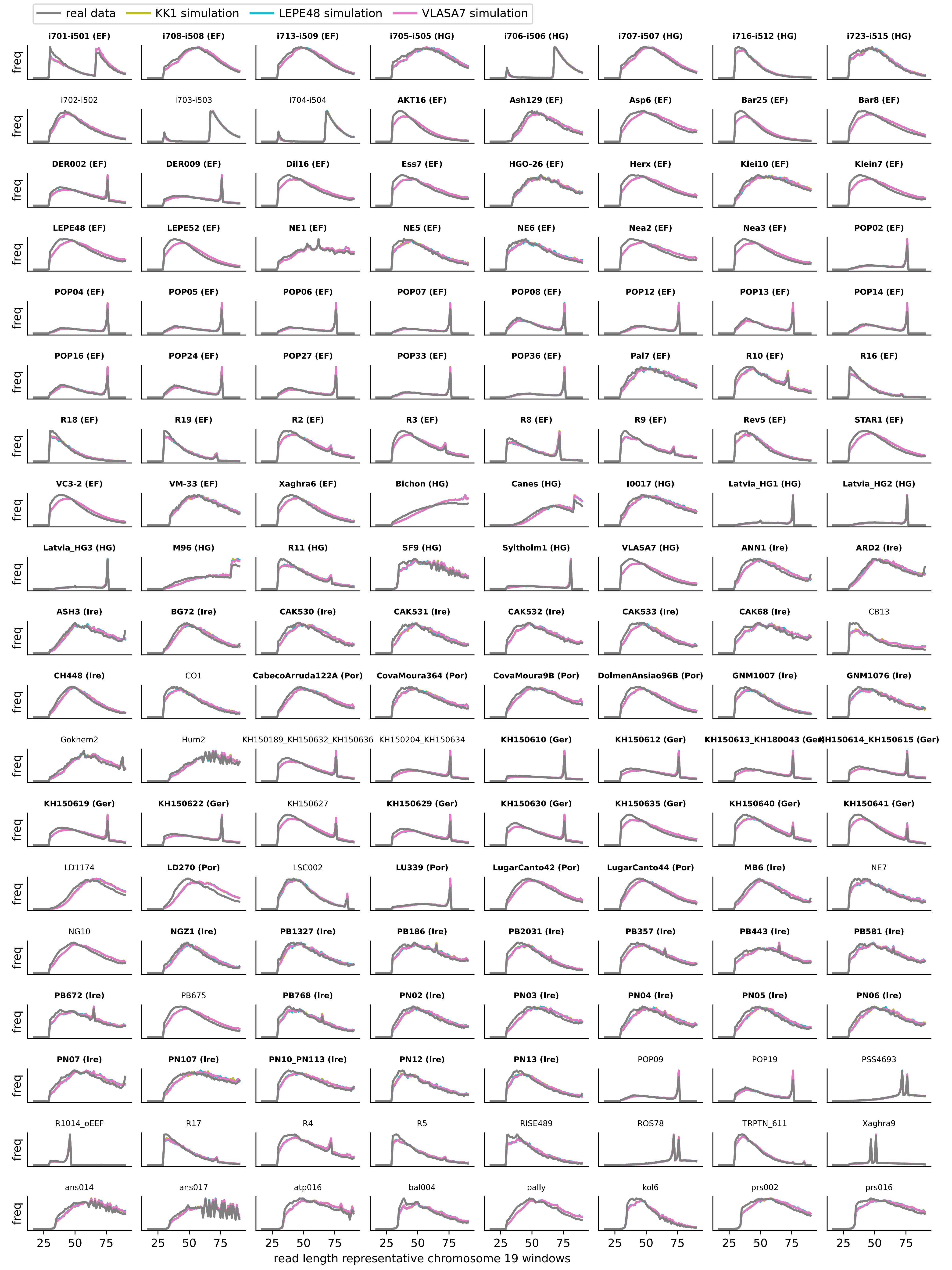

real data
KK1 simulation
LEPE48 simulation
VLASA7 simulation
i701-i501 (EF)
i708-i508 (EF)
i713-i509 (EF)
i705-i505 (HG)
i706-i506 (HG)
i707-i507 (HG)
i716-i512 (HG)
i723-i515 (HG)
i702-i502
i703-i503
i704-i504
AKT16 (EF)
Ash129 (EF)
Asp6 (EF)
Bar25 (EF)
Bar8 (EF)
DER002 (EF)
DER009 (EF)
Dil16 (EF)
Ess7 (EF)
HGO-26 (EF)
Herx (EF)
Klei10 (EF)
Klein7 (EF)
LEPE48 (EF)
LEPE52 (EF)
NE1 (EF)
NE5 (EF)
NE6 (EF)
Nea2 (EF)
Nea3 (EF)
POP02 (EF)
POP04 (EF)
POP05 (EF)
POP06 (EF)
POP07 (EF)
POP08 (EF)
POP12 (EF)
POP13 (EF)
POP14 (EF)
POP16 (EF)
POP24 (EF)
POP27 (EF)
POP33 (EF)
POP36 (EF)
Pal7 (EF)
R10 (EF)
R16 (EF)
R18 (EF)
R19 (EF)
R2 (EF)
R3 (EF)
R8 (EF)
R9 (EF)
Rev5 (EF)
STAR1 (EF)
VC3-2 (EF)
VM-33 (EF)
Xaghra6 (EF)
Bichon (HG)
Canes (HG)
I0017 (HG)
Latvia_HG1 (HG)
Latvia_HG2 (HG)
Latvia_HG3 (HG)
M96 (HG)
R11 (HG)
SF9 (HG)
Syltholm1 (HG)
VLASA7 (HG)
ANN1 (Ire)
ARD2 (Ire)
ASH3 (Ire)
BG72 (Ire)
CAK530 (Ire)
CAK531 (Ire)
CAK532 (Ire)
CAK533 (Ire)
CAK68 (Ire)
CB13
CH448 (Ire)
CO1
CabecoArruda122A (Por)
CovaMoura364 (Por)
CovaMoura9B (Por)
DolmenAnsiao96B (Por)
GNM1007 (Ire)
GNM1076 (Ire)
Gokhem2
Hum2
KH150189_KH150632_KH150636
KH150204_KH150634
KH150610 (Ger)
KH150612 (Ger)
KH150613_KH180043 (Ger
KH150614_KH150615 (Ger)
KH150619 (Ger)
KH150622 (Ger)
KH150627
KH150629 (Ger)
KH150630 (Ger)
KH150635 (Ger)
KH150640 (Ger)
KH150641 (Ger)
LD1174
LD270 (Por)
LSC002
LU339 (Por)
LugarCanto42 (Por)
LugarCanto44 (Por)
MB6 (Ire)
NE7
NG10
NGZ1 (Ire)
PB1327 (Ire)
PB186 (Ire)
PB2031 (Ire)
PB357 (Ire)
PB443 (Ire)
PB581 (Ire)
PB672 (Ire)
PB675
PB768 (Ire)
PN02 (Ire)
PN03 (Ire)
PN04 (Ire)
PN05 (Ire)
PN06 (Ire)
PN07 (Ire)
PN107 (Ire)
PN10_PN113 (Ire)
PN12 (Ire)
PN13 (Ire)
POP09
POP19
PSS4693
R1014_oEEF
R17
R4
R5
RISE489
ROS78
TRPTN_611
Xaghra9
ans014
ans017
atp016
bal004
bally
kol6
prs002
prs016
freq
25 50 75
read length representative chromosome 19 windows

**Extended Data Fig. 5. Read length distribution of real and simulated datasets.** Only for chromosome 19-mapped reads after read de-duplication. The colors represent the type of sample, either real (gray) or simulated (other colors).

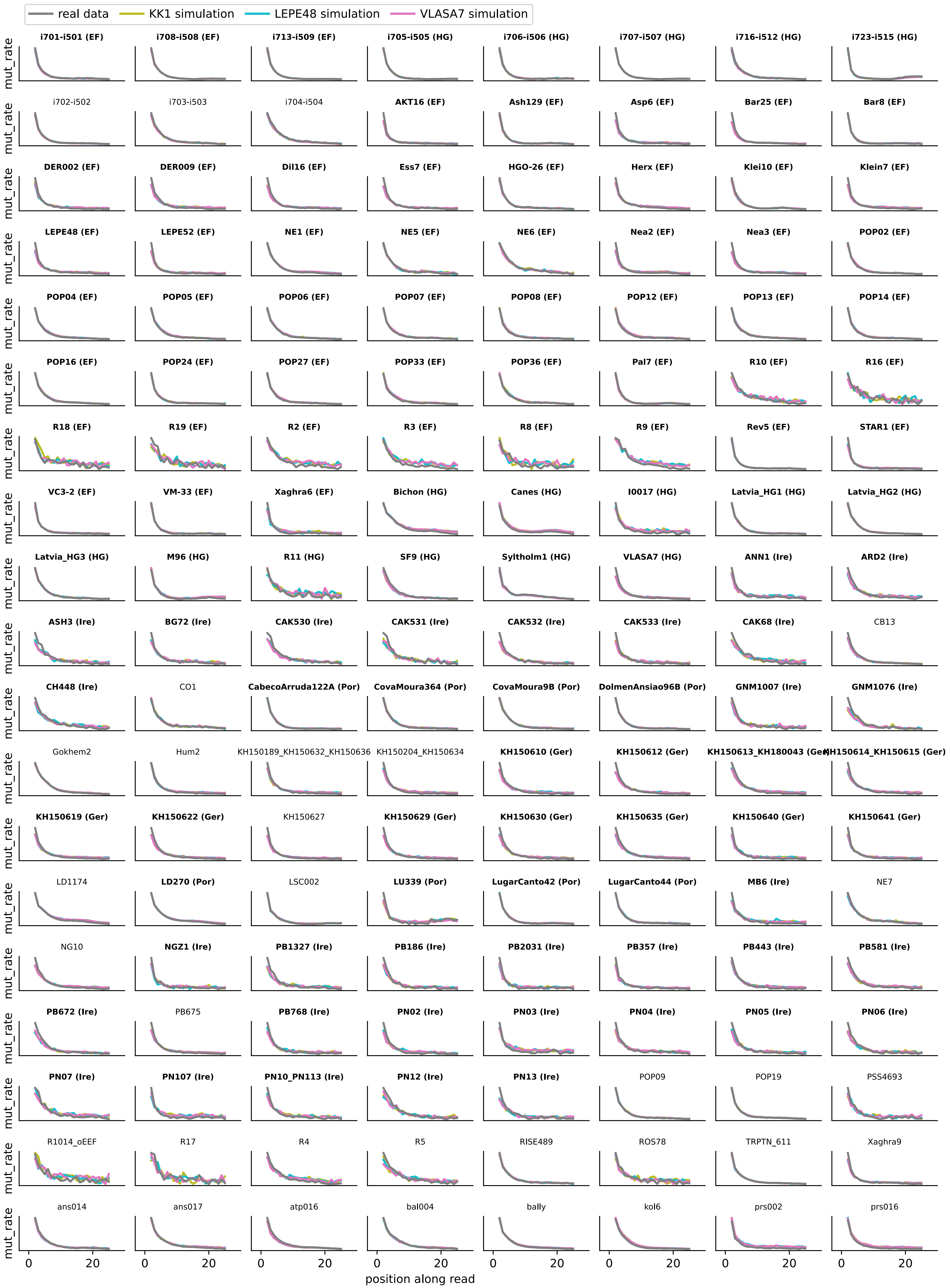
real data
KK1 simulation
LEPE48 simulation
VLASA7 simulation
i701-i501 (EF)
i708-i508 (EF)
i713-i509 (EF)
i705-i505 (HG)
i706-i506 (HG)
i707-i507 (HG)
i716-i512 (HG)
i723-i515 (HG)
i702-i502
i703-i503
i704-i504
AKT16 (EF)
Ash129 (EF)
Asp6 (EF)
Bar25 (EF)
Bar8 (EF)
DER002 (EF)
DER009 (EF)
Dil16 (EF)
Ess7 (EF)
HGO-26 (EF)
Herx (EF)
Klei10 (EF)
Klein7 (EF)
LEPE48 (EF)
LEPE52 (EF)
NE1 (EF)
NE5 (EF)
NE6 (EF)
Nea2 (EF)
Nea3 (EF)
POP02 (EF)
POP04 (EF)
POP05 (EF)
POP06 (EF)
POP07 (EF)
POP08 (EF)
POP12 (EF)
POP13 (EF)
POP14 (EF)
POP16 (EF)
POP24 (EF)
POP27 (EF)
POP33 (EF)
POP36 (EF)
Pal7 (EF)
R10 (EF)
R16 (EF)
R18 (EF)
R19 (EF)
R2 (EF)
R3 (EF)
R8 (EF)
R9 (EF)
Rev5 (EF)
STAR1 (EF)
VC3-2 (EF)
VM-33 (EF)
Xaghra6 (EF)
Bichon (HG)
Canes (HG)
I0017 (HG)
Latvia_HG1 (HG)
Latvia_HG2 (HG)
Latvia_HG3 (HG)
M96 (HG)
R11 (HG)
SF9 (HG)
Syltholm1 (HG)
VLASA7 (HG)
ANN1 (Ire)
ARD2 (Ire)
ASH3 (Ire)
BG72 (Ire)
CAK530 (Ire)
CAK531 (Ire)
CAK532 (Ire)
CAK533 (Ire)
CAK68 (Ire)
CB13
CH448 (Ire)
CO1
CabecoArruda122A (Por)
CovaMoura364 (Por)
CovaMoura9B (Por)
DolmenAnsiao96B (Por)
GNM1007 (Ire)
GNM1076 (Ire)
Gokhem2
Hum2
KH150189_KH150632_KH150636
KH150204_KH150634
KH150610 (Ger)
KH150612 (Ger)
KH150613_KH180043 (Ger)
KH150614_KH150615 (Ger)
KH150619 (Ger)
KH150622 (Ger)
KH150627
KH150629 (Ger)
KH150630 (Ger)
KH150635 (Ger)
KH150640 (Ger)
KH150641 (Ger)
LD1174
LD270 (Por)
LSC002
LU339 (Por)
LugarCanto42 (Por)
LugarCanto44 (Por)
MB6 (Ire)
NE7
NG10
NGZ1 (Ire)
PB1327 (Ire)
PB186 (Ire)
PB2031 (Ire)
PB357 (Ire)
PB443 (Ire)
PB581 (Ire)
PB672 (Ire)
PB675
PB768 (Ire)
PN02 (Ire)
PN03 (Ire)
PN04 (Ire)
PN05 (Ire)
PN06 (Ire)
PN07 (Ire)
PN107 (Ire)
PN10_PN113 (Ire)
PN12 (Ire)
PN13 (Ire)
POP09
POP19
PSS4693
R1014_oEEF
R17
R4
R5
RISE489
ROS78
TRPTN_611
Xaghra9
ans014
ans017
atp016
bal004
bally
kol6
prs002
prs016
mut_rate
0
20
position along read

**Extended Data Fig. 6. PMD profile of real and simulated datasets.** 5′ C>T mutation rate at different read positions, for real (gray) and simulated (other colors) datasets.

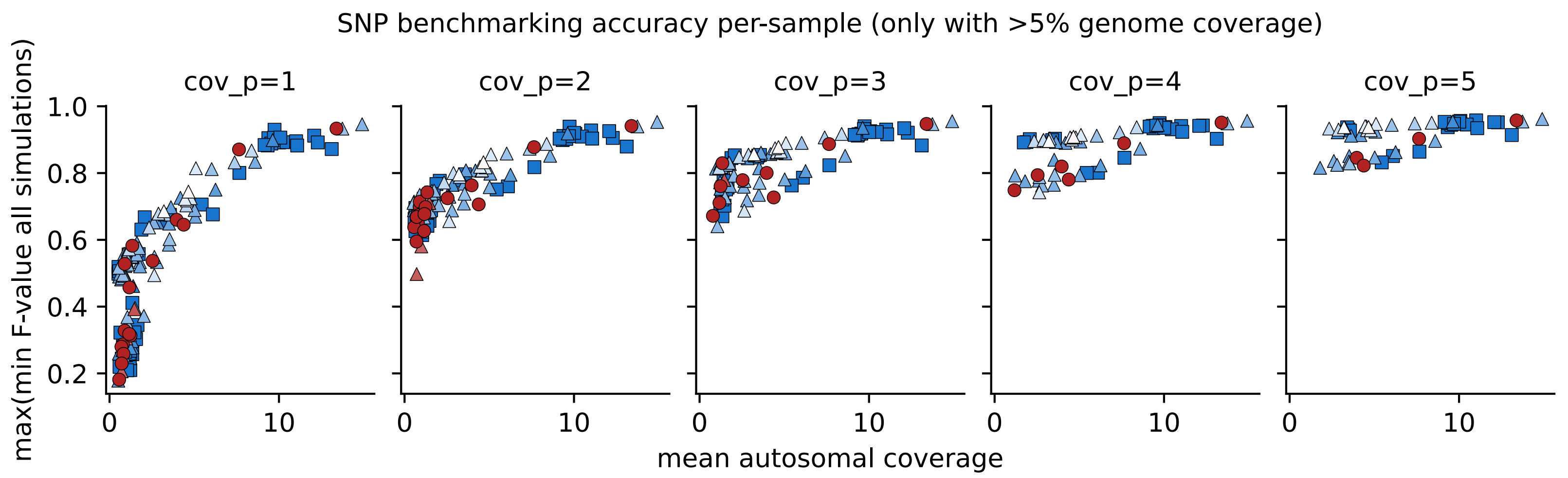

SNP benchmarking accuracy per-sample (only with >5% genome coverage)
cov_p=1
cov_p=2
cov_p=3
cov_p=4
cov_p=5
max(min F-value all simulations)
mean autosomal coverage
1.0
0.8
0.6
0.4
0.2
0
10

**Extended Data Fig. 7. Validation that coverage is the major constraint of SNP calling performance.** Correlation between mean autosomal coverage per sample and SNP calling performance in simulations (F-value) for different cov_p values (minimum coverage per position in all samples). The colors and shapes are equivalent to those in **Fig. 1b**.

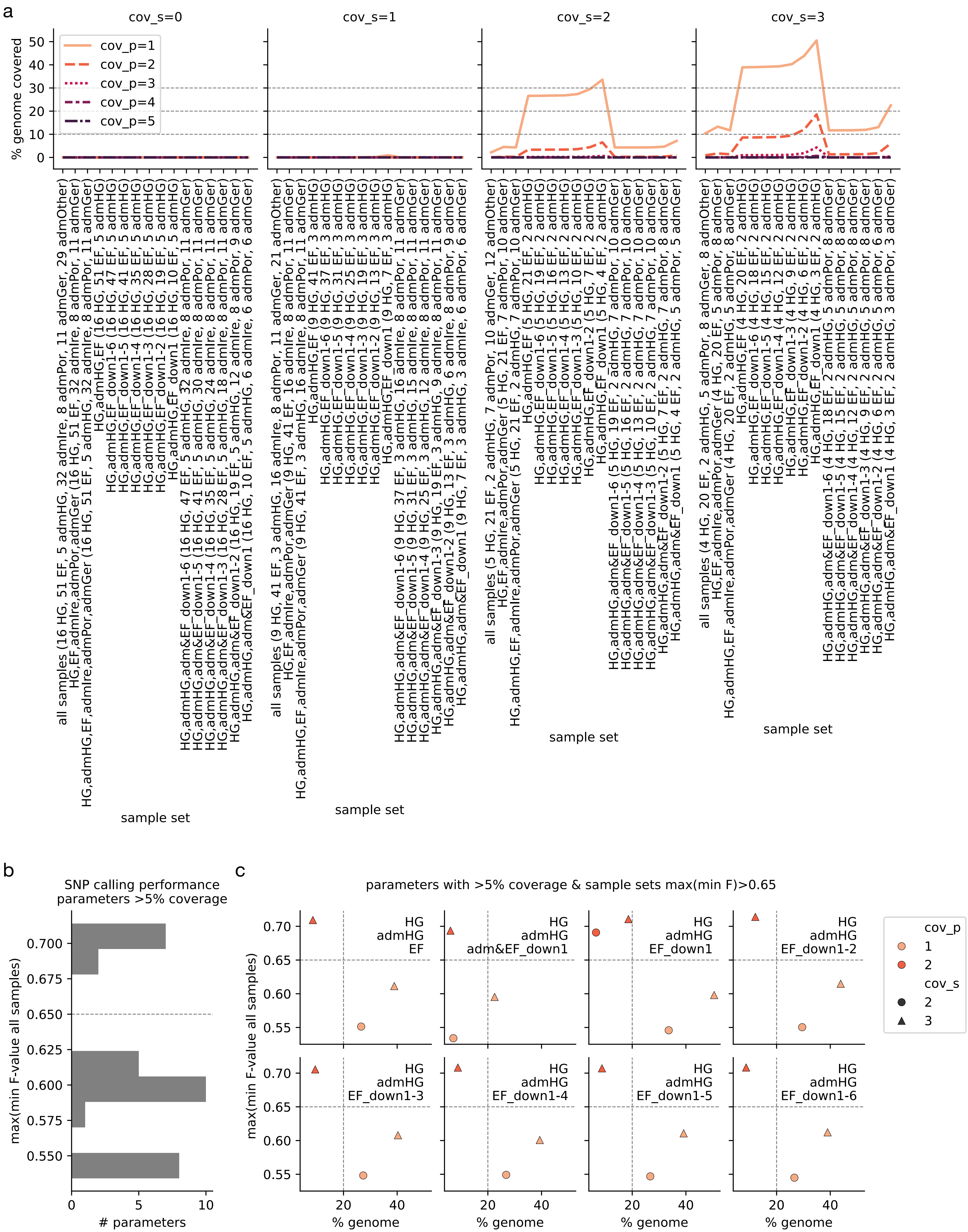
a
cov_s=0
cov_s=1
cov_s=2
cov_s=3
% genome covered
50
40
30
20
10
0
cov_p=1
cov_p=2
cov_p=3
cov_p=4
cov_p=5
all samples (16 HG, 51 EF, 5 admHG, 32 admIre, 8 admPor, 11 admGer, 29 admOther)
HG,EF,admIre,admPor,admGer (16 HG, 51 EF, 32 admIre, 8 admPor, 11 admGer)
HG,admHG,EF,admIre,admPor,admGer (16 HG, 51 EF, 5 admHG, 32 admIre, 8 admPor, 11 admGer)
HG,admHG,EF (16 HG, 51 EF, 5 admHG)
HG,admHG,EF_down1-6 (16 HG, 47 EF, 5 admHG)
HG,admHG,EF_down1-5 (16 HG, 41 EF, 5 admHG)
HG,admHG,EF_down1-4 (16 HG, 35 EF, 5 admHG)
HG,admHG,EF_down1-3 (16 HG, 28 EF, 5 admHG)
HG,admHG,EF_down1-2 (16 HG, 19 EF, 5 admHG)
HG,admHG,EF_down1 (16 HG, 10 EF, 5 admHG)
HG,admHG,adm&EF_down1-6 (16 HG, 47 EF, 5 admHG, 32 admIre, 8 admPor, 11 admGer)
HG,admHG,adm&EF_down1-5 (16 HG, 41 EF, 5 admHG, 30 admIre, 8 admPor, 11 admGer)
HG,admHG,adm&EF_down1-4 (16 HG, 35 EF, 5 admHG, 24 admIre, 8 admPor, 11 admGer)
HG,admHG,adm&EF_down1-3 (16 HG, 28 EF, 5 admHG, 18 admIre, 8 admPor, 11 admGer)
HG,admHG,adm&EF_down1-2 (16 HG, 19 EF, 5 admHG, 12 admIre, 8 admPor, 9 admGer)
HG,admHG,adm&EF_down1 (16 HG, 10 EF, 5 admHG, 6 admIre, 6 admPor, 6 admGer)
sample set
all samples (9 HG, 41 EF, 3 admHG, 16 admIre, 8 admPor, 11 admGer, 21 admOther)
HG,EF,admIre,admPor,admGer (9 HG, 41 EF, 16 admIre, 8 admPor, 11 admGer)
HG,admHG,EF,admIre,admPor,admGer (9 HG, 41 EF, 3 admHG, 16 admIre, 8 admPor, 11 admGer)
HG,admHG,EF (9 HG, 41 EF, 3 admHG)
HG,admHG,EF_down1-6 (9 HG, 37 EF, 3 admHG)
HG,admHG,EF_down1-5 (9 HG, 31 EF, 3 admHG)
HG,admHG,EF_down1-4 (9 HG, 25 EF, 3 admHG)
HG,admHG,EF_down1-3 (9 HG, 19 EF, 3 admHG)
HG,admHG,EF_down1-2 (9 HG, 13 EF, 3 admHG)
HG,admHG,EF_down1 (9 HG, 7 EF, 3 admHG)
HG,admHG,adm&EF_down1-6 (9 HG, 37 EF, 3 admHG, 16 admIre, 8 admPor, 11 admGer)
HG,admHG,adm&EF_down1-5 (9 HG, 31 EF, 3 admHG, 15 admIre, 8 admPor, 11 admGer)
HG,admHG,adm&EF_down1-4 (9 HG, 25 EF, 3 admHG, 12 admIre, 8 admPor, 11 admGer)
HG,admHG,adm&EF_down1-3 (9 HG, 19 EF, 3 admHG, 9 admIre, 8 admPor, 11 admGer)
HG,admHG,adm&EF_down1-2 (9 HG, 13 EF, 3 admHG, 6 admIre, 8 admPor, 9 admGer)
HG,admHG,adm&EF_down1 (9 HG, 7 EF, 3 admHG, 3 admIre, 6 admPor, 6 admGer)
sample set
all samples (5 HG, 21 EF, 2 admHG, 7 admIre, 7 admPor, 10 admGer, 12 admOther)
HG,EF,admIre,admPor,admGer (5 HG, 21 EF, 7 admIre, 7 admPor, 10 admGer)
HG,admHG,EF,admIre,admPor,admGer (5 HG, 21 EF, 2 admHG, 7 admIre, 7 admPor, 10 admGer)
HG,admHG,EF (5 HG, 21 EF, 2 admHG)
HG,admHG,EF_down1-6 (5 HG, 19 EF, 2 admHG)
HG,admHG,EF_down1-5 (5 HG, 16 EF, 2 admHG)
HG,admHG,EF_down1-4 (5 HG, 13 EF, 2 admHG)
HG,admHG,EF_down1-3 (5 HG, 10 EF, 2 admHG)
HG,admHG,EF_down1-2 (5 HG, 7 EF, 2 admHG)
HG,admHG,EF_down1 (5 HG, 4 EF, 2 admHG)
HG,admHG,adm&EF_down1-6 (5 HG, 19 EF, 2 admHG, 7 admIre, 7 admPor, 10 admGer)
HG,admHG,adm&EF_down1-5 (5 HG, 16 EF, 2 admHG, 7 admIre, 7 admPor, 10 admGer)
HG,admHG,adm&EF_down1-4 (5 HG, 13 EF, 2 admHG, 7 admIre, 7 admPor, 10 admGer)
HG,admHG,adm&EF_down1-3 (5 HG, 10 EF, 2 admHG, 7 admIre, 7 admPor, 10 admGer)
HG,admHG,adm&EF_down1-2 (5 HG, 7 EF, 2 admHG, 7 admIre, 7 admPor, 8 admGer)
HG,admHG,adm&EF_down1 (5 HG, 4 EF, 2 admHG, 5 admIre, 5 admPor, 5 admGer)
sample set
all samples (4 HG, 20 EF, 2 admHG, 5 admIre, 5 admPor, 8 admGer, 8 admOther)
HG,EF,admIre,admPor,admGer (4 HG, 20 EF, 5 admIre, 5 admPor, 8 admGer)
HG,admHG,EF,admIre,admPor,admGer (4 HG, 20 EF, 2 admHG, 5 admIre, 5 admPor, 8 admGer)
HG,admHG,EF (4 HG, 20 EF, 2 admHG)
HG,admHG,EF_down1-6 (4 HG, 18 EF, 2 admHG)
HG,admHG,EF_down1-5 (4 HG, 15 EF, 2 admHG)
HG,admHG,EF_down1-4 (4 HG, 12 EF, 2 admHG)
HG,admHG,EF_down1-3 (4 HG, 9 EF, 2 admHG)
HG,admHG,EF_down1-2 (4 HG, 6 EF, 2 admHG)
HG,admHG,EF_down1 (4 HG, 3 EF, 2 admHG)
HG,admHG,adm&EF_down1-6 (4 HG, 18 EF, 2 admHG, 5 admIre, 5 admPor, 8 admGer)
HG,admHG,adm&EF_down1-5 (4 HG, 15 EF, 2 admHG, 5 admIre, 5 admPor, 8 admGer)
HG,admHG,adm&EF_down1-4 (4 HG, 12 EF, 2 admHG, 5 admIre, 5 admPor, 8 admGer)
HG,admHG,adm&EF_down1-3 (4 HG, 9 EF, 2 admHG, 5 admIre, 5 admPor, 8 admGer)
HG,admHG,adm&EF_down1-2 (4 HG, 6 EF, 2 admHG, 5 admIre, 5 admPor, 6 admGer)
HG,admHG,adm&EF_down1 (4 HG, 3 EF, 2 admHG, 3 admIre, 3 admPor, 3 admGer)
sample set
b
SNP calling performance
parameters >5% coverage
max(min F-value all samples)
0.700
0.675
0.650
0.625
0.600
0.575
0.550
0
5
10
### parameters
c
parameters with >5% coverage & sample sets max(min F)>0.65
max(min F-value all samples)
0.70
0.65
0.60
0.55
HG
admHG
EF
HG
admHG
adm&EF_down1
HG
admHG
EF_down1
HG
admHG
EF_down1-2
HG
admHG
EF_down1-3
HG
admHG
EF_down1-4
HG
admHG
EF_down1-5
HG
admHG
EF_down1-6
20
40
% genome
cov_p
1
2
cov_s
2
3

**Extended Data Fig. 8. Benchmarking of SNP calling parameters. a.** To benchmark SNP calling parameters we first checked, for different cov_s (minimum average coverage per sample) and cov_p (minimum coverage per position in all samples) values, the percent of the genome covered (y-axis) across different sample sets (x-axis). These sample subsets include different combinations of HG, EF, admIre (nIre population), admPor (nPor population), admGer (nGer population), admHG (admixed individuals with proportion EF < 0.55), EF_down1-<n> (n subsets of genetically-balanced EF individuals, see **Extended Data Fig. 9**) and adm&EF_down1-<n> (n subsets of genetically-balanced EF and admixed individuals from nIre, nPor and nGer, see **Extended Data Fig. 9**). **b.** Distribution of the F-value (harmonic mean between precision and recall), calculated from the overlap between called and expected (inserted during read simulation) SNPs by different cov_s, cov_p and sample_set parameters. Only parameter combinations yielding >5% genome coverage were considered. The horizontal line indicates the threshold used to identify parameters yielding acceptable performance. **c.** Percent of genome coverage vs calling performance (F-value), for the sample sets yielding some cov_s / cov_p with acceptable performance (F-value > 0.65).

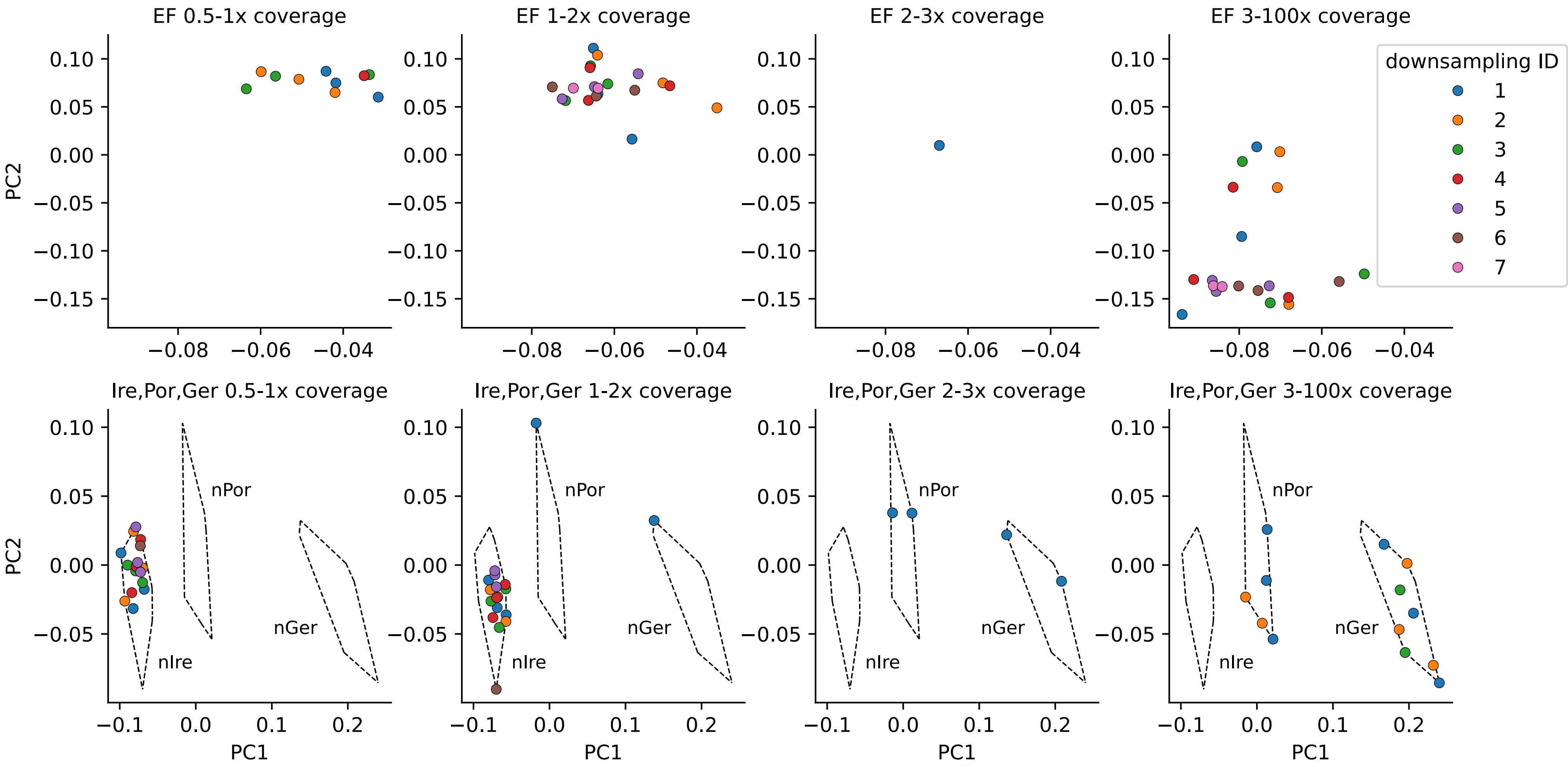

EF 0.5-1x coverage
EF 1-2x coverage
EF 2-3x coverage
EF 3-100x coverage
downsampling ID
1
2
3
4
5
6
7
PC2
PC1
Ire,Por,Ger 0.5-1x coverage
Ire,Por,Ger 1-2x coverage
Ire,Por,Ger 2-3x coverage
Ire,Por,Ger 3-100x coverage
nPor
nGer
nIre

**Extended Data Fig. 9. Definition of downsampled populations for SNP calling benchmarking.** To define EF_down1-<n> and adm&EF_down1-<n> sample subsets for SNP calling parameter benchmarking (see **Extended Data Fig. 8a**), we implemented a subsampling strategy involving the splitting of samples (EF and admixed ones) homogeneously across the first two PCs, and stratifying across mean coverage bins. These plots show the PCA positioning of EF and admixed samples, depending on their mean autosomal coverage bins (columns) and type of samples (EF or admixed, in rows).

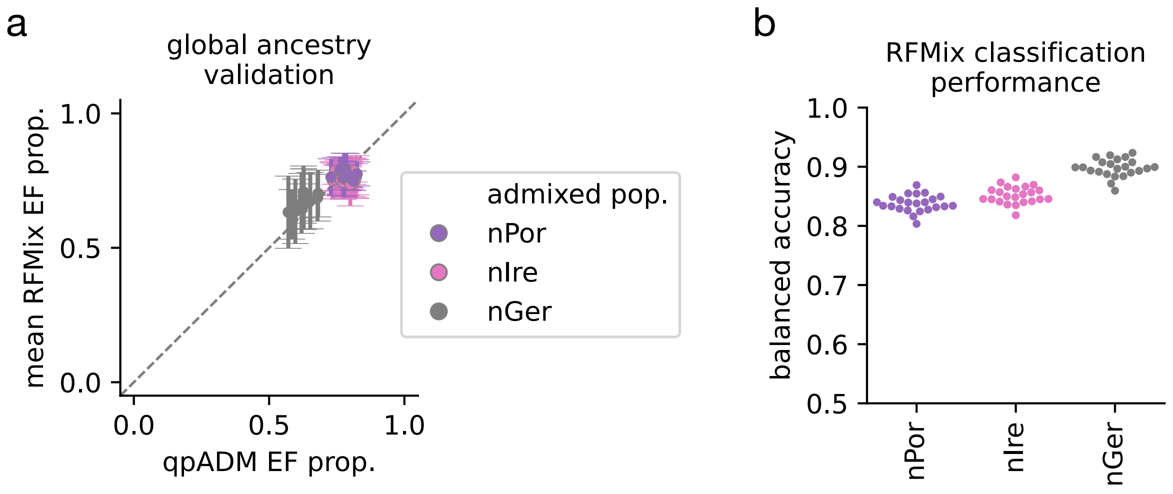

a
global ancestry validation
mean RFMix EF prop.
qpADM EF prop.
1.0
0.5
0.0
0.0
0.5
1.0
admixed pop.
nPor
nIre
nGer
b
RFMix classification performance
balanced accuracy
1.0
0.9
0.8
0.7
0.6
0.5
nPor
nIre
nGer

**Extended Data Fig. 10. Validation of the accuracy of RFMix for adaptive admixture scans. a.** Global EF ancestry proportion of admixed individuals when calculated by qpAdm (x-axis) or RFMix (y-axis). The y values include the mean and standard deviation across chromosomes. **b.** Balanced accuracy of the Local Ancestry Tracts (LAT) prediction for each of the admixed populations, as provided by RFMix.

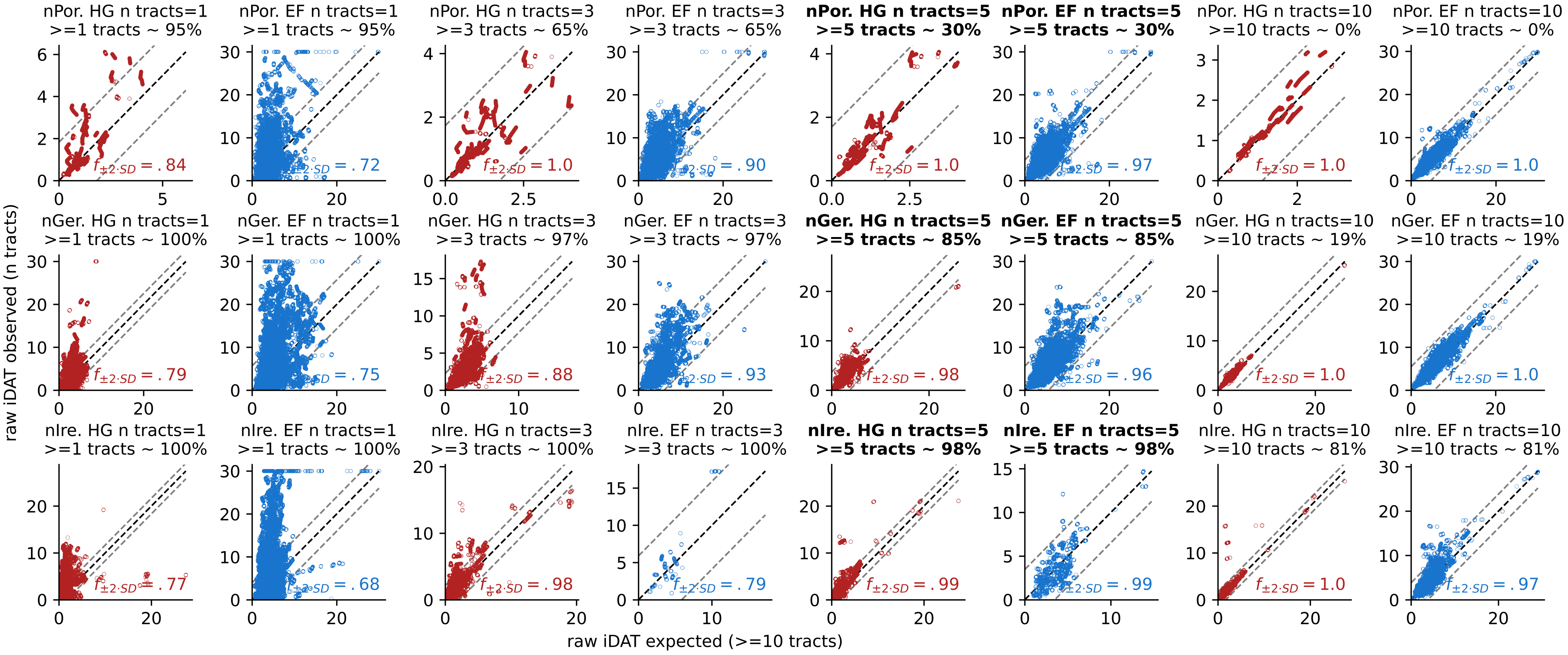

nPor. HG n tracts=1 >=1 tracts ~ 95%
nPor. EF n tracts=1 >=1 tracts ~ 95%
nPor. HG n tracts=3 >=3 tracts ~ 65%
nPor. EF n tracts=3 >=3 tracts ~ 65%
nPor. HG n tracts=5 >=5 tracts ~ 30%
nPor. EF n tracts=5 >=5 tracts ~ 30%
nPor. HG n tracts=10 >=10 tracts ~ 0%
nPor. EF n tracts=10 >=10 tracts ~ 0%
nGer. HG n tracts=1 >=1 tracts ~ 100%
nGer. EF n tracts=1 >=1 tracts ~ 100%
nGer. HG n tracts=3 >=3 tracts ~ 97%
nGer. EF n tracts=3 >=3 tracts ~ 97%
nGer. HG n tracts=5 >=5 tracts ~ 85%
nGer. EF n tracts=5 >=5 tracts ~ 85%
nGer. HG n tracts=10 >=10 tracts ~ 19%
nGer. EF n tracts=10 >=10 tracts ~ 19%
nIre. HG n tracts=1 >=1 tracts ~ 100%
nIre. EF n tracts=1 >=1 tracts ~ 100%
nIre. HG n tracts=3 >=3 tracts ~ 100%
nIre. EF n tracts=3 >=3 tracts ~ 100%
nIre. HG n tracts=5 >=5 tracts ~ 98%
nIre. EF n tracts=5 >=5 tracts ~ 98%
nIre. HG n tracts=10 >=10 tracts ~ 81%
nIre. EF n tracts=10 >=10 tracts ~ 81%
raw iDAT observed (n tracts)
raw iDAT expected (>=10 tracts)

**Extended Data Fig. 11. Effect of filtering out positions with insufficient LATs for iDAT calculations.** To identify an adequate threshold for a minimum number of Local Ancestry Tracts (LATs) for iDAT calculations, we kept positions with >=10 LATs of a given ancestral population (EF or HG), and calculated how much would the $iDAT_{HG}$ and $iDAT_{EF}$ calculation change if we only considered n=1, 3, 5 or 10 randomly selected LATs. The plots show the correlation between the iDAT measured based on all LATs (x-axis), or based on a n LATs (y-axis). The rows correspond to different admixed populations, and the columns indicate different n values and type of iDAT ($iDAT_{HG}$ in red, $iDAT_{EF}$ in blue). The float values indicate the fraction of sites of the genome where the difference in iDAT measures (based on all, or only n LATs) is within 2 standard deviations of all differences across the genome. In the title the % indicates the percent of the genome kept when only considering positions with at least n LATs for both ancestral populations. In bold we highlight the chosen threshold (n=5).

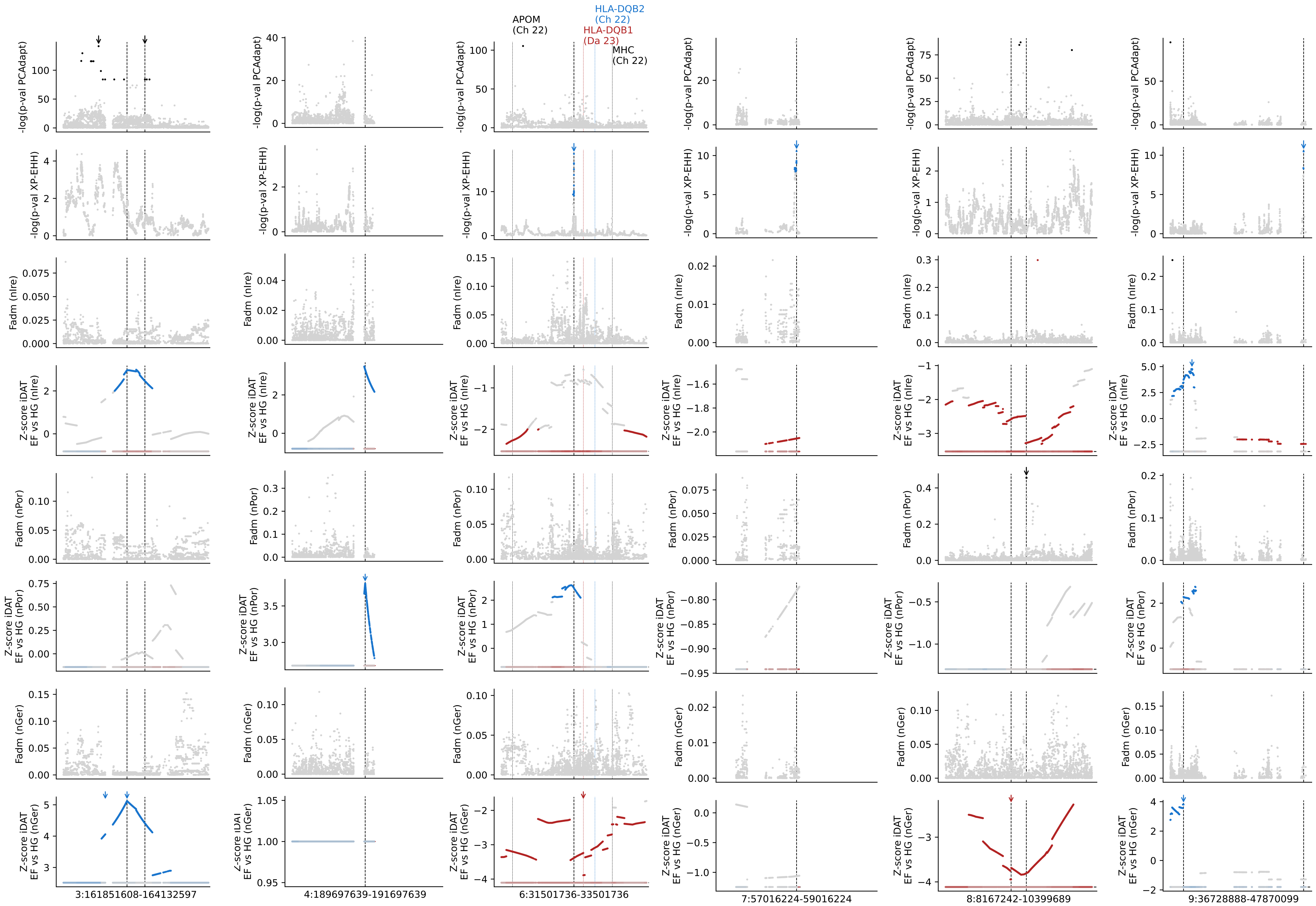

APOM
(Ch 22)
HLA-DQB2
(Ch 22)
HLA-DQB1
(Da 23)
MHC
(Ch 22)
-log(p-val PCAdapt)
-log(p-val XP-EHH)
Fadm (nIre)
Z-score iDAT
EF vs HG (nIre)
Fadm (nPor)
Z-score iDAT
EF vs HG (nPor)
Fadm (nGer)
Z-score iDAT
EF vs HG (nGer)
3:161851608-164132597
4:189697639-191697639
6:31501736-33501736
7:57016224-59016224
8:8167242-10399689
9:36728888-47870099

**Extended Data Fig. 12. First set of six chromosomes with significant hits in multiple selection scans.** This is a zoom into **Fig. 3**, but for regions around the positions with top hits that pass the significance threshold in multiple selection scans. The vertical black lines reflect such positions. The slim vertical lines reflect SNPs that have been found in previous studies of selection during the Mesolithic-Neolithic transition, as in **Extended Data Fig. 14**.
