## Extended figures 13-15 for "Ancient whole genomes reveal regional selection during adaptation to Neolithic lifestyle in Western Eurasia"

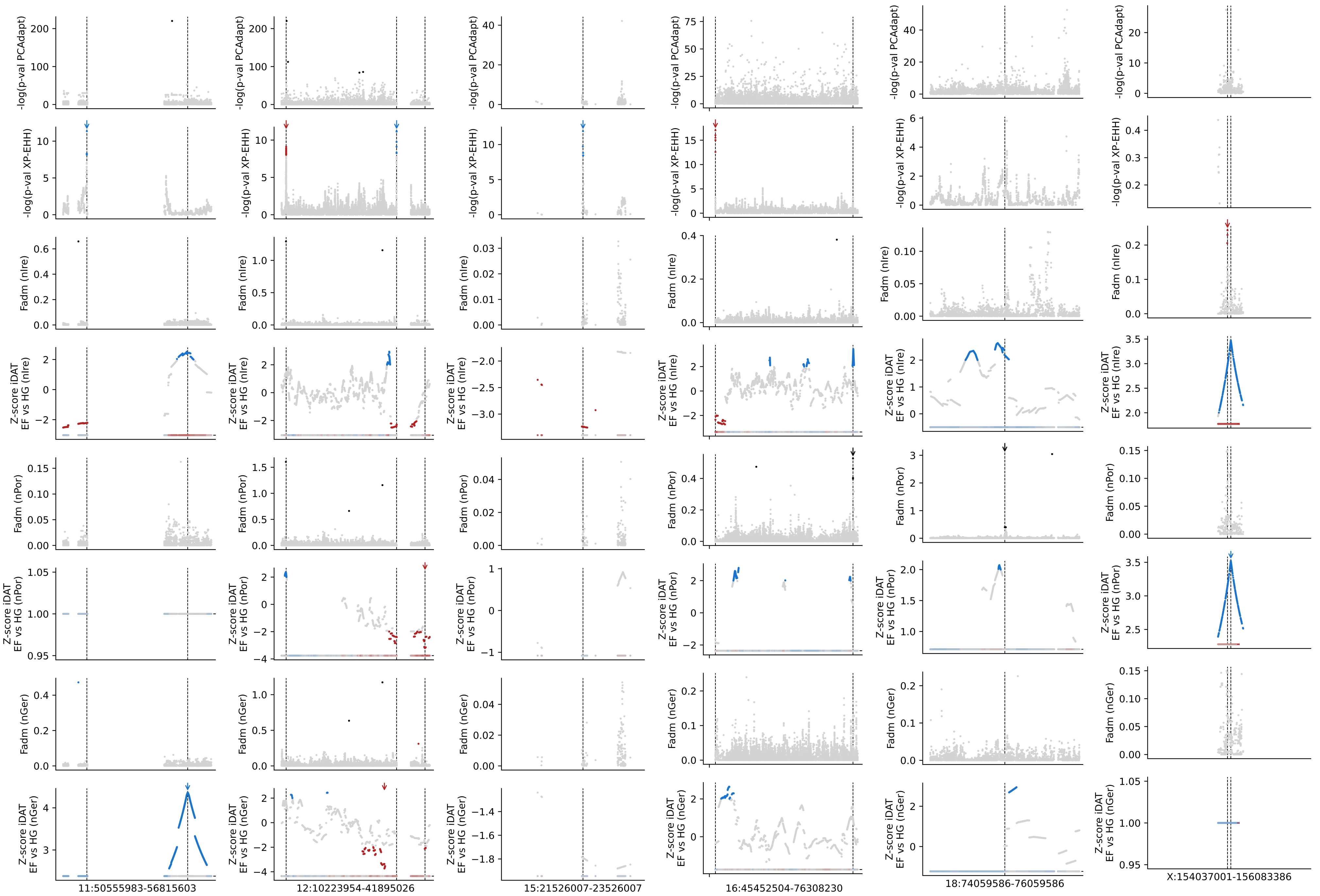
-log(p-val PCAdapt)
-log(p-val XP-EHH)
Fadm (nIre)
Z-score iDAT
EF vs HG (nIre)
Fadm (nPor)
Z-score iDAT
EF vs HG (nPor)
Fadm (nGer)
Z-score iDAT
EF vs HG (nGer)
11:50555983-56815603
12:10223954-41895026
15:21526007-23526007
16:45452504-76308230
18:74059586-76059586
X:154037001-156083386

**Extended Data Fig. 13. Second set of six chromosomes with significant hits in multiple selection scans.** The same as **Extended Data Fig. 13**, but for the remaining six chromosomes.

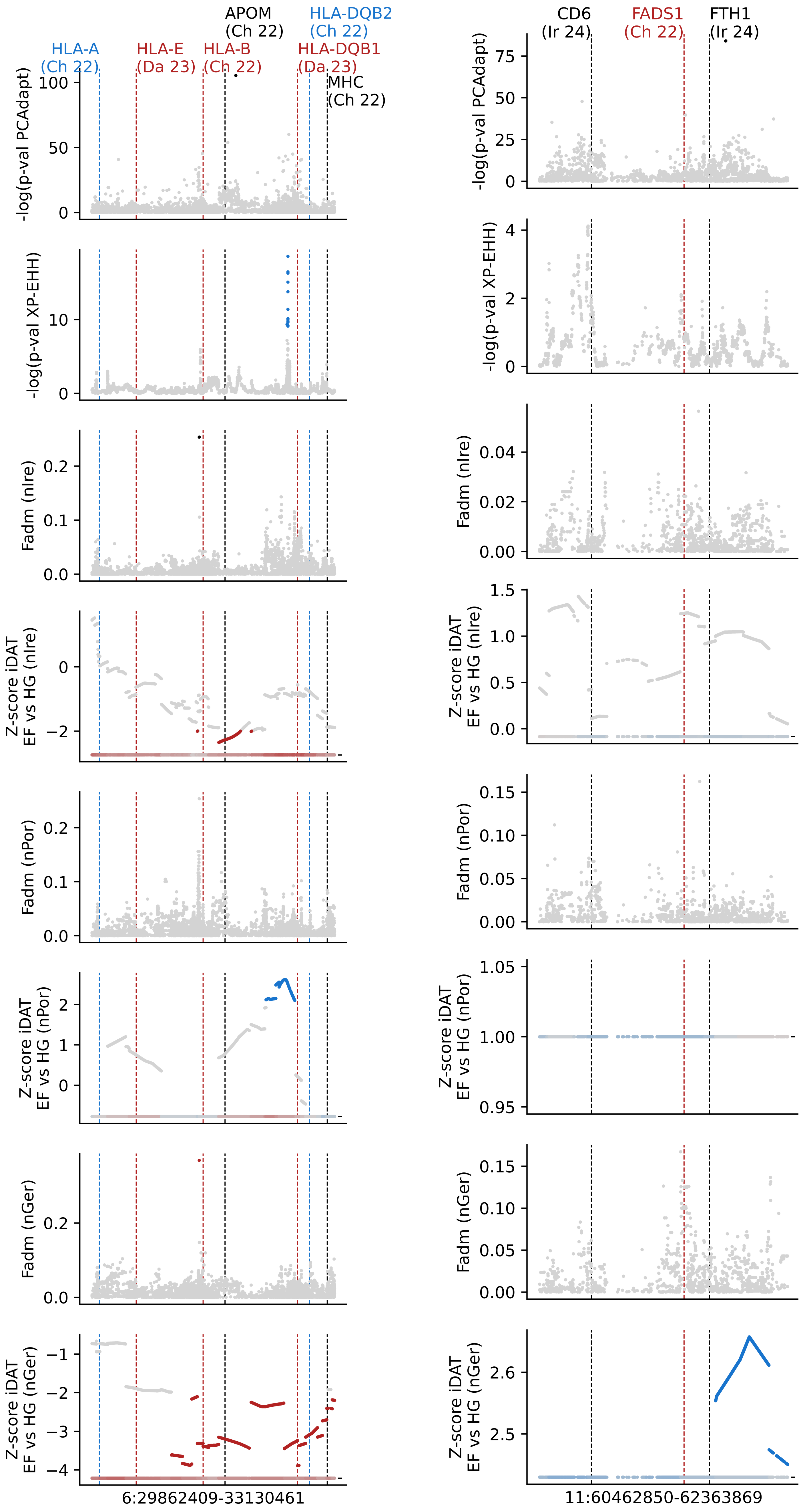

HLA-A (Ch 22)
HLA-E (Da 23)
HLA-B (Ch 22)
APOM (Ch 22)
HLA-DQB1 (Da 23)
HLA-DQB2 (Ch 22)
MHC (Ch 22)
CD6 (Ir 24)
FADS1 (Ch 22)
FTH1 (Ir 24)
-log(p-val PCAdapt)
-log(p-val XP-EHH)
Fadm (nIre)
Z-score iDAT EF vs HG (nIre)
Fadm (nPor)
Z-score iDAT EF vs HG (nPor)
Fadm (nGer)
Z-score iDAT EF vs HG (nGer)
6:29862409-33130461
11:60462850-62363869

**Extended Data Fig. 14. Selection scans on expected regions with overlaps.** This is a zoom into **Fig. 3**, but around positions under selection identified in previous studies. The dashed vertical lines reflect the reported SNP under selection, together with the affected gene and the study of origin (Le 22[6], Ch 22[12], Ir 24[14] and Da 23[13]). The color of the line reflects whether previous selection was described to be favouring EF alleles (blue), HG alleles (red), or whether direction was unknown (black). Note that we only show regions with >1 of these SNPs within <1Mb. Also, these two regions include regions where we did find overlapping selective signatures.

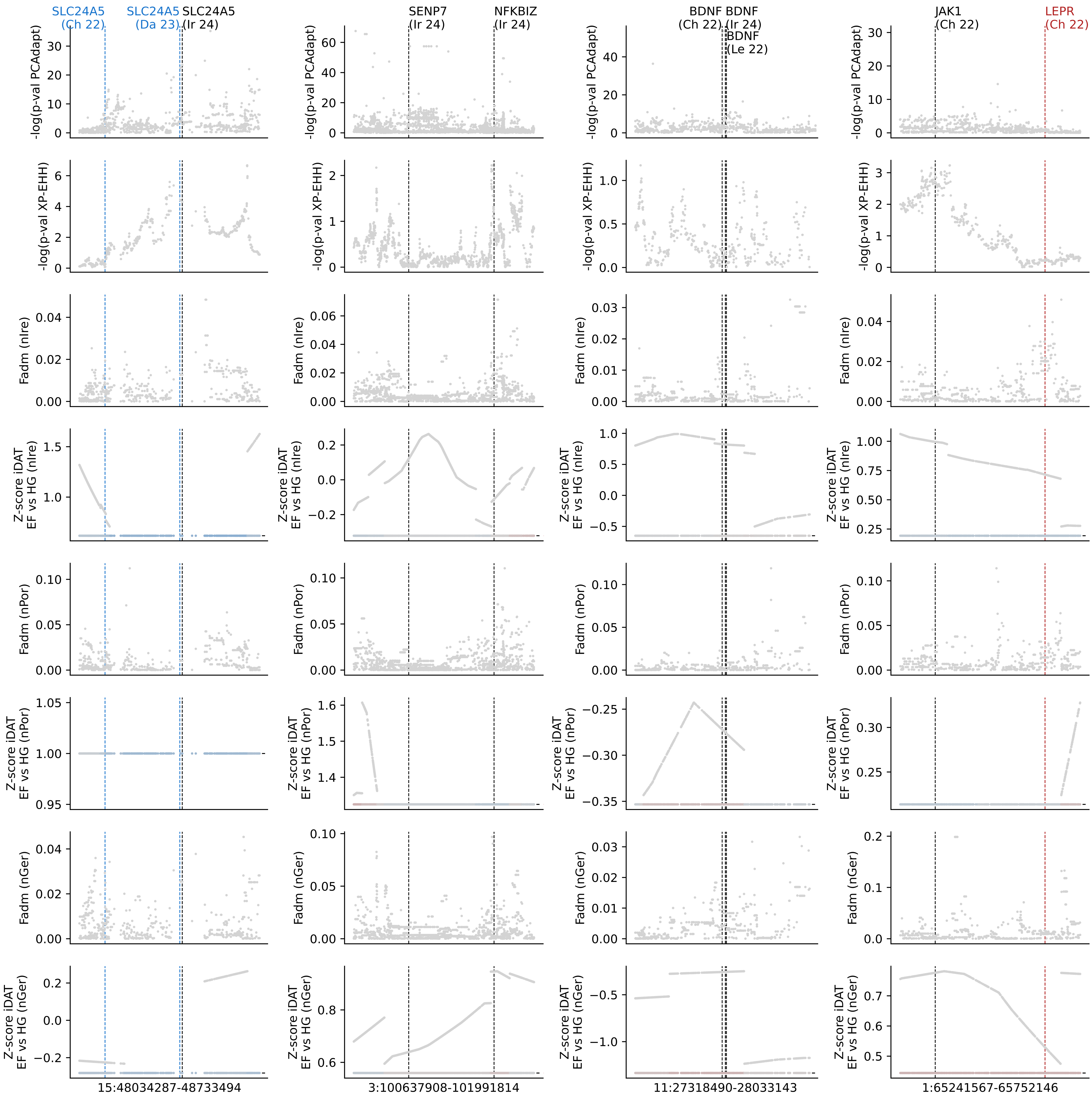
SLC24A5
(Ch 22)
SLC24A5
(Da 23)
SLC24A5
(Ir 24)
SENP7
(Ir 24)
NFKBIZ
(Ir 24)
BDNF
(Ch 22)
BDNF
(Ir 24)
BDNF
(Le 22)
JAK1
(Ch 22)
LEPR
(Ch 22)
-log(p-val PCAdapt)
-log(p-val XP-EHH)
Fadm (nIre)
Z-score iDAT EF vs HG (nIre)
Fadm (nPor)
Z-score iDAT EF vs HG (nPor)
Fadm (nGer)
Z-score iDAT EF vs HG (nGer)
15:48034287-48733494
3:100637908-101991814
11:27318490-28033143
1:65241567-65752146

**Extended Data Fig. 15. Selection scans on expected regions with no overlaps.** The same as **Extended Data Fig. 14**, but for regions where we did not find selective signatures.
