## Extended figures 16-24 for "Ancient whole genomes reveal regional selection during adaptation to Neolithic lifestyle in Western Eurasia"

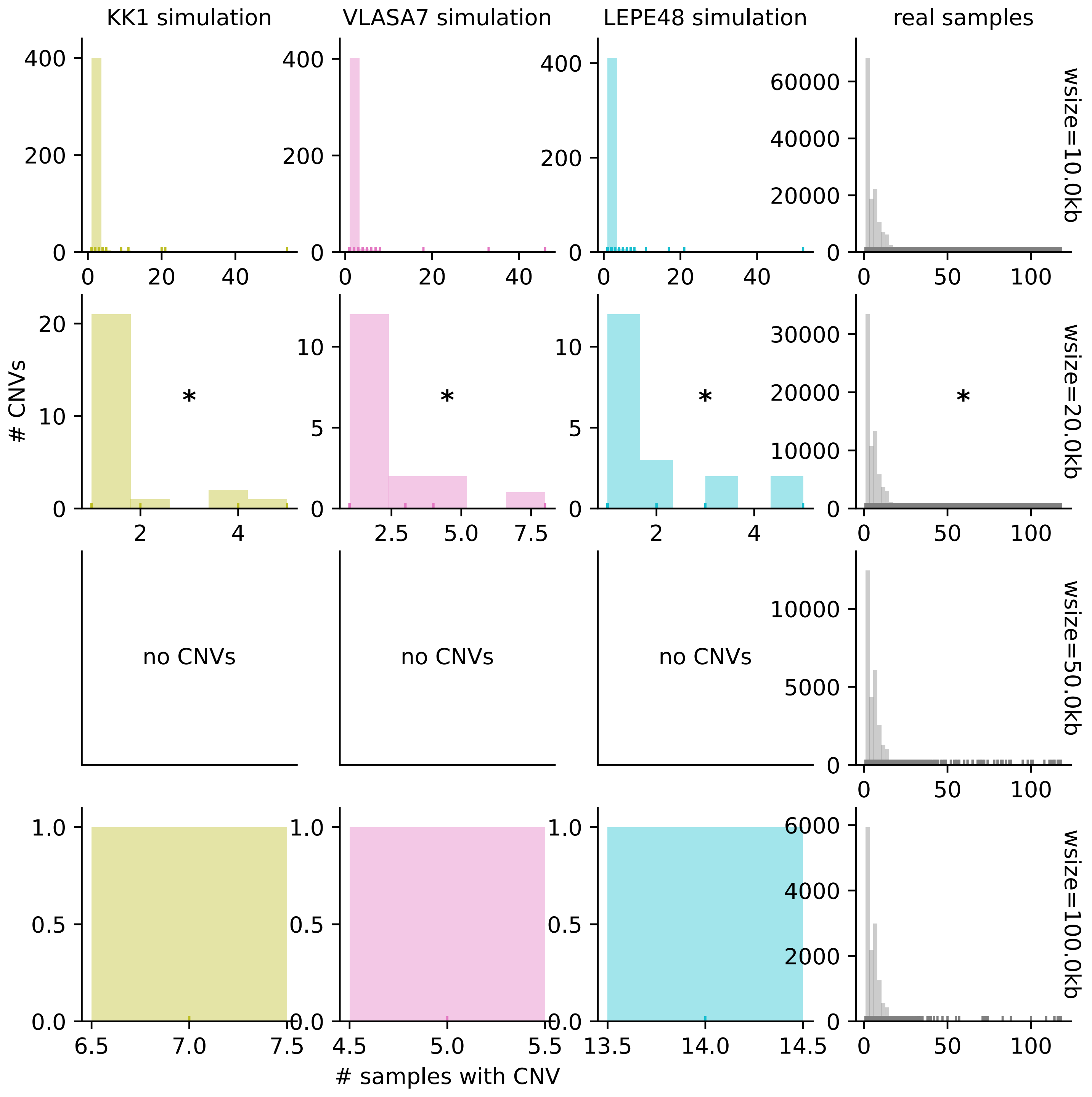

KK1 simulation
VLASA7 simulation
LEPE48 simulation
real samples
wsize=10.0kb
wsize=20.0kb
wsize=50.0kb
wsize=100.0kb
### CNVs
### samples with CNV
no CNVs
no CNVs
no CNVs
*
*
*
*

**Extended Data Fig. 16. Effect of window size on CNV calling results.** To choose a valid window size for CNV calling (rows), we checked the distribution of CNV frequency in each of the negative control simulations (first three columns, where no CNVs are expected), and the real samples (fourth column). The asterisk indicates the chosen window size (20kb).

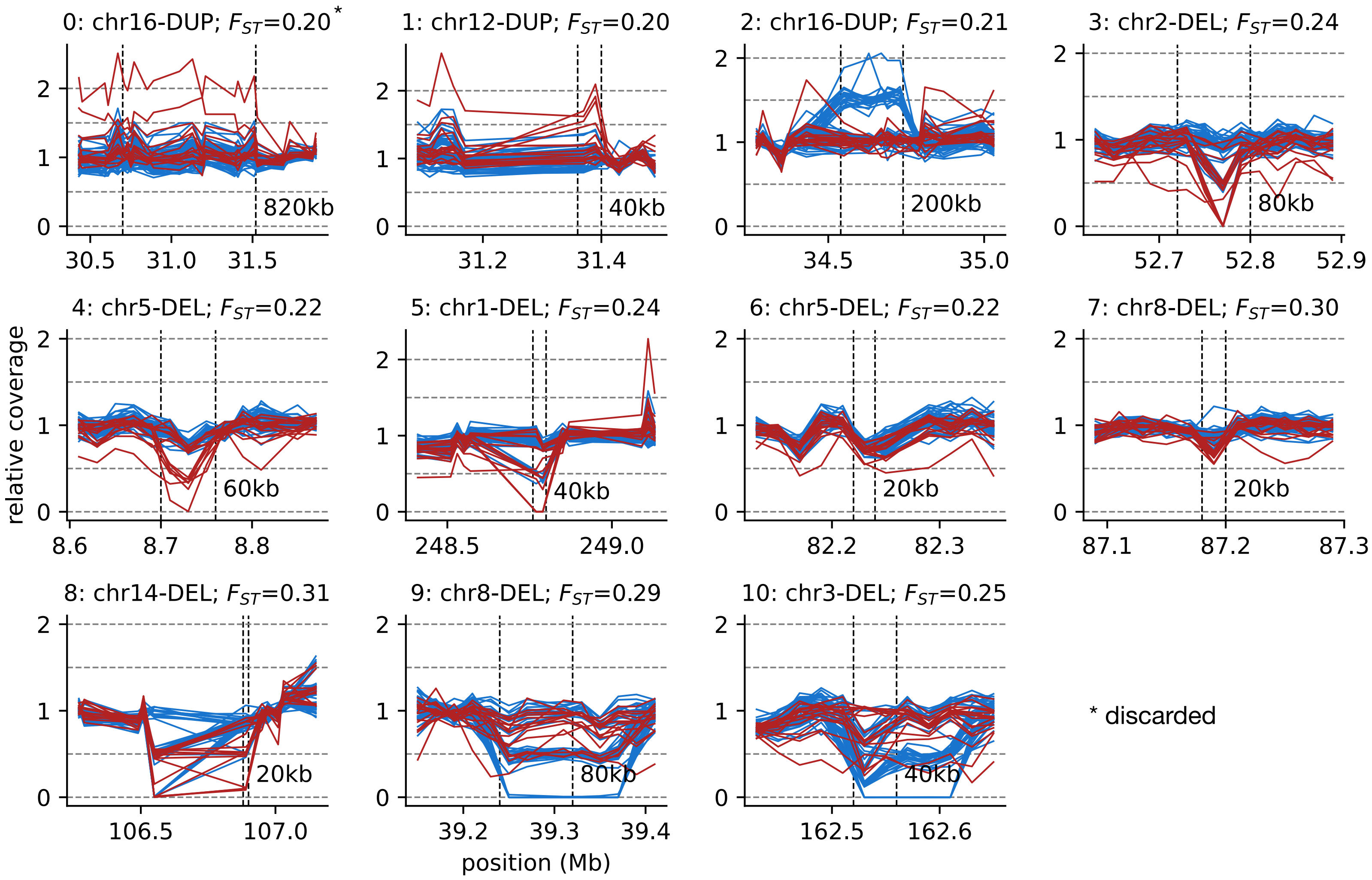
0: chr16-DUP; $F_{ST}$=0.20*
1: chr12-DUP; $F_{ST}$=0.20
2: chr16-DUP; $F_{ST}$=0.21
3: chr2-DEL; $F_{ST}$=0.24
4: chr5-DEL; $F_{ST}$=0.22
5: chr1-DEL; $F_{ST}$=0.24
6: chr5-DEL; $F_{ST}$=0.22
7: chr8-DEL; $F_{ST}$=0.30
8: chr14-DEL; $F_{ST}$=0.31
9: chr8-DEL; $F_{ST}$=0.29
10: chr3-DEL; $F_{ST}$=0.25
820kb
40kb
200kb
80kb
60kb
40kb
20kb
20kb
20kb
80kb
40kb
relative coverage
position (Mb)
* discarded

**Extended Data Fig. 17. All CNVs with differential distribution between the ancestral populations.** For each CNV, in different panels, each line indicates the relative coverage for a given EF (blue) or HG (red) individual around the duplicated or deleted region. The vertical lines indicate the CNV region. We discarded the duplication from the top-left panel, given that the large Fst may be an artifact of noisy coverage of certain HG individuals.

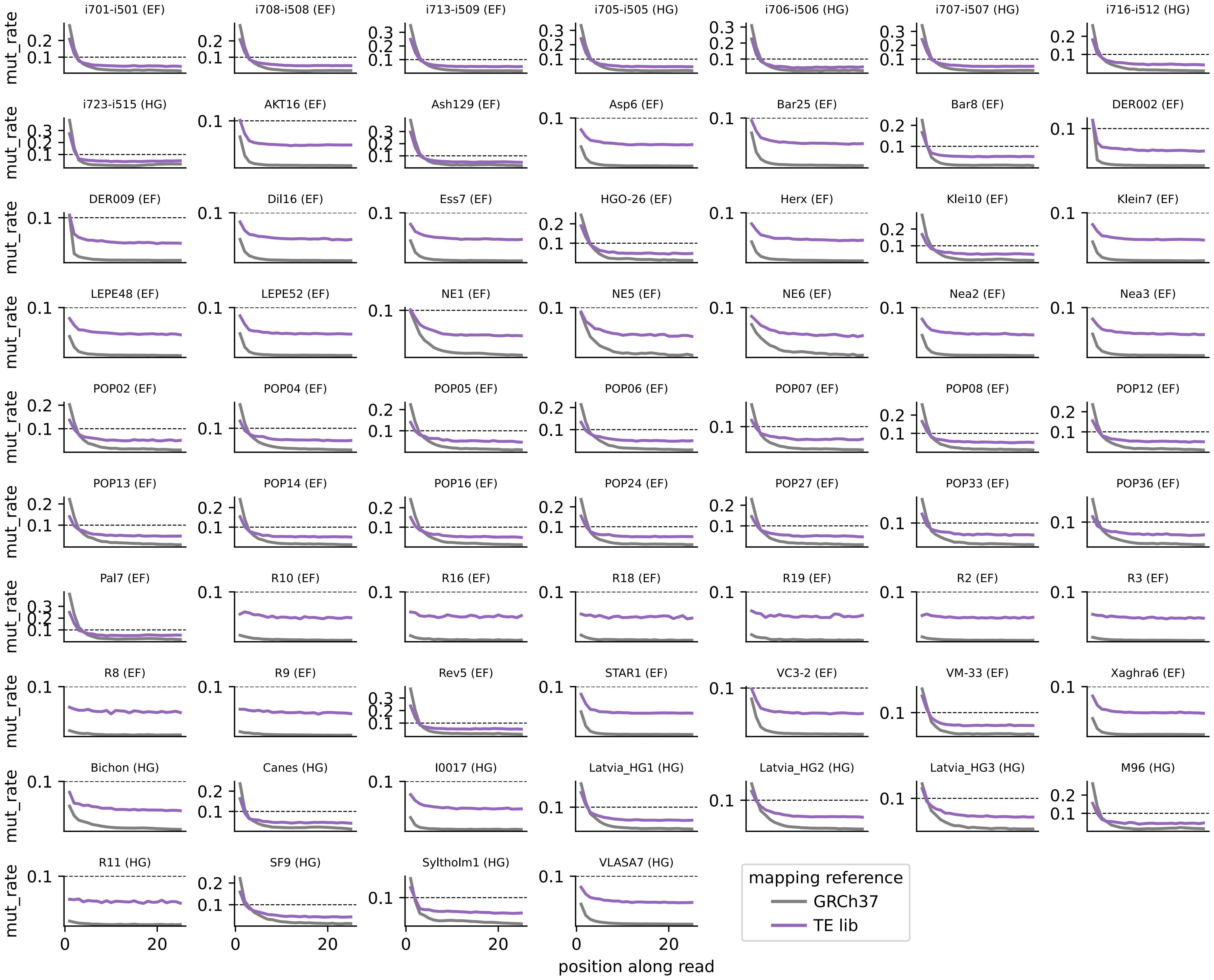

i701-i501 (EF)
i708-i508 (EF)
i713-i509 (EF)
i705-i505 (HG)
i706-i506 (HG)
i707-i507 (HG)
i716-i512 (HG)
i723-i515 (HG)
AKT16 (EF)
Ash129 (EF)
Asp6 (EF)
Bar25 (EF)
Bar8 (EF)
DER002 (EF)
DER009 (EF)
Dil16 (EF)
Ess7 (EF)
HGO-26 (EF)
Herx (EF)
Klei10 (EF)
Klein7 (EF)
LEPE48 (EF)
LEPE52 (EF)
NE1 (EF)
NE5 (EF)
NE6 (EF)
Nea2 (EF)
Nea3 (EF)
POP02 (EF)
POP04 (EF)
POP05 (EF)
POP06 (EF)
POP07 (EF)
POP08 (EF)
POP12 (EF)
POP13 (EF)
POP14 (EF)
POP16 (EF)
POP24 (EF)
POP27 (EF)
POP33 (EF)
POP36 (EF)
Pal7 (EF)
R10 (EF)
R16 (EF)
R18 (EF)
R19 (EF)
R2 (EF)
R3 (EF)
R8 (EF)
R9 (EF)
Rev5 (EF)
STAR1 (EF)
VC3-2 (EF)
VM-33 (EF)
Xaghra6 (EF)
Bichon (HG)
Canes (HG)
I0017 (HG)
Latvia_HG1 (HG)
Latvia_HG2 (HG)
Latvia_HG3 (HG)
M96 (HG)
R11 (HG)
SF9 (HG)
Syltholm1 (HG)
VLASA7 (HG)
mut_rate
position along read
mapping reference
GRCh37
TE lib

**Extended Data Fig. 18**. **Validation of PMD profiles for reads mapping to TE library.** 5’ C>T mutation rate for different positions along the read when mapping the reads against the GRCh37 or the TE library references.

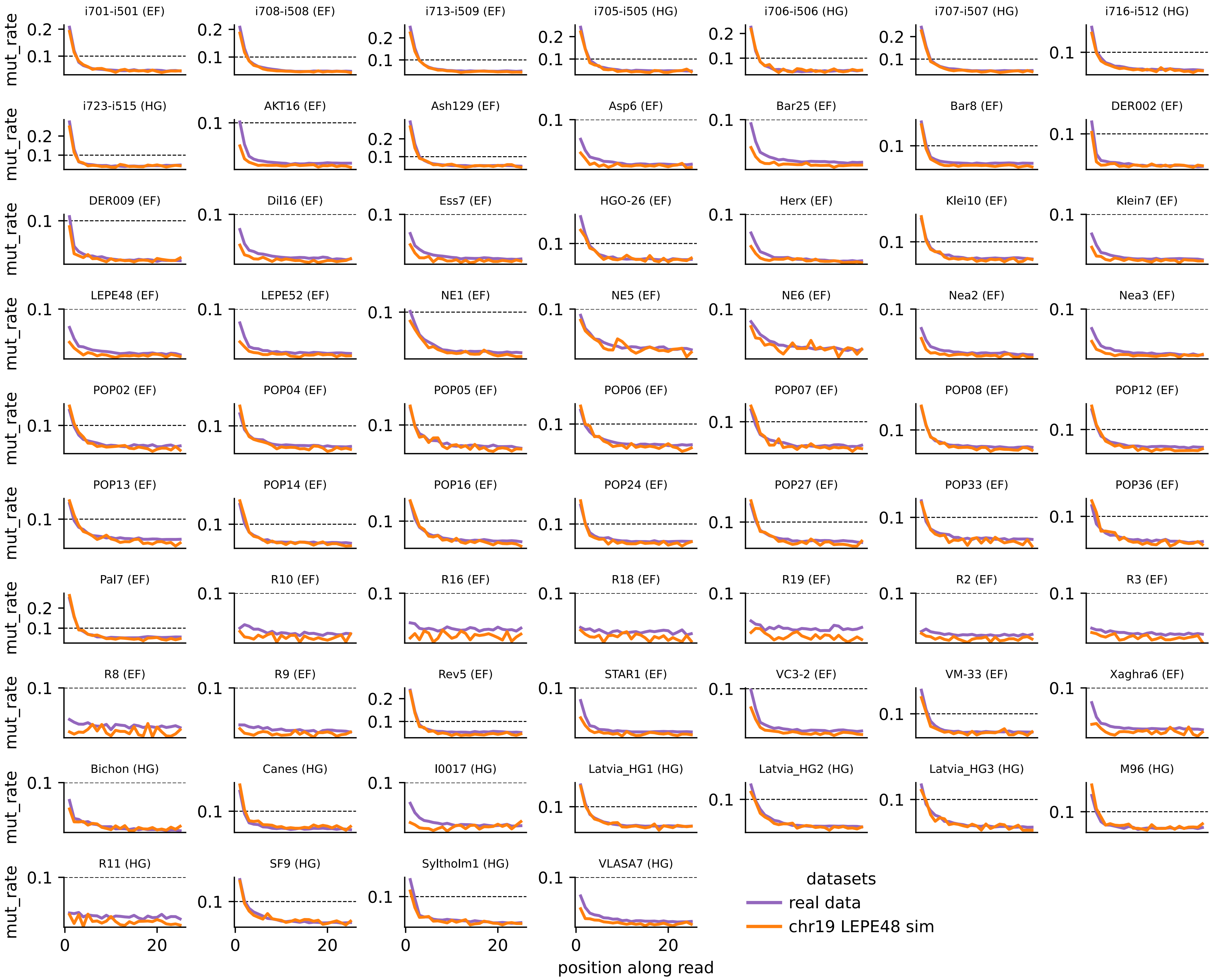

i701-i501 (EF)
i708-i508 (EF)
i713-i509 (EF)
i705-i505 (HG)
i706-i506 (HG)
i707-i507 (HG)
i716-i512 (HG)
i723-i515 (HG)
AKT16 (EF)
Ash129 (EF)
Asp6 (EF)
Bar25 (EF)
Bar8 (EF)
DER002 (EF)
DER009 (EF)
Dil16 (EF)
Ess7 (EF)
HGO-26 (EF)
Herx (EF)
Klei10 (EF)
Klein7 (EF)
LEPE48 (EF)
LEPE52 (EF)
NE1 (EF)
NE5 (EF)
NE6 (EF)
Nea2 (EF)
Nea3 (EF)
POP02 (EF)
POP04 (EF)
POP05 (EF)
POP06 (EF)
POP07 (EF)
POP08 (EF)
POP12 (EF)
POP13 (EF)
POP14 (EF)
POP16 (EF)
POP24 (EF)
POP27 (EF)
POP33 (EF)
POP36 (EF)
Pal7 (EF)
R10 (EF)
R16 (EF)
R18 (EF)
R19 (EF)
R2 (EF)
R3 (EF)
R8 (EF)
R9 (EF)
Rev5 (EF)
STAR1 (EF)
VC3-2 (EF)
VM-33 (EF)
Xaghra6 (EF)
Bichon (HG)
Canes (HG)
I0017 (HG)
Latvia_HG1 (HG)
Latvia_HG2 (HG)
Latvia_HG3 (HG)
M96 (HG)
R11 (HG)
SF9 (HG)
Syltholm1 (HG)
VLASA7 (HG)
mut_rate
0.1
0.2
0
20
position along read
datasets
real data
chr19 LEPE48 sim

**Extended Data Fig. 19. Validation of PMD profiles for reads mapping to TE library, for real or simulated datasets.** 5' C>T mutation rate for different positions along the read when mapping reads against the TE library. The colors indicate the different datasets mapped (real or negative simulations from the LEPE48 SNP set).

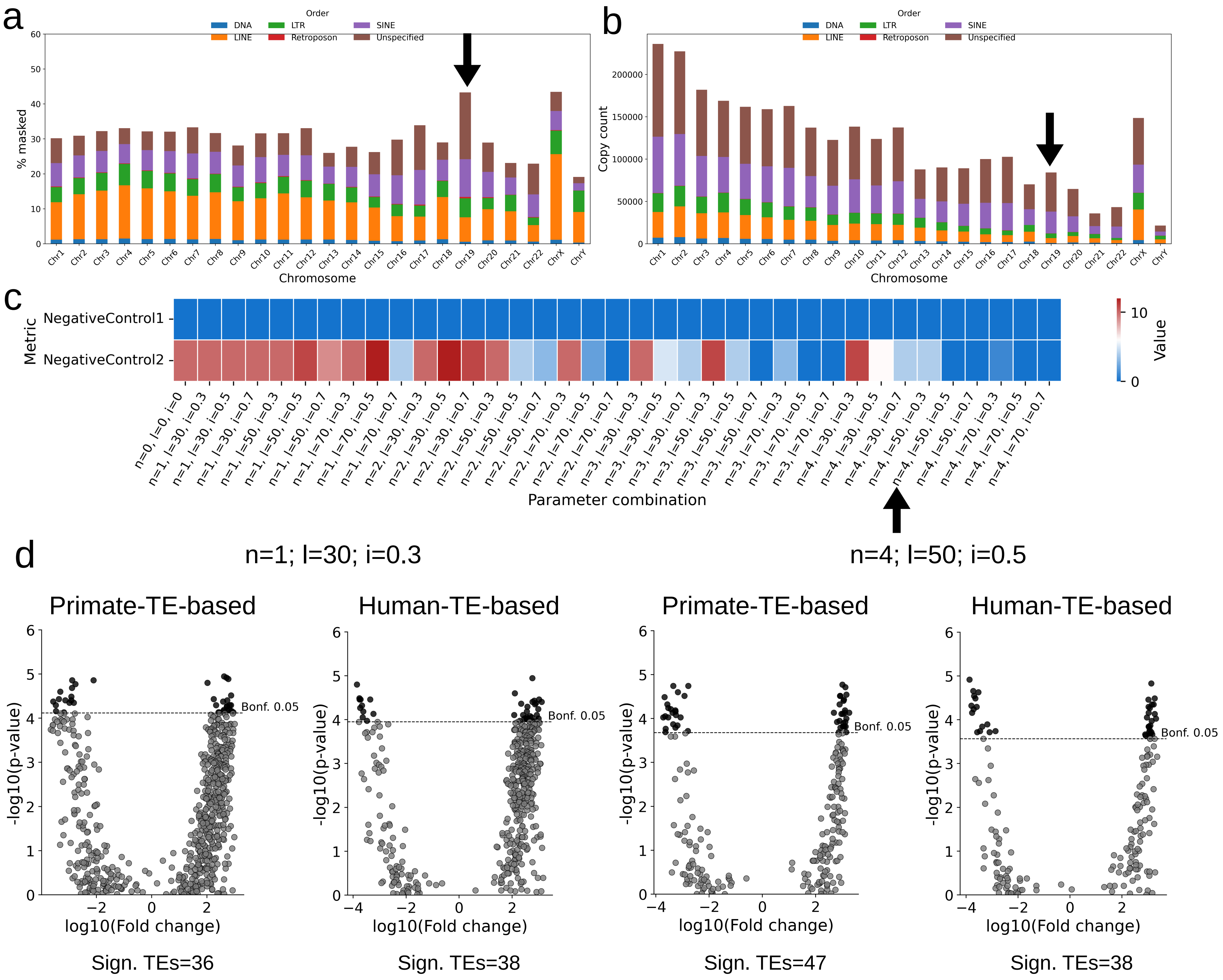

a
b
Order
DNA
LINE
LTR
Retroposon
SINE
Unspecified
% masked
Copy count
Chromosome
c
Metric
NegativeControl1
NegativeControl2
Value
Parameter combination
d
n=1; l=30; i=0.3
n=4; l=50; i=0.5
Primate-TE-based
Human-TE-based
-log10(p-value)
log10(Fold change)
Bonf. 0.05
Sign. TEs=36
Sign. TEs=38
Sign. TEs=47
Sign. TEs=38

**Extended Data Fig. 20. Overall evaluation results of AncienTE negative control datasets. a.** TE proportions per chromosome in the human reference assembly GRCh37. **b.** Number of TE copies per chromosome in the human reference assembly GRCh37. The chromosome 19 (with an arrow) was selected for further simulations. **c.** Heat map representing the number of TE with statistical significance found by AncienTE in the negative dataset 1 (raw short-reads simulated based on the same chromosome 19 sequence), and in the negative control 2 dataset (real short-reads mapped against a single-copy gene library). An arrow indicates the final parameter selection. **d.** Volcano plots showing the results for using two TE libraries: a primate-based and a human-specific (see materials and methods). AncienTE was run using two sets of parameters, a permissive setting (n = 1, l = 30, i = 0.3) and a more stringent setting (n = 4, l = 50, i = 0.5).

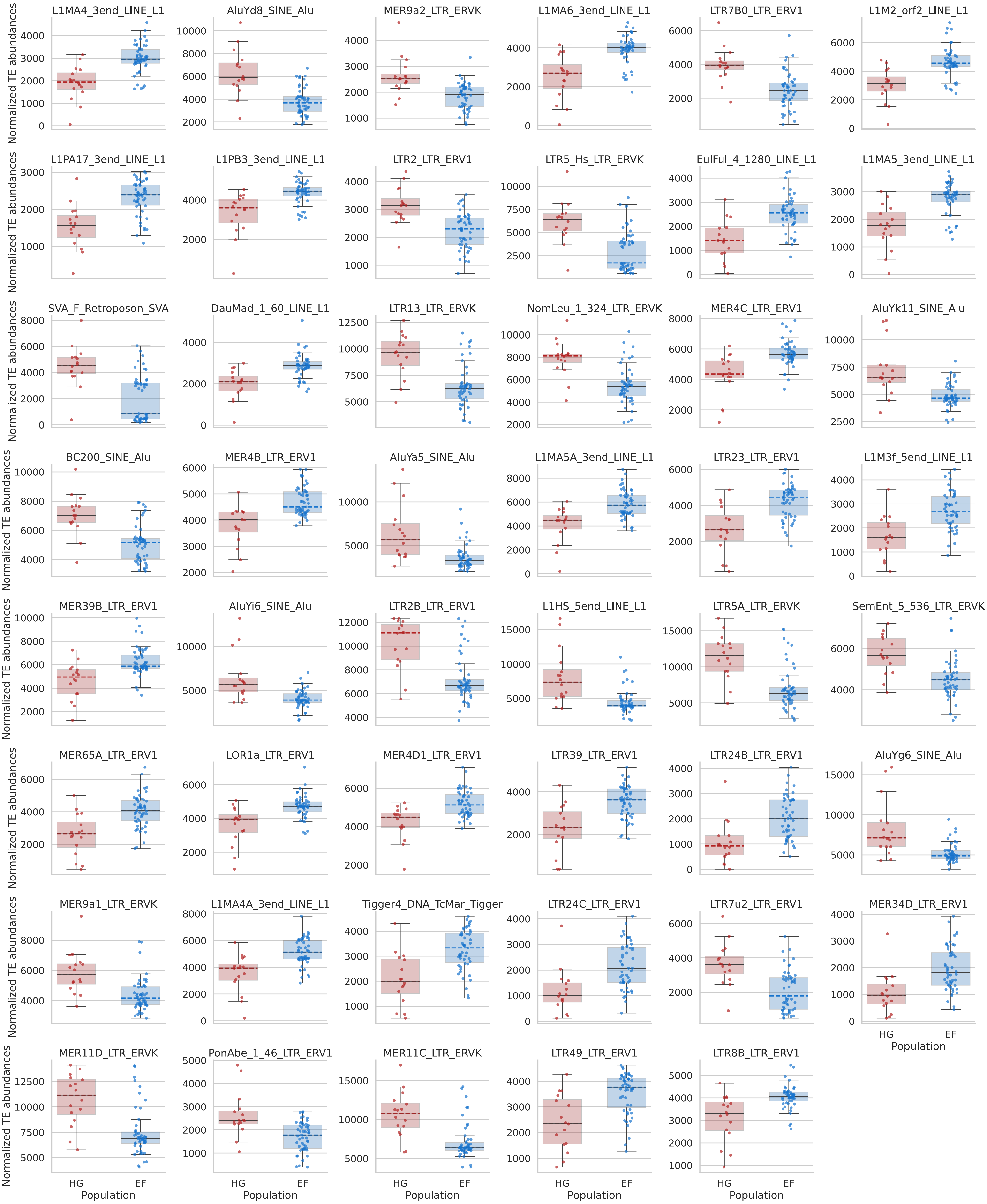

L1MA4_3end_LINE_L1
AluYd8_SINE_Alu
MER9a2_LTR_ERVK
L1MA6_3end_LINE_L1
LTR7B0_LTR_ERV1
L1M2_orf2_LINE_L1
L1PA17_3end_LINE_L1
L1PB3_3end_LINE_L1
LTR2_LTR_ERV1
LTR5_Hs_LTR_ERVK
EulFul_4_1280_LINE_L1
L1MA5_3end_LINE_L1
SVA_F_Retroposon_SVA
DauMad_1_60_LINE_L1
LTR13_LTR_ERVK
NomLeu_1_324_LTR_ERVK
MER4C_LTR_ERV1
AluYk11_SINE_Alu
BC200_SINE_Alu
MER4B_LTR_ERV1
AluYa5_SINE_Alu
L1MA5A_3end_LINE_L1
LTR23_LTR_ERV1
L1M3f_5end_LINE_L1
MER39B_LTR_ERV1
AluYi6_SINE_Alu
LTR2B_LTR_ERV1
L1HS_5end_LINE_L1
LTR5A_LTR_ERVK
SemEnt_5_536_LTR_ERVK
MER65A_LTR_ERV1
LOR1a_LTR_ERV1
MER4D1_LTR_ERV1
LTR39_LTR_ERV1
LTR24B_LTR_ERV1
AluYg6_SINE_Alu
MER9a1_LTR_ERVK
L1MA4A_3end_LINE_L1
Tigger4_DNA_TcMar_Tigger
LTR24C_LTR_ERV1
LTR7u2_LTR_ERV1
MER34D_LTR_ERV1
MER11D_LTR_ERVK
PonAbe_1_46_LTR_ERV1
MER11C_LTR_ERVK
LTR49_LTR_ERV1
LTR8B_LTR_ERV1
Normalized TE abundances
HG
EF
Population

**Extended Data Fig. 21. Significantly differentiated TE families between HG and EF.** Boxplots showing abundance distributions for the 47 statistically differentiated TE families between HG and EF individuals.

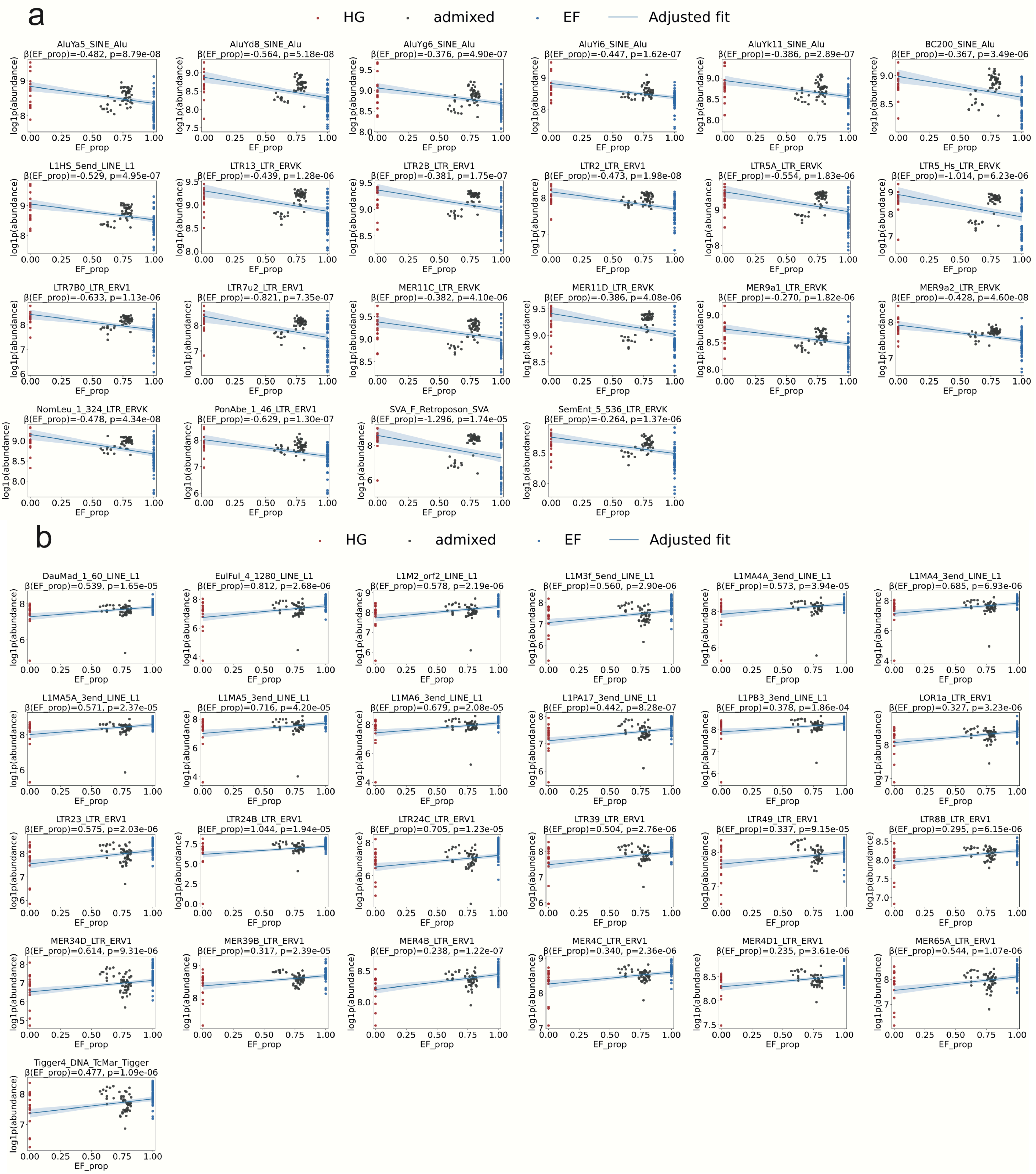
a
HG
admixed
EF
Adjusted fit
AluYa5_SINE_Alu
β(EF_prop)=-0.482, p=8.79e-08
AluYd8_SINE_Alu
β(EF_prop)=-0.564, p=5.18e-08
AluYg6_SINE_Alu
β(EF_prop)=-0.376, p=4.90e-07
AluYi6_SINE_Alu
β(EF_prop)=-0.447, p=1.62e-07
AluYk11_SINE_Alu
β(EF_prop)=-0.386, p=2.89e-07
BC200_SINE_Alu
β(EF_prop)=-0.367, p=3.49e-06
L1HS_5end_LINE_L1
β(EF_prop)=-0.529, p=4.95e-07
LTR13_LTR_ERVK
β(EF_prop)=-0.439, p=1.28e-06
LTR2B_LTR_ERV1
β(EF_prop)=-0.381, p=1.75e-07
LTR2_LTR_ERV1
β(EF_prop)=-0.473, p=1.98e-08
LTR5A_LTR_ERVK
β(EF_prop)=-0.554, p=1.83e-06
LTR5_Hs_LTR_ERVK
β(EF_prop)=-1.014, p=6.23e-06
LTR7B0_LTR_ERV1
β(EF_prop)=-0.633, p=1.13e-06
LTR7u2_LTR_ERV1
β(EF_prop)=-0.821, p=7.35e-07
MER11C_LTR_ERVK
β(EF_prop)=-0.382, p=4.10e-06
MER11D_LTR_ERVK
β(EF_prop)=-0.386, p=4.08e-06
MER9a1_LTR_ERVK
β(EF_prop)=-0.270, p=1.82e-06
MER9a2_LTR_ERVK
β(EF_prop)=-0.428, p=4.60e-08
NomLeu_1_324_LTR_ERVK
β(EF_prop)=-0.478, p=4.34e-08
PonAbe_1_46_LTR_ERV1
β(EF_prop)=-0.629, p=1.30e-07
SVA_F_Retroposon_SVA
β(EF_prop)=-1.296, p=1.74e-05
SemEnt_5_536_LTR_ERVK
β(EF_prop)=-0.264, p=1.37e-06
log1p(abundance)
EF_prop
b
HG
admixed
EF
Adjusted fit
DauMad_1_60_LINE_L1
β(EF_prop)=0.539, p=1.65e-05
EulFul_4_1280_LINE_L1
β(EF_prop)=0.812, p=2.68e-06
L1M2_orf2_LINE_L1
β(EF_prop)=0.578, p=2.19e-06
L1M3f_5end_LINE_L1
β(EF_prop)=0.560, p=2.90e-06
L1MA4A_3end_LINE_L1
β(EF_prop)=0.573, p=3.94e-05
L1MA4_3end_LINE_L1
β(EF_prop)=0.685, p=6.93e-06
L1MA5A_3end_LINE_L1
β(EF_prop)=0.571, p=2.37e-05
L1MA5_3end_LINE_L1
β(EF_prop)=0.716, p=4.20e-05
L1MA6_3end_LINE_L1
β(EF_prop)=0.679, p=2.08e-05
L1PA17_3end_LINE_L1
β(EF_prop)=0.442, p=8.28e-07
L1PB3_3end_LINE_L1
β(EF_prop)=0.378, p=1.86e-04
LOR1a_LTR_ERV1
β(EF_prop)=0.327, p=3.23e-06
LTR23_LTR_ERV1
β(EF_prop)=0.575, p=2.03e-06
LTR24B_LTR_ERV1
β(EF_prop)=1.044, p=1.94e-05
LTR24C_LTR_ERV1
β(EF_prop)=0.705, p=1.23e-05
LTR39_LTR_ERV1
β(EF_prop)=0.504, p=2.76e-06
LTR49_LTR_ERV1
β(EF_prop)=0.337, p=9.15e-05
LTR8B_LTR_ERV1
β(EF_prop)=0.295, p=6.15e-06
MER34D_LTR_ERV1
β(EF_prop)=0.614, p=9.31e-06
MER39B_LTR_ERV1
β(EF_prop)=0.317, p=2.39e-05
MER4B_LTR_ERV1
β(EF_prop)=0.238, p=1.22e-07
MER4C_LTR_ERV1
β(EF_prop)=0.340, p=2.36e-06
MER4D1_LTR_ERV1
β(EF_prop)=0.235, p=3.61e-06
MER65A_LTR_ERV1
β(EF_prop)=0.544, p=1.07e-06
Tigger4_DNA_TcMar_Tigger
β(EF_prop)=0.477, p=1.09e-06
log1p(abundance)
EF_prop

**Extended Data Fig. 22. Linear modelling of TE family abundance as a function of ancestry proportion.** Scatter plots showing the relationship between TE family abundance (log-transformed) and ancestry proportion (ancestry_prop) for the most significantly differentiated TE families. Each point represents one individual. Red and blue points denote individuals with predominantly HG and EF ancestry, respectively, whereas admixed individuals are shown in black. Solid lines represent fitted linear regression models with 95% confidence intervals (shaded areas). To assess whether TE abundance was associated with ancestry independently of sequencing depth and sample age, we fitted a linear model of the form y ~ ancestry_prop + Coverage + Date_BCE, where y corresponds to TE family abundance. The β coefficient for ancestry proportion and corresponding p-value are indicated above each panel. Plots were sorted by TE families being enriched in **a.** HG and in **b.** EF.

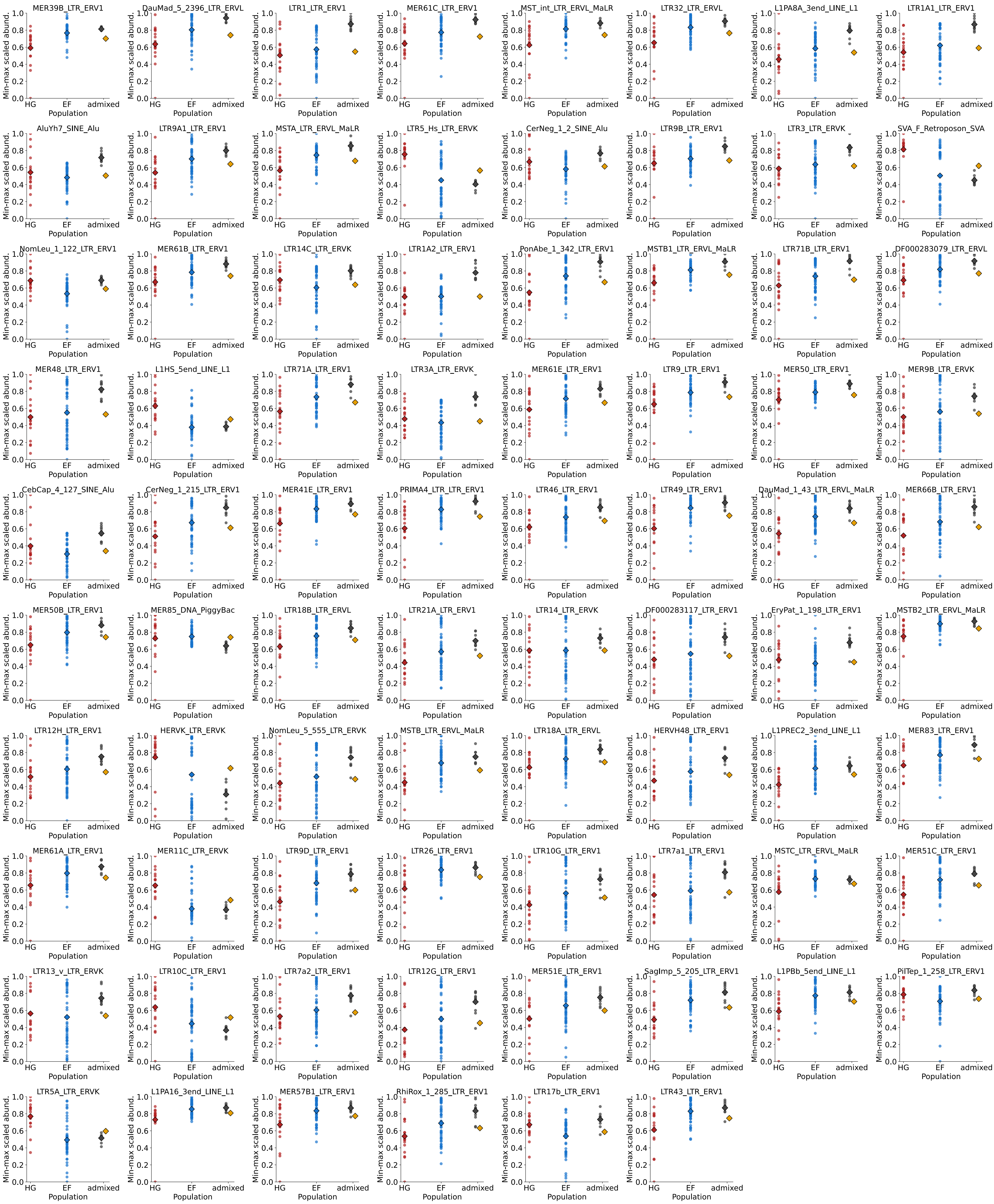
MER39B_LTR_ERV1
DauMad_5_2396_LTR_ERVL
LTR1_LTR_ERV1
MER61C_LTR_ERV1
MST_int_LTR_ERVL_MaLR
LTR32_LTR_ERVL
L1PA8A_3end_LINE_L1
LTR1A1_LTR_ERV1
AluYh7_SINE_Alu
LTR9A1_LTR_ERV1
MSTA_LTR_ERVL_MaLR
LTR5_Hs_LTR_ERVK
CerNeg_1_2_SINE_Alu
LTR9B_LTR_ERV1
LTR3_LTR_ERVK
SVA_F_Retroposon_SVA
NomLeu_1_122_LTR_ERV1
MER61B_LTR_ERV1
LTR14C_LTR_ERVK
LTR1A2_LTR_ERV1
PonAbe_1_342_LTR_ERV1
MSTB1_LTR_ERVL_MaLR
LTR71B_LTR_ERV1
DF000283079_LTR_ERVL
MER48_LTR_ERV1
L1HS_5end_LINE_L1
LTR71A_LTR_ERV1
LTR3A_LTR_ERVK
MER61E_LTR_ERV1
LTR9_LTR_ERV1
MER50_LTR_ERV1
MER9B_LTR_ERVK
CebCap_4_127_SINE_Alu
CerNeg_1_215_LTR_ERV1
MER41E_LTR_ERV1
PRIMA4_LTR_LTR_ERV1
LTR46_LTR_ERV1
LTR49_LTR_ERV1
DauMad_1_43_LTR_ERVL_MaLR
MER66B_LTR_ERV1
MER50B_LTR_ERV1
MER85_DNA_PiggyBac
LTR18B_LTR_ERVL
LTR21A_LTR_ERV1
LTR14_LTR_ERVK
DF000283117_LTR_ERV1
EryPat_1_198_LTR_ERV1
MSTB2_LTR_ERVL_MaLR
LTR12H_LTR_ERV1
HERVK_LTR_ERVK
NomLeu_5_555_LTR_ERVK
MSTB_LTR_ERVL_MaLR
LTR18A_LTR_ERVL
HERVH48_LTR_ERV1
L1PREC2_3end_LINE_L1
MER83_LTR_ERV1
MER61A_LTR_ERV1
MER11C_LTR_ERVK
LTR9D_LTR_ERV1
LTR26_LTR_ERV1
LTR10G_LTR_ERV1
LTR7a1_LTR_ERV1
MSTC_LTR_ERVL_MaLR
MER51C_LTR_ERV1
LTR13_v_LTR_ERVK
LTR10C_LTR_ERV1
LTR7a2_LTR_ERV1
LTR12G_LTR_ERV1
MER51E_LTR_ERV1
SagImp_5_205_LTR_ERV1
L1PBb_5end_LINE_L1
PilTep_1_258_LTR_ERV1
LTR5A_LTR_ERVK
L1PA16_3end_LINE_L1
MER57B1_LTR_ERV1
RhiRox_1_285_LTR_ERV1
LTR17b_LTR_ERV1
LTR43_LTR_ERV1
Min-max scaled abund.
HG
EF
admixed
Population
0.0
0.2
0.4
0.6
0.8
1.0

**Extended Data Fig. 23. Significant differentiated TE families in the nGer population.** Analyses of the 78 significantly differentiated TE families in nGer admixed individuals, showing observed and expected abundances.

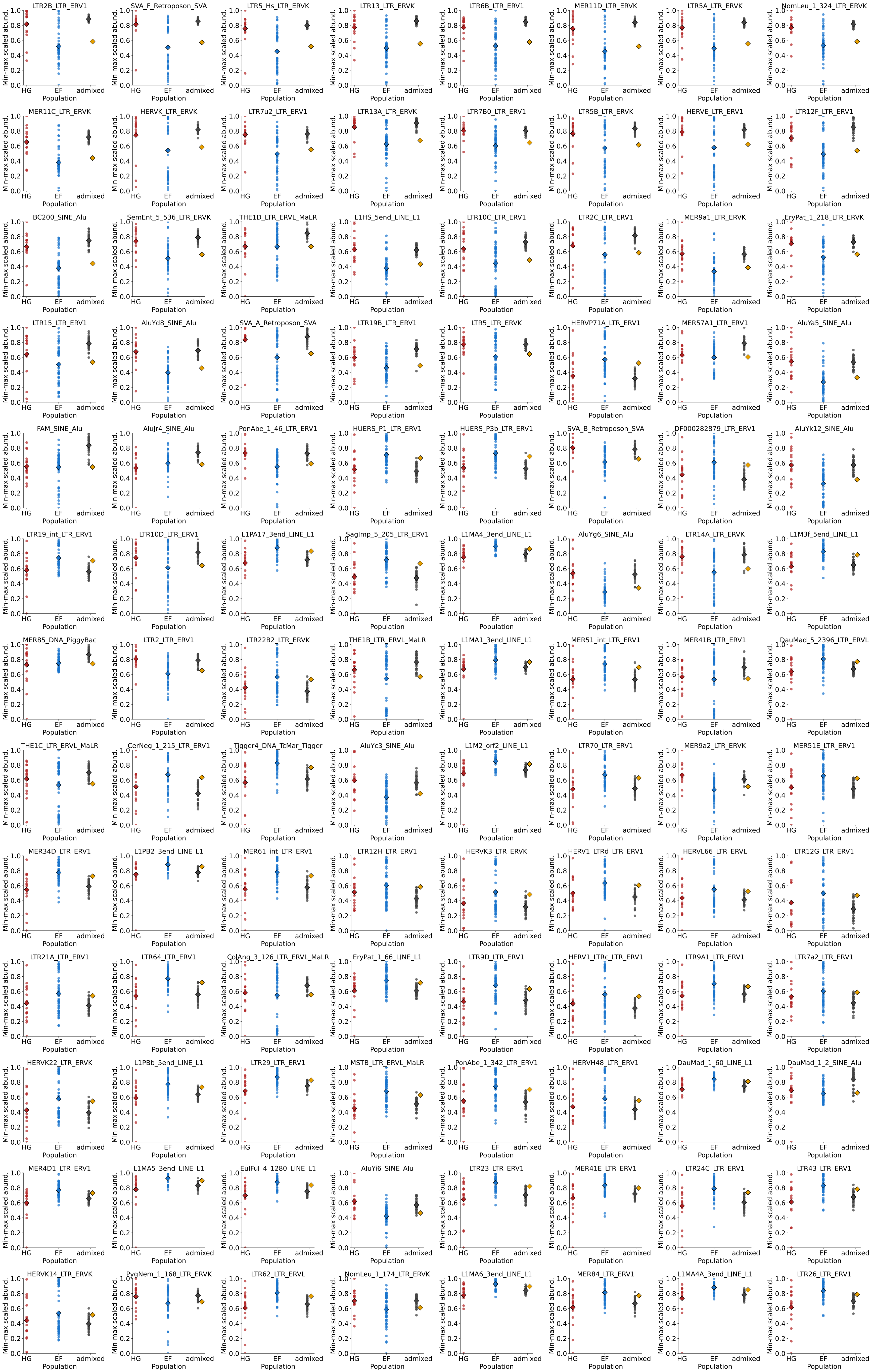
LTR2B_LTR_ERV1
SVA_F_Retroposon_SVA
LTR5_Hs_LTR_ERVK
LTR13_LTR_ERVK
LTR6B_LTR_ERV1
MER11D_LTR_ERVK
LTR5A_LTR_ERVK
NomLeu_1_324_LTR_ERVK
MER11C_LTR_ERVK
HERVK_LTR_ERVK
LTR7u2_LTR_ERV1
LTR13A_LTR_ERVK
LTR7B0_LTR_ERV1
LTR5B_LTR_ERVK
HERVE_LTR_ERV1
LTR12F_LTR_ERV1
BC200_SINE_Alu
SemEnt_5_536_LTR_ERVK
THE1D_LTR_ERVL_MaLR
L1HS_5end_LINE_L1
LTR10C_LTR_ERV1
LTR2C_LTR_ERV1
MER9a1_LTR_ERVK
EryPat_1_218_LTR_ERVK
LTR15_LTR_ERV1
AluYd8_SINE_Alu
SVA_A_Retroposon_SVA
LTR19B_LTR_ERV1
LTR5_LTR_ERVK
HERVP71A_LTR_ERV1
MER57A1_LTR_ERV1
AluYa5_SINE_Alu
FAM_SINE_Alu
AluJr4_SINE_Alu
PonAbe_1_46_LTR_ERV1
HUERS_P1_LTR_ERV1
HUERS_P3b_LTR_ERV1
SVA_B_Retroposon_SVA
DF000282879_LTR_ERV1
AluYk12_SINE_Alu
LTR19_int_LTR_ERV1
LTR10D_LTR_ERV1
L1PA17_3end_LINE_L1
SagImp_5_205_LTR_ERV1
L1MA4_3end_LINE_L1
AluYg6_SINE_Alu
LTR14A_LTR_ERVK
L1M3f_5end_LINE_L1
MER85_DNA_PiggyBac
LTR2_LTR_ERV1
LTR22B2_LTR_ERVK
THE1B_LTR_ERVL_MaLR
L1MA1_3end_LINE_L1
MER51_int_LTR_ERV1
MER41B_LTR_ERV1
DauMad_5_2396_LTR_ERVL
THE1C_LTR_ERVL_MaLR
CerNeg_1_215_LTR_ERV1
Tigger4_DNA_TcMar_Tigger
AluYc3_SINE_Alu
L1M2_orf2_LINE_L1
LTR70_LTR_ERV1
MER9a2_LTR_ERVK
MER51E_LTR_ERV1
MER34D_LTR_ERV1
L1PB2_3end_LINE_L1
MER61_int_LTR_ERV1
LTR12H_LTR_ERV1
HERVK3_LTR_ERVK
HERV1_LTRd_LTR_ERV1
HERVL66_LTR_ERVL
LTR12G_LTR_ERV1
LTR21A_LTR_ERV1
LTR64_LTR_ERV1
ColAng_3_126_LTR_ERVL_MaLR
EryPat_1_66_LINE_L1
LTR9D_LTR_ERV1
HERV1_LTRc_LTR_ERV1
LTR9A1_LTR_ERV1
LTR7a2_LTR_ERV1
HERVK22_LTR_ERVK
L1PBb_5end_LINE_L1
LTR29_LTR_ERV1
MSTB_LTR_ERVL_MaLR
PonAbe_1_342_LTR_ERV1
HERVH48_LTR_ERV1
DauMad_1_60_LINE_L1
DauMad_1_2_SINE_Alu
MER4D1_LTR_ERV1
L1MA5_3end_LINE_L1
EulFul_4_1280_LINE_L1
AluYi6_SINE_Alu
LTR23_LTR_ERV1
MER41E_LTR_ERV1
LTR24C_LTR_ERV1
LTR43_LTR_ERV1
HERVK14_LTR_ERVK
PygNem_1_168_LTR_ERVK
LTR62_LTR_ERVL
NomLeu_1_174_LTR_ERVK
L1MA6_3end_LINE_L1
MER84_LTR_ERV1
L1MA4A_3end_LINE_L1
LTR26_LTR_ERV1
Min-max scaled abund.
Population
HG
EF
admixed

**Extended Data Fig. 24. Top 104 (out of 171) most significantly differentiated TE families in the nIre population.** Analyses of the top 104 (out of 171) most differentiated TE families in nIre admixed individuals, showing observed and expected abundances.
